# A cell atlas of human and mouse synovium from early and advanced stages of knee osteoarthritis: BHLHE40 regulates fibroblast activation

**DOI:** 10.1101/2025.03.05.641539

**Authors:** Kabriya Thavaratnam, Eric Gracey, Anusha Ratneswaran, Jason S. Rockel, Shabana Vohra, Chiara Pastrello, Flore Stappers, Guillaume Planckaert, Katrina Hueniken, Johana Garcia, Paramvir Kaur, Starlee S. Lively, Pratibha Potla, Carolien Vlieghe, Y. Raja Rampersaud, Nizar N. Mahomed, Jan Victor, Nele Arnout, Igor Jurisica, Rajiv Gandhi, Dirk Elewaut, Mohit Kapoor

## Abstract

Osteoarthritis (OA) is a destructive joint disease affecting multiple tissues, including synovium. Previous studies have identified some distinct fibroblast subtypes within synovium; however, the characterization of fibroblast subsets during distinct stages of knee (K)OA disease, and their contributions to the endogenous mechanisms that drive synovial fibrosis during KOA, are not well characterized. Here we profile synovium from early- (KL I) and advanced- (KL III/IV) stages of radiographic KOA. First, bulk-RNA sequencing of early- and advanced-staged KOA synovial tissue revealed transcriptomic differences between the two disease stages. Using single-nuclei RNA sequencing (snRNA-seq) and flow cytometry, we identified distinct fibroblast subsets and uncovered an endotypic shift in fibroblast subsets during KOA pathogenesis, transitioning from DPP4+ in early-stage to ITGB8+ in advanced-stages. SnRNA-seq of synovium from mice with experimental KOA revealed analogous populations of Dpp4+ and Itgb8+ fibroblasts in tissue from early and advanced model stages. Human advanced-stage KOA synovial tissue had stronger expression of matrisome-annotated genes compared to early-stage tissue. BHLHE40, a crucial transcriptional regulator of ECM related genes, was identified as upregulated in ITGB8+ fibroblasts compared to DPP4+ fibroblasts. Using primary human OA fibroblasts in vitro, and conditional knock out mice in vivo, we found that fibroblast-intrinsic loss of *BHLHE40* increased fibrosis-related gene expression, enhanced fibroblast activation and induced severe synovial fibrosis in vivo. In contrast, overexpression of BHLHE40 in vitro was able to suppress TGF-β-induced fibroblast activation. Overall, this study provides a comprehensive cellular atlas of KOA synovium and has identified BHLHE40 as a crucial regulator of fibroblast-mediated synovial fibrosis.

## INTRODUCTION

Osteoarthritis (OA) is a complex and multifactorial joint disease affecting over 500 million people globally (*1*). As the most prevalent form of arthritis, OA significantly impacts the quality of life of afflicted individuals and imposes a substantial socio-economic healthcare burden (*2, 3*). The knee is the most common joint afflicted by OA, with patients experiencing chronic joint pain and stiffness, culminating in a progressive decline in mobility (*4, 5*). To measure the severity of knee (K)OA joint degeneration, radiographs are graded using the Kellgren-Lawrence (KL) system, with higher grades associated with more advanced disease (*6*). The field has acknowledged the importance of pre-radiographic, or “early OA”, as a critical timepoint in which effective disease modifying therapeutics/treatments could be administered (*7*). In this study, we refer to KL I as early-stage and KL III/IV as advanced-stage radiographic KOA.

Initially, OA was believed to be a disease of the articular cartilage; however, it is now acknowledged as a disease of the whole joint affecting multiple tissues (*8*). OA pathogenesis involves degradation of the articular cartilage, subchondral bone remodeling, and synovial fibrosis (*9*). Extracellular matrix (ECM)-rich synovium lines the joint capsule and functions to support joint health through the production of synovial fluid (required for lubrication), transport of nutrients into the joint, and removal of debris and waste (*10*). The synovium is made up of lining and sub-lining layers that are comprised of several different cell types including fibroblasts, macrophages, lymphocytes, and mast cells, among others (*11*). The most abundant cell type found in normal synovium is the fibroblast-like synoviocyte (referred to as fibroblasts throughout this manuscript). These cells secrete lubricating molecules, provide plasma-derived nutrients for joint tissues, and produce ECM components to maintain tissue architecture (*12-14*).

OA synovium presents with severe structural changes including inflammation, hyperplasia of the lining layer, and increased ECM deposition (fibrosis) (*15, 16*). These histological changes are reflected by cellular and molecular changes evident even before cartilage degeneration becomes visible, which include fibroblast proliferation, angiogenesis, mononuclear cell infiltration, and production of pro-inflammatory mediators (*17*). Although some studies have investigated fibroblast populations associated with lining and sublining layers in advanced stages of OA (*18, 19*), fibroblast subtypes in early and advanced stages of KOA, and their contributions to synovial fibrosis, are not well characterized.

To examine synovial fibroblasts in KOA, we took a step-wise approach using bulk RNA sequencing, single-nuclei RNA sequencing (snRNA-seq), flow cytometry, and advanced bioinformatic analyses, coupled with transgenic and surgical OA mouse models. We first created a catalogue of cells in the synovium, focusing on distinct fibroblast sub-populations present in early and advanced stages of radiographic KOA. We showcased an endotypic shift of fibroblast subset populations over disease stages and uncovered the contribution of the transcriptional regulator *BHLHE40* to fibroblast activation, ECM regulation and synovial fibrosis during KOA. To the best of our knowledge, this study is the first to identify stage-specific fibroblast subsets and describe *BHLHE40* as a crucial mediator involved in synovial fibrosis during KOA.

## RESULTS

### Distinct transcriptomic profiles of early- and advanced-stage KOA synovium

To begin exploring synovial pathologies during early and advanced stages of KOA, we first acquired synovial tissue samples from patients with KL I radiographic KOA (early-stage, undergoing arthroscopic surgery) and from patients with KL III/IV radiographic KOA [advanced-stage, undergoing total knee arthroplasty (TKA)] from a single center (Schroeder Arthritis Institute, Toronto Western Hospital, University Health Network, Toronto, ON, Canada). Harvested tissues were processed for histological investigation, or flash frozen for subsequent sequencing studies (Fig. 1A; table S1). Discernible structural differences were observed in the synovium of early and advanced stages of radiographic KOA using the Masson’s Trichrome staining. Compared to early-stage synovia, enhanced deposition of ECM and thickening of the synovial lining layer in advanced-stage tissues was notable (Fig. 1B, fig. S1A).

**Fig. 1.**
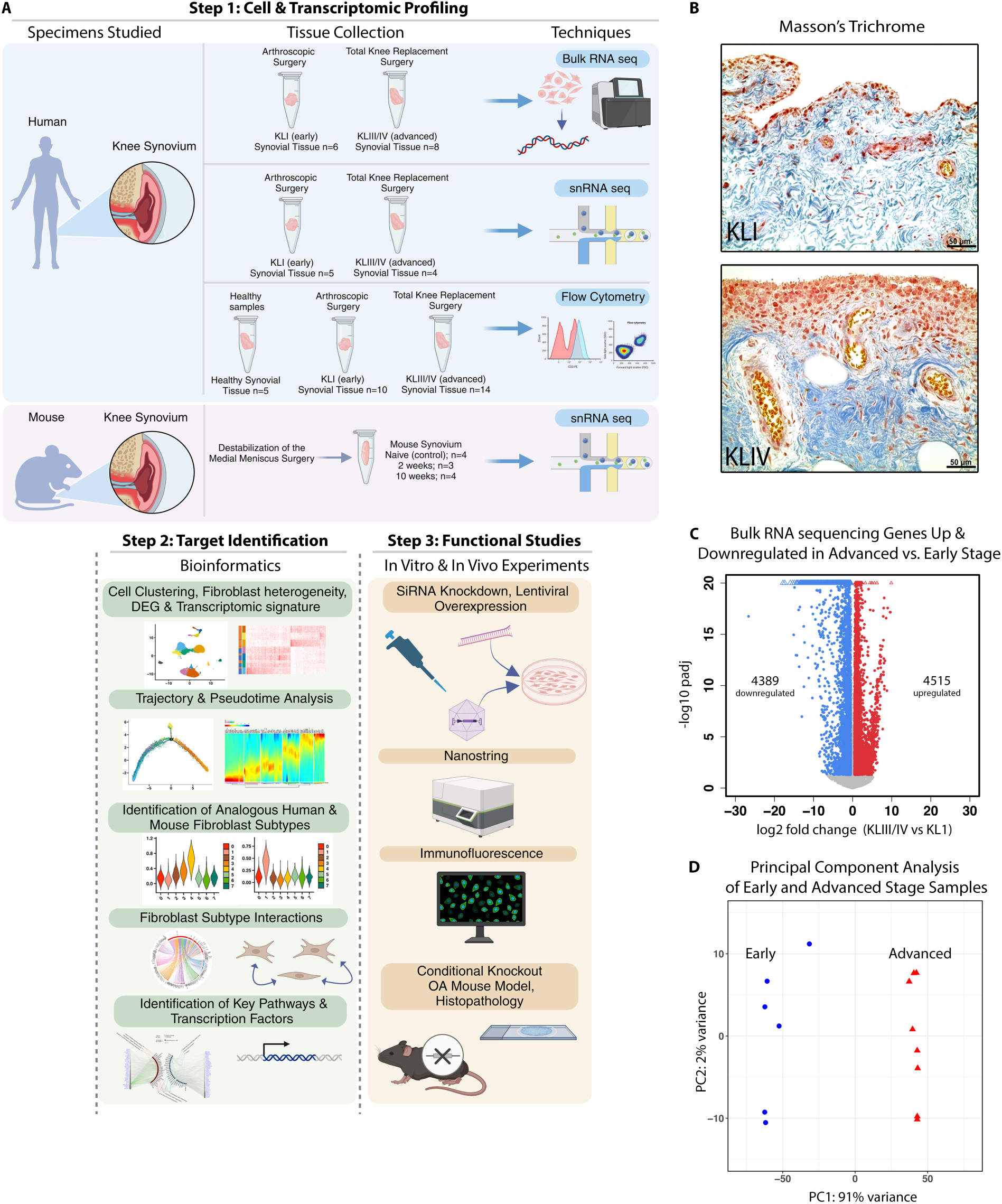
Structural and transcriptomic shifts in synovia correlate with knee OA progression. A) Schematic outlining the workflow of human and mouse experiments to identify cellular populations of synovial tissues. B) Masson’s Trichrome staining of KL I (left) and KL IV (right) graded radiographic knee OA synovia. C) Volcano plot showing the log2 fold change (log2FC) of differentially-expressed genes (DEGs) in advanced stage samples compared to early samples. Genes with a log2FC > 0.5, adjusted p-value <0.05 are upregulated while genes with a log2FC < -0.5, adjusted p-value <0.05 are downregulated. D) Principal component analysis of gene expression profiles of early- and advanced-stage samples from bulk RNA sequencing. Figure was created, in part, using Biorender.

To evaluate transcriptomic differences between early- and advanced-stage synovia, we subjected synovial tissue from early (KL I; n=6) and advanced stages (KL III/IV; n=8) of radiographic KOA to bulk RNA sequencing, identifying a total of 4,515 upregulated genes and 4,389 downregulated genes in advanced-compared to early-stage tissues (Fig. 1C, table S2). Principal component analysis confirmed separation of samples based on disease stage, with PC1 explaining 91% of the variance (Fig. 1D). These findings prompted us to perform a more detailed investigation to determine which cell types may be driving synovial transcriptomic profiles during early and advanced stages of KOA.

### Cellular composition of early- and advanced-stage KOA synovium

We next subjected nine KOA synovium samples (five KL I and four KL III/IV) to snRNA-seq. After quality control and filtering, 25,285 nuclei were analyzed. Unsupervised clustering identified nine distinct cell types using canonical cell type markers (Fig. 2A, B, table S3). Identified cell types, listed from most to least abundant, were fibroblasts, macrophages, endothelial cells, adipocytes, lymphocytes, mural cells, dendritic cells, proliferating cells and mast cells. While the proportions of cell types identified did have some variation between patients, the largest shift in proportion was observed between disease stages (Fig. 2A). In all subjects, fibroblasts were the predominant cell type identified, comprising approximately 50% of the sequenced nuclei in each synovial tissue sample.

**Fig. 2.**
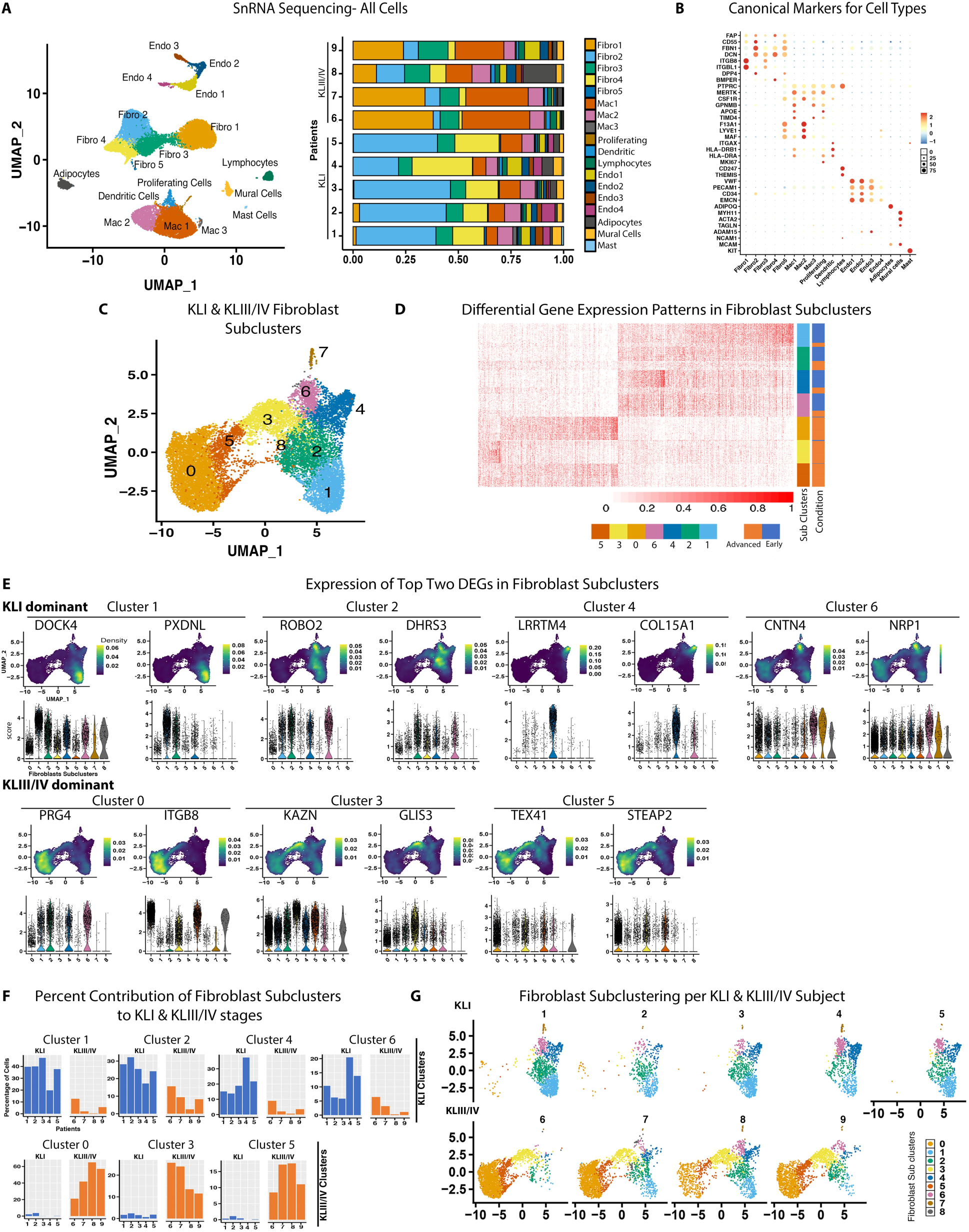
SnRNA-seq reveals major cell types and fibroblast subtypes in early- and advanced-stage knee OA synovia. A) Uniform manifold approximation and projection (UMAP) of all clusters identified in human synovia (left) through snRNA-seq. Proportion of cell types found in each sample (n=9) shown as a stacked bar graph (right). B) Dot plot showing the expression of marker genes used to annotate the clusters from KOA human synovia. C) All fibroblast subclusters identified in KL I and KL III/IV graded radiographic knee OA synovia represented as a UMAP. D) Heatmap displaying gene expression differences between KL I- and KL III/IV-dominant fibroblast subclusters. E) Density and violin plots of top two DEGs from fibroblast subclusters. F) Bar graph showing the percent contribution of all fibroblast subclusters to KLI and KLIII/IV stages. G) Individual patient UMAPs of fibroblast subclusters.

### Nine transcriptomically distinct fibroblast subpopulations identified in human KOA synovium

Since fibroblasts were the predominant cell type identified in human KOA synovium, our subsequent analyses focused on this cell type. Unbiased clustering analysis was performed on nuclei determined to be fibroblasts, revealing nine transcriptionally distinct fibroblast subtypes based on differentially-expressed gene (DEG) profiles (Fig. 2C&D). These subtypes were defined by unique DEG lists consisting of a maximum of 235 genes (subcluster 0), to a minimum of 3 genes (subcluster 5). Of note, subcluster 8 had 0 DEGs. The top 5 genes associated with fibroblast subclusters 0 to 7, based on decreasing Log2FC (q<0.05, Log2FC≥0.5, min.pct≥0.25) included subcluster 0: *PRG4*, *ITGB8, SEMA5A, CRTAC1, CLIC5*; subcluster 1: *DOCK4*, *PXDNL, KCNB2, FHOD3, KANK1*; subcluster 2: *ROBO2, DHRS3, AFF3, HMCN1, FBLN1*; subcluster 3: *KAZN, GLIS3, PTGFR, CXCL12, RUNX1*; subcluster 4: *LRRTM4, COL15A1, LAMA2, ABCA10, BMPER*; subcluster 5: *TEX41, STEAP2, EFNA5*; subcluster 6: *CNTN4, NRP1, PAK3, ELN, MBNL1;* subcluster 7: *MTSS1, TRIM22, SYNPO2, KLF7, MGLL* (Fig. 2E, fig. S1B & S2).

We next compared our identified fibroblast subclusters to characterizations from previously published studies. Fibroblasts from advanced-stage OA synovium have been categorized into two distinct populations: synovial sublining fibroblasts (SSF) and synovial intimal fibroblasts (SIF) (*20*). In our study, cluster 0 fibroblasts closely resembled the SIF (lining) population, while cluster 1 fibroblasts were more similar to the SSF (sublining) population (fig. S3A). Compared to the AMP-RA dataset of rheumatoid arthritis and late stage OA fibroblasts (*21*), early-stage OA fibroblasts from our study (clusters 1, 2, 4, and 6) resembled that study’s identified SC-F1 (CD34+) population, while advanced-stage OA fibroblasts from our study (clusters 0, 3, and 5) resembled the SC-F4 (CD55+) population (fig. S3B). These comparisons indicate that the fibroblasts identified in our study align with those from previously published studies.

### Endotypic shift in synovial fibroblasts from early to advanced KOA

Interestingly, when we investigated whether there was a disproportionate contribution of nuclei from early- or advanced-stage tissues to the identified fibroblast clusters, we found that early-stage tissues contributed proportionally more nuclei to clusters 1, 2, 4 and 6, while advanced-stage tissues contributed proportionally more nuclei to clusters 0, 3 and 5 (Fig. 2F, G, fig. S1C). Subclusters 7 and 8 showed no significant differences in proportion of nuclei contributed by early or advanced-stage tissues. Subcluster 0, with the highest proportion of nuclei derived from advanced-stage KOA synovial tissues, was the predominant fibroblast subtype in advanced-stage tissue. In contrast, subcluster 1, with the highest proportion of nuclei contributed from early-stage tissues, was the predominant fibroblast subtype from early-stage KOA synovial tissues.

We next used trajectory analysis to determine if fibroblasts from early-stage tissue had the potential to differentiate into those found in advanced-stage tissue. Trajectory analysis suggested that advanced-stage fibroblast subtypes (clusters 0, 3, and 5) may be derived from early-stage fibroblast populations through intermediate populations via progressive changes in gene expression (Fig. 3A&B, fig. S4A). These data suggest an endophenotypic shift in fibroblast subtypes in synovium from early to advanced stages of KOA. Of note, key genes contributing to the determined trajectory included *DOCK4*, *PXDNL*, *ROBO2*, *DHRS3*, *LRRTM4*, *COL15A1* and *CNTN4,* which were also highly expressed in early-stage predominant clusters 1, 2, 4, and 6, and *PRG4, ITGB8, KAZN, GLIS3* and *TEX41*, which were also highly expressed in advanced-stage predominant clusters 0, 3, and 5 (Fig. 3C, fig. S4B).

**Fig. 3.**
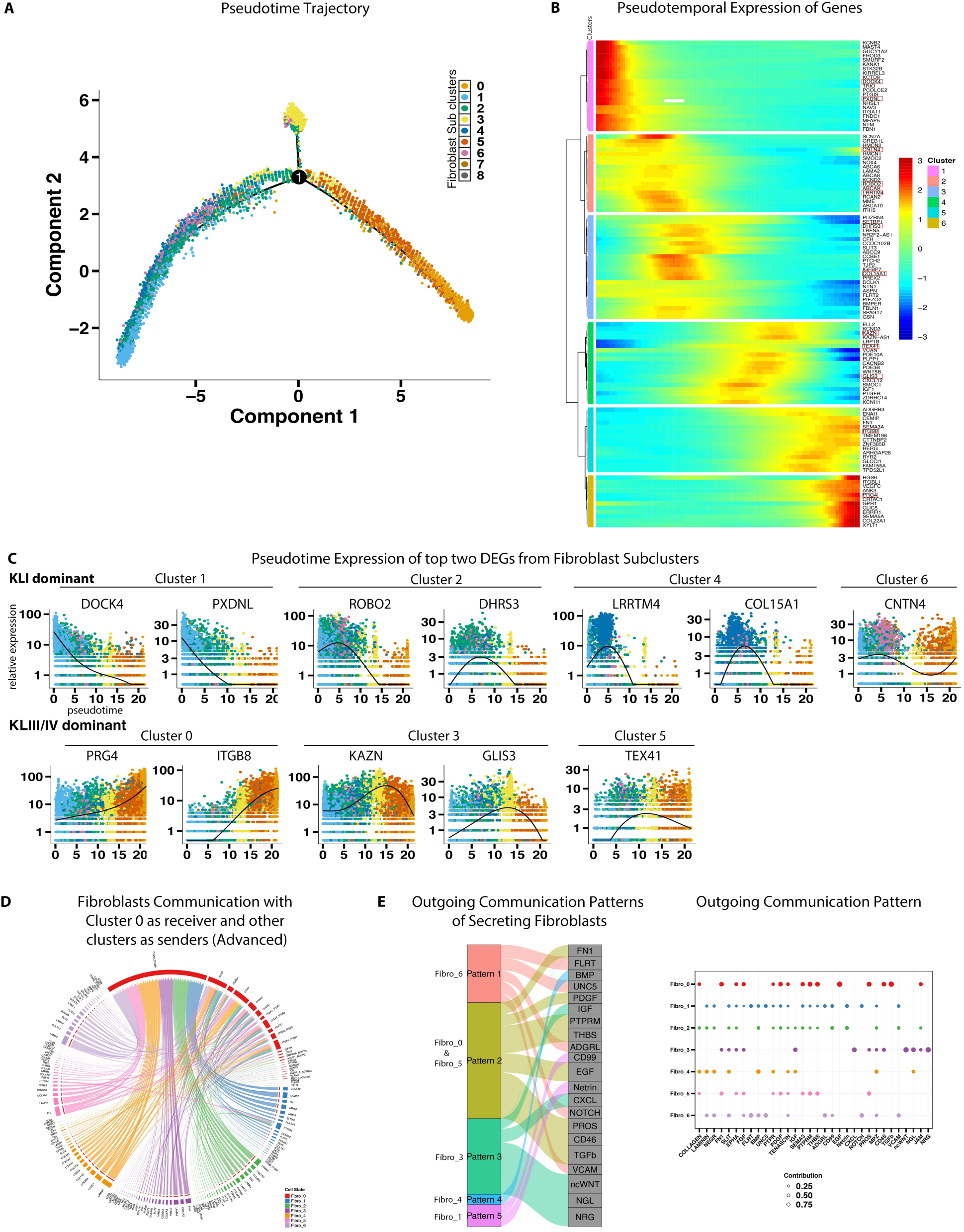
Advanced-stage fibroblasts may arise from early-stage fibroblasts and have unique communication patterns. A) Trajectory analysis showing changes in gene expression in fibroblast subclusters across pseudotime. B) Pseudotemporal changes in gene expression patterns across fibroblast subclusters. C) Jitter plots showing gene expression changes of top two DEGs across pseudotime in fibroblast subclusters. D) Chord diagram illustrating putative cell-cell signaling interactions between fibroblast cluster 0 expressing receptors and all other major fibroblast subclusters expressing ligands. E) River plot showing outgoing communication pattern of secreting cells where outgoing pattens reveal how the sender cells coordinate with each other as well as how they coordinate with certain signaling pathways to drive communication (left). Dot plot showing contribution of outgoing communication of pathways in different fibroblast sub-clusters (right).

We further investigated putative cell-cell interaction patterns between major cell types and subtypes using CellChat v2.0 (*22*). Comparing the top four major cell types (fibroblasts, macrophages, endothelial cells, adipocytes), signaling within fibroblasts exhibited the highest interaction strength, prompting us to further investigate interactions among fibroblast subtypes (fig. S5A, B). Within fibroblast subtypes, the strongest interaction strengths were observed directed towards fibroblast cluster 0, with the second strongest interaction strengths directed towards fibroblast cluster 1 (fig. S5C). To further define cell-cell communication between fibroblast cluster 0 and other subtypes, we investigated ligand-receptor interactions with fibroblast cluster 0 set as the “receiver,” expressing receptors, and all other fibroblast subtypes set as “senders,” expressing ligands (Fig. 3D, fig. S5D, E). A number of communication patterns to fibroblast cluster 0 from other fibroblast subtypes were identified, with the highest proportion of signals being received by fibroblast subcluster 0. We also found that the various fibroblast subtypes had multiple signaling patterns composed of various pathways. Interestingly fibroblast subtypes 0 and 5 both followed pattern 2, which was associated with TGF-β, FN1, PDGF and other outgoing communication pathways. In contrast, fibroblast subtypes 1, 3, 4, and 6 were associated with distinct communication patterns linked to various unique outgoing communication pathways including NOTCH, VCAM, and non-canonical (nc)WNT. (Fig. 3E, fig. S5F). Putative ligand-receptor interactions between fibroblast subsets (senders) and cluster 1 (receiver) are detailed in Fig S5G.

Overall, fibroblast subcluster 0 was found to be predominant in the advanced stages of the disease while subcluster 1 was predominant in the synovium of early-stage KOA with subcluster 0 having the strongest interaction strength. Furthermore, we determined that synovial fibroblasts likely undergo an endotypic shift from early to advanced stages of KOA.

### Flow cytometry confirms distinct fibroblast sub-populations in KOA

To independently confirm the presence of major fibroblast subclusters observed by snRNA-seq, an independent cohort of early- and advanced-stage KOA patients was generated at a second center (Ghent, Belgium). This cohort also included anatomically matched healthy synovia obtained from post-mortem tissue donors with no history of musculoskeletal disease. Suprapatellar pouch synovial tissues were formalin fixed for histology and cryopreserved for flow cytometry (Fig. 4A). Cryopreserved synovial tissue was thawed and enzymatically digested. Consistent with previous reports of synovial hypertrophy in OA (*1*), a progressive increase in cell yield was observed from healthy tissues through to advanced-stage KOA synovia (fig. S6A).

**Fig. 4.**
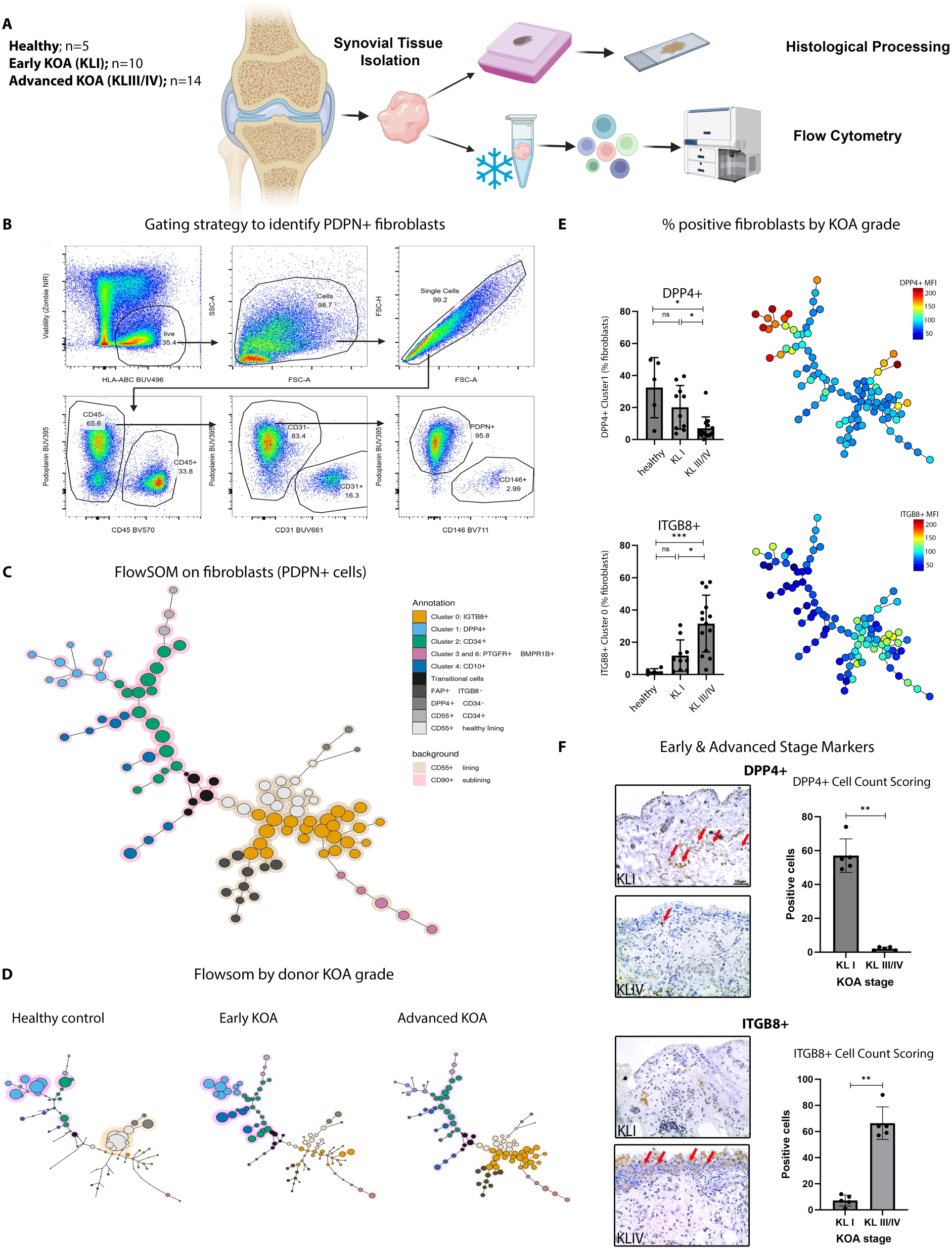
DPP4 and ITGB8 define early- and advanced stage fibroblast subsets. An independent cohort was generated to explore synovial fibroblast endotype transition with KOA progression. A) Schematic outlining the workflow of human synovial tissues used in flow cytometry and histology experiments. B) Gating strategy used to identify PDPN+ fibroblasts. C) FloSOM illustration of PDPN+ fibroblast subclusters, segregated by lining (CD55+) and sublining (CD90+). D) FloSOM illustration showing separate healthy control, early KOA and advanced KOA data. E) Percent positive ITGB8+ and DPP4+ fibroblasts depicted by KOA grade. Multiple unpaired t-tests were performed and corrected for multiple testing. F) IHC staining of both DPP4+ and ITGB8+ cells (examples indicated by red arrows) in KL I- and KL III/IV-graded radiographic KOA human synovial tissue and positive cell count scoring. Positive cell counts were statistically analyzed using an unpaired, nonparametric, Mann Whitney test. Data in (E) and (F) are presented as mean ± standard deviation. Not significant (n.s.), *, P ≤ 0.05, **, P ≤ 0.01, and ***, P ≤ 0.001. Figure was created, in part, using Biorender.

A 19-marker flow cytometry panel was assembled to examine subsets of fibroblasts identified by snRNA-seq. First, pan-MHCI (HLA-ABC) and a fixable viability marker were used to separate live cells from debris, following which, lineage markers were used to gate out immune cells (CD45+), endothelial cells (CD31+) and mural cells (CD146+) before using PDPN to identify fibroblasts (Fig. 4B). As expected, immune cells and fibroblasts were the dominant cell types identified in synovia (fig S6B). Of note, we found a moderate increase in immune cells with progressive KOA; noteworthy given that OA is often used as “non-inflammatory” control tissue for inflammatory arthritis studies (*21*).

A clear shift in broad KOA fibroblast populations was observed when using conventional markers for subsets and activation (fig S6C-E). CD90 and CD55 are commonly used to identify sublining and lining fibroblasts respectively (*23*). Using these markers, we found an increase in the frequency of lining fibroblasts in advanced-compared to early-stage KOA synovia, consistent with lining layer hyperplasia formation observed by histology (Fig. 1B) (*1*). Further, fibroblast activation protein (FAP) and CD63, additional fibroblast activation markers, were elevated with KOA compared to healthy controls, but were only moderately increased with stage of KOA. FlowSOM was next used to unbiasedly cluster fibroblasts in our flow cytometry data using markers identified by snRNA-seq (Fig. 4C). By examining the intensity of markers on each cluster, we identified 11 metaclusters. Many of the metaclusters appeared to match those in our snRNA-seq analyses based on expression profiles of key markers including DPP4, CD34, ITGB8 and PTGFR (fig. S6F). Two notable metaclusters included DPP4+CD34+ fibroblasts, which endotypically resembled snRNA-seq subcluster 1 fibroblasts, and ITGB8+ fibroblasts, which resembled snRNA-seq subcluster 0 fibroblasts. Thus, subcluster 0, the major predominant fibroblast population in advanced-stage KOA synovia, will now be referred to as ITGB8+ fibroblasts, and subcluster 1, the predominant fibroblast population in early-stage KOA synovial tissues, will now be referred to as DPP4+ fibroblasts.

We found a gradient in the distribution of fibroblasts across metaclusters from healthy to early- and advanced-stage KOA, with ITGB8+CD55+ fibroblast clusters coming primarily from advanced-stage KOA synovia (Fig. 4D). Examination of these clusters on a per subject basis revealed a significant reduction in the frequency of DPP4+ fibroblasts and an increase in ITGB8+ fibroblasts with KOA progression (Fig. 4E, fig. S6G). Immunohistochemistry (IHC) further showed that ITGB8+ cells were more frequent in advanced-stage tissue (Fig. 4F). Conversely, DPP4+ cells were more frequent in early-stage tissues compared to advanced-stage tissue.

Overall, this data confirms the endotypic shift in fibroblasts from early- to advanced-stages of the disease and demonstrates subtler phenotypic changes from healthy to early disease stage. Consistent with the findings from our snRNA-seq data, flow cytometry further validated DPP4+ fibroblasts as the major fibroblast subcluster found in the synovium during early-stage KOA whereas ITGB8+ fibroblasts are the major fibroblast subcluster found in synovium of advanced-stage KOA.

### Itgb8+ synovial fibroblasts expand during KOA in a mouse model

Having observed the same fibroblast endotypes with different techniques in two independent human OA cohorts, we next sought to determine if similar early- and advanced-stage predominant fibroblast populations were present in murine KOA. Destabilization of the medial meniscus (DMM) surgery was performed in mice and synovia was collected at two (n=3) or ten weeks (n=4) post-surgery. Naïve mouse synovia (non-surgical) was collected as a control (n=4).

The collected tissue was subjected to snRNA-seq (Fig. 5A). A total of 19,304 nuclei were analyzed after filtering and subjected to unsupervised clustering, which resolved a total of ten distinct cell types utilizing canonical cell surface markers (fig. S7A-C). Further clustering analysis of fibroblasts identified eight transcriptionally distinct fibroblast subclusters (Fig. 5B, fig. S7D, table S4).

**Fig. 5.**
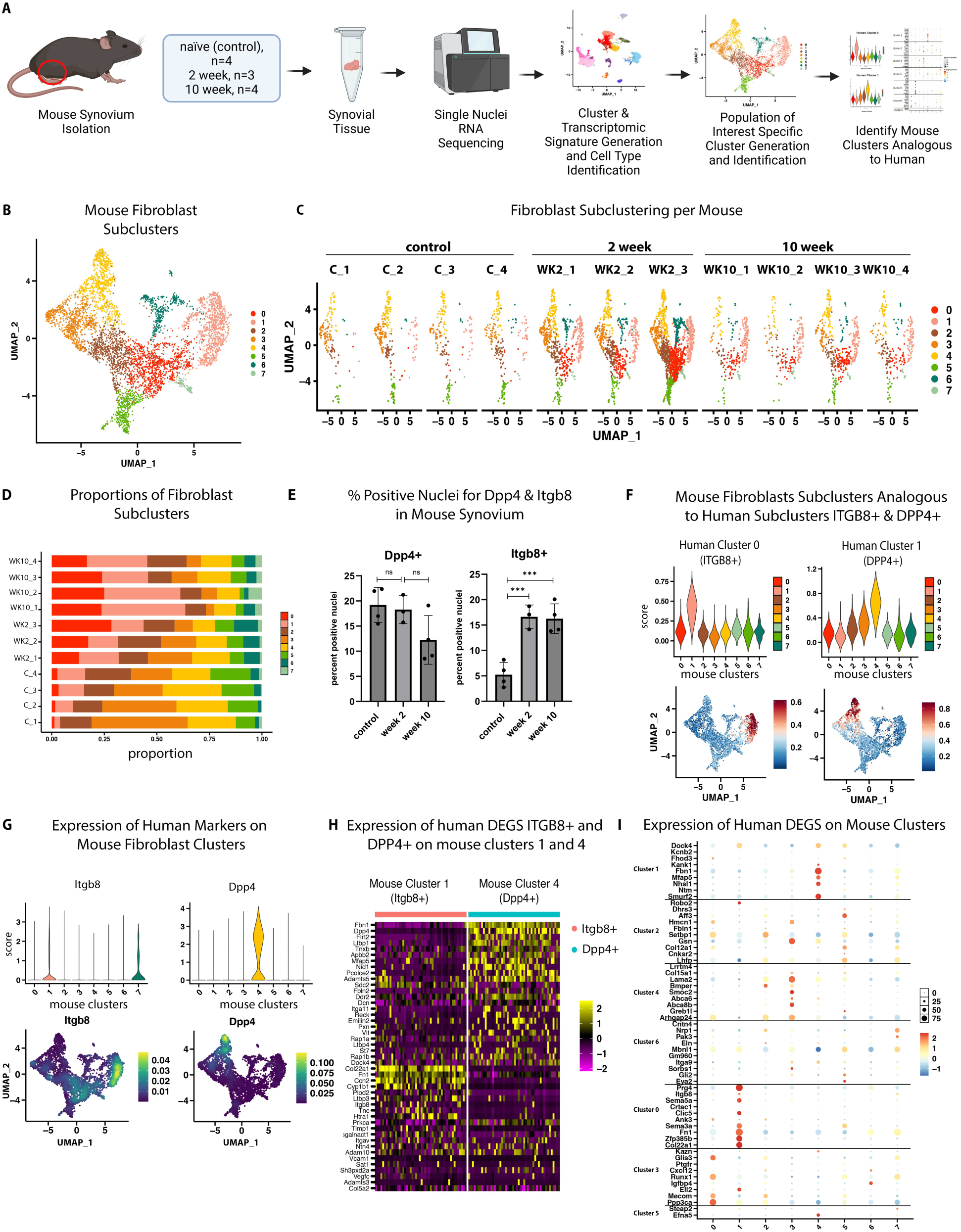
Itgb8+ synovial fibroblasts emerge during surgically-induced mouse KOA. A) Schematic of mouse synovial tissue isolation and snRNA sequencing. B) Mouse synovial fibroblast subclusters identified in naïve (non-surgical control; n=4), 2-week (n=3) and 10-week (n=4) post-DMM surgery visualized by UMAP. C) Individual fibroblast UMAPs of each mouse synovial sample analyzed. D) Stacked bar graph illustrating proportions of mouse fibroblast subclusters. E) Percent positive nuclei for Dpp4 and Itgb8 in mouse synovia in control, 2-week and 10-week conditions. Data presented as mean ± standard deviation. Data was log-transformed and analyzed by one-way ANOVA with post-hoc tests corrected for multiple comparisons usingthe two-stage step-up method of Benjamini, Krieger, and Yekutieli. Not significant (n.s.), *, P ≤ 0.05, **, P ≤ 0.01, and ***, P ≤ 0.001. F) Violin plots and density plots showing mouse clusters (1 and 4) analogous to human clusters 0 (ITGB8+) & 1(DPP4+) based on genes signature. G) Expression of ITGB8 and DPP4, on mouse fibroblast clusters. H) Expression heatmap of top human DEGs from ITGB8+ and DPP4+ fibroblasts on mouse clusters 1 and 4. I) Expression dot plot of top human DEGs from each fibroblast subcluster onto mouse fibroblast clusters. Figure was created, in part, using Biorender.

A change in the proportion of the fibroblast subtypes was identified as the model progressed from early (two weeks) to advanced stages (10 weeks) (Fig. 5C, D). Higher proportions of Dpp4+ nuclei were present in synovia from control and week 2 post-surgery mice, whilst higher proportions of Itgb8+ nuclei were found in tissues from both week 2 and week 10 post-surgical mice compared to controls (Fig. 5E). These findings emphasize that transcriptomically distinct fibroblast subsets are also present at various stages of murine KOA, akin to our findings in synovia from human KOA.

We next sought to identify if analogous mouse fibroblast subtypes resembled human fibroblast subtypes identified by snRNA-seq analysis of human synovia. Gene signatures from human fibroblast subpopulations were assessed in each mouse fibroblast subtype, illustrating that human ITGB8+ fibroblasts (cluster 0) were most similar to mouse cluster 1 fibroblasts, while human DPP4+ fibroblasts (cluster 1) were most similar to mouse cluster 4 fibroblasts (Fig. 5F, fig. S7E). *Itgb8* exhibited highest expression in mouse fibroblast cluster 1, with *Col22a1*, *Fn1*, and *Ccn2*, also highly expressed in both mouse fibroblast cluster 1 and the human ITGB8+ population (cluster 0). In contrast, *Dpp4* was predominantly expressed in mouse fibroblast cluster 4, with *Fbn1*, *Flrt2* and *Ltbp1* also highly expressed in both mouse fibroblast cluster 4 and the human DPP4+ fibroblast population (cluster 1) (Fig. 5G&H). Additionally, the expression of top DEGs from each human fibroblast subcluster in mouse subclusters confirmed similarities between human ITGB8+ and DPP4+ fibroblasts and mouse fibroblast subclusters 1 and 4, respectively (Fig. 5I). These findings support similarities between key human and mouse fibroblast subtypes during progressive stages of KOA.

### DPP4+ and ITGB8+ fibroblasts are associated with ECM-related pathways

Following investigations of fibroblast subpopulations in two independent human cohorts, and in a KOA mouse model, we next investigated putative functional roles of major human fibroblast subclusters. Pathway enrichment analysis was performed on DEGs from human fibroblast subclusters 0-6 using gene ontology and pathDIP v4 (*24*). Comprehensive pathway analysis identified many pathways enriched in each fibroblast subset (fig. S8). Interestingly both predominant human fibroblast subpopulations from advanced and early-stage synovia (ITGB8+ and DPP4+, respectively) shared a common pathway including: extracellular matrix organization and GO biological processes including: extracellular matrix organization, extracellular structure organization, axon guidance, axonogenesis, neuron projection guidance, positive regulation of neuron projection development and regulation of cell morphogenesis (fig. S8). It is established that excessive fibroblast proliferation and ECM deposition can contribute to the development of synovial fibrosis (*25, 26*). Given that ITGB8+ and DPP4+ fibroblasts were both associated with ECM-related pathways, we directed our subsequent studies to investigate these two major subpopulations.

### Identification of BHLHE40 as an upstream transcriptional regulator linked to ECM pathways

A total of 48 ECM-related DEGs from ITGB8+ (24 genes) and DPP4+ (24 genes) fibroblast populations were enriched with ECM-related pathways (Fig. 6A&B). These genes were then used to search Catrin (*27*) to identify putative upstream transcription factors regulating these genes (Fig. 6A, table S5). Catrin is an integrated database of transcription regulatory networks, linking transcription factors (TFs) to target genes (https://ophid.utoronto.ca/Catrin). Using this analysis, we identified that ECM pathway-related DEGs from ITGB8+ fibroblasts were putatively regulated by 31 TFs (table S6A), while ECM pathway-related DEGs from DPP4+ fibroblasts were putatively regulated by 39 TFs (table S6B).

**Fig. 6.**
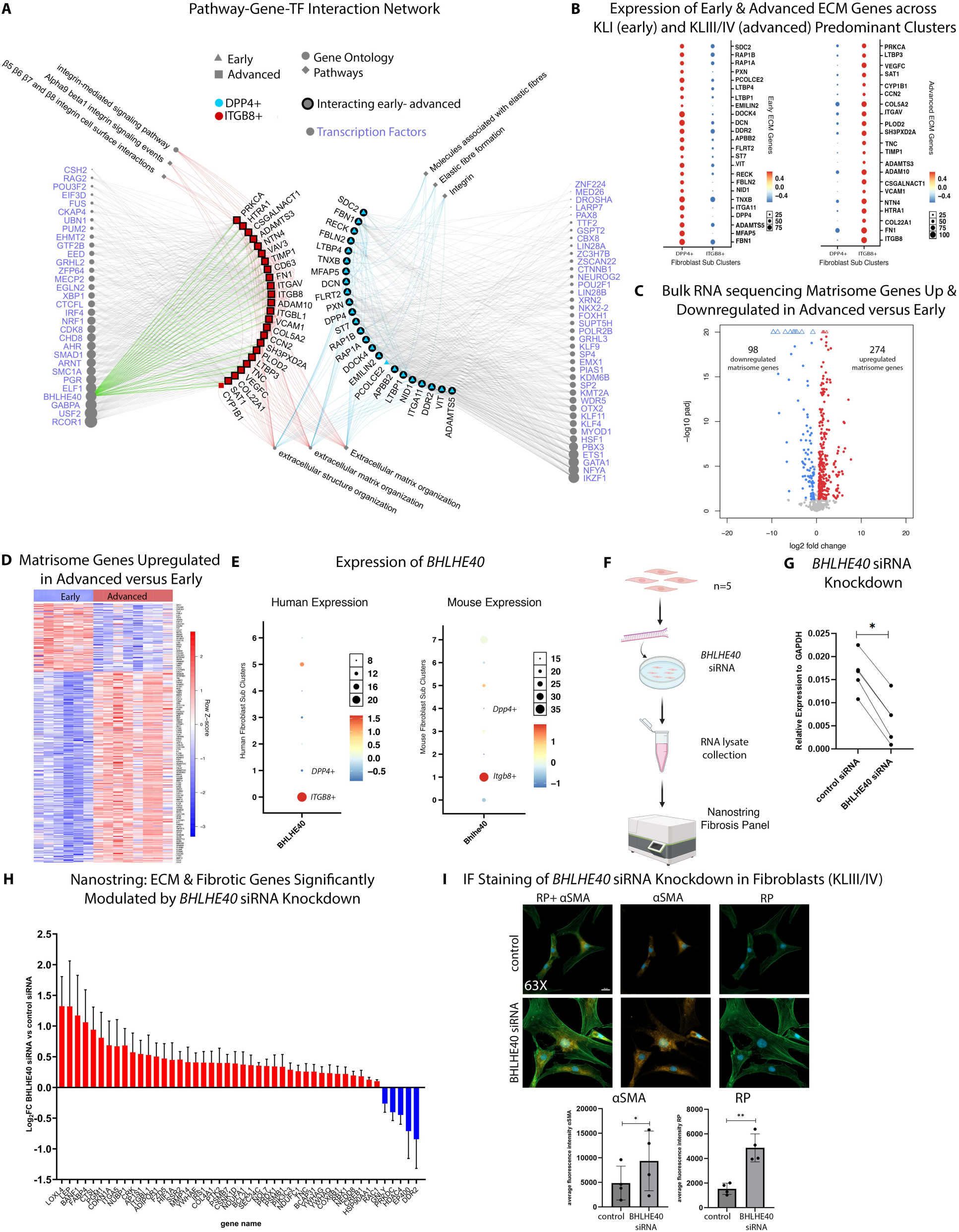
BHLHE40 is a key transcription factor regulating synovial fibroblast activation. A) Pathway-gene-TF interaction network showing putative transcription factors regulating DPP4+ and ITGB8+ fibroblast ECM genes and their enriched pathways. B) Expression of advanced and early ECM genes across KLI (early, DPP4+) and KLIII/IV (advanced, ITGB8+) predominant clusters. C) Volcano plot of matrisome-annotated genes up and downregulated in advanced-versus early-stage bulk RNA sequencing. Genes with log2FC > 0.5, adj. p-value < 0.05 are upregulated, while log2FC < -0.5, adj. p-value < 0.05 are downregulated. D) Heatmap illustrating matrisome genes upregulated in advanced-versus early-stage tissues. E) *BHLHE40/Bhlhe40* expression across fibroblast subclusters from human and mouse data. F) Schematic of in vitro BHLHE40 siRNA knockdown in advanced-stage fibroblasts. G) Relative gene expression of siRNA knockdown of *BHLHE40* in advanced-stage fibroblasts (n=5) in vitro. Data was analyzed by two-tailed paired T-tests. * P ≤ 0.05. H) Graph showing Log2FC of ECM and fibrotic genes significantly modulated by BHLHE40 siRNA knockdown. I) α-SMA and rhodamine phalloidin (RP) staining of *BHLHE40* siRNA-treated advanced-stage fibroblasts. Data presented as mean ± standard deviation. Data was log transformed and analyzed by two-tailed paired T-tests. *, P ≤ 0.05. Figure was created, in part, using Biorender.

To investigate differences in ECM-related transcriptomes in early- and advanced-stage KOA synovia, we also assessed DEGs in our bulk-RNAseq analysis for matrisome (ECM-associated genes) annotations. Out of a total of 1027 matrisome genes, 731 were collectively expressed in early and advanced stage KOA synovia. Of the 731 expressed genes, 50.69% (n=372) were differentially-expressed in advanced-versus early-stage KOA synovia, with 37.48% (n=274) upregulated in advanced-stage tissues and 13.41% (n=98) upregulated in early-stage tissues (Fig. 6C, & D, fig. S9A).

We next focused on the advanced-stage KOA synovia, particularly ITGB8+ fibroblasts, due to high ECM gene expression and evident synovial fibrosis observed in tissues from this disease stage. Further investigation of the ECM-regulatory transcription factors from our network analysis found that *BHLHE40* was the only TF that putatively regulated at least 80% (20/24) of the ECM gene list in ITGB8+ cells, and its expression was significantly higher between ITGB8+ and DPP4+ fibroblasts in humans (Fig. 6E, table S6A, S7). *BHLHE40* (DEC1/STRA13) is a broadly expressed transcription factor with repressive functions (*28*). It was initially identified for its minor role in circadian rhythms (*29*), and more recently was found to regulate immune cell function (*30, 31*). Since our data suggested that *BHLHE40* could be a crucial upstream transcriptional regulator linked to ECM pathways, this led us to further evaluate its role in fibroblast activation and synovial fibrosis.

### BHLHE40 negatively regulates ECM expression and fibroblast activation in vitro

To begin investigating the link between *BHLHE40* and the ECM transcriptome, fibroblast activation and fibrosis, we targeted *BHLHE40* in cultures of fibroblasts obtained from synovia of advanced-stage KOA. Using siRNA knockdown, we reduced *BHLHE40* expression by an average of 66.4% (KL III/IV; n=5) (Fig. 6F, G). RNA collected from *BHLHE40* siRNA- and control siRNA-treated cultures were subjected to analysis using a custom-made NanoString panel comprising of 777 fibrosis-related genes and ECM genes related to BHLHE40. Knockdown of *BHLHE40* in fibroblasts led to a notable increase in the expression of 41 fibrosis-related genes and a significant decrease in expression of only 5 genes (Fig. 6H, fig. S9B&C, table S8). Next, we investigated the effects of *BHLHE40* silencing on fibroblast activation. Activated fibroblasts express α-smooth muscle actin (αSMA), a marker of fibroblast activation (*32*), and induce stress fiber formation (*14*). Cultured fibroblasts with siRNA-targeted *BHLHE40* knockdown were stained for αSMA and rhodamine phalloidin. *BHLHE40* knockdown revealed fibroblast activation associated with enhanced stress fiber formation, cell spreading and increased αSMA expression (Fig. 6I). These data suggest that BHLHE40 may play a role in inhibiting fibroblast activation and ECM gene expression, consistent with its proposed function as a transcriptional repressor.

### BHLHE40 conditional knockout mice exhibit severe synovial fibrosis

To further investigate the influence of BHLHE40 on ECM regulation and synovial fibrosis in vivo, we generated a conditional knock out (CKO) mouse. Here, *Bhlhe40* was deleted by Cre-mediated recombination in cells expressing Col6a1 (*33*), which include synovial fibroblasts (Fig. 7A). CKO mouse knee fibroblasts showed a significant reduction in *Bhlhe40* gene expression compared to those from wild-type (WT) mice, without affecting expression in synovial macrophages (Fig. 7B), confirming selectivity of targeted deletion of Bhlhe40 in knee synovial fibroblasts in vivo.

**Fig. 7.**
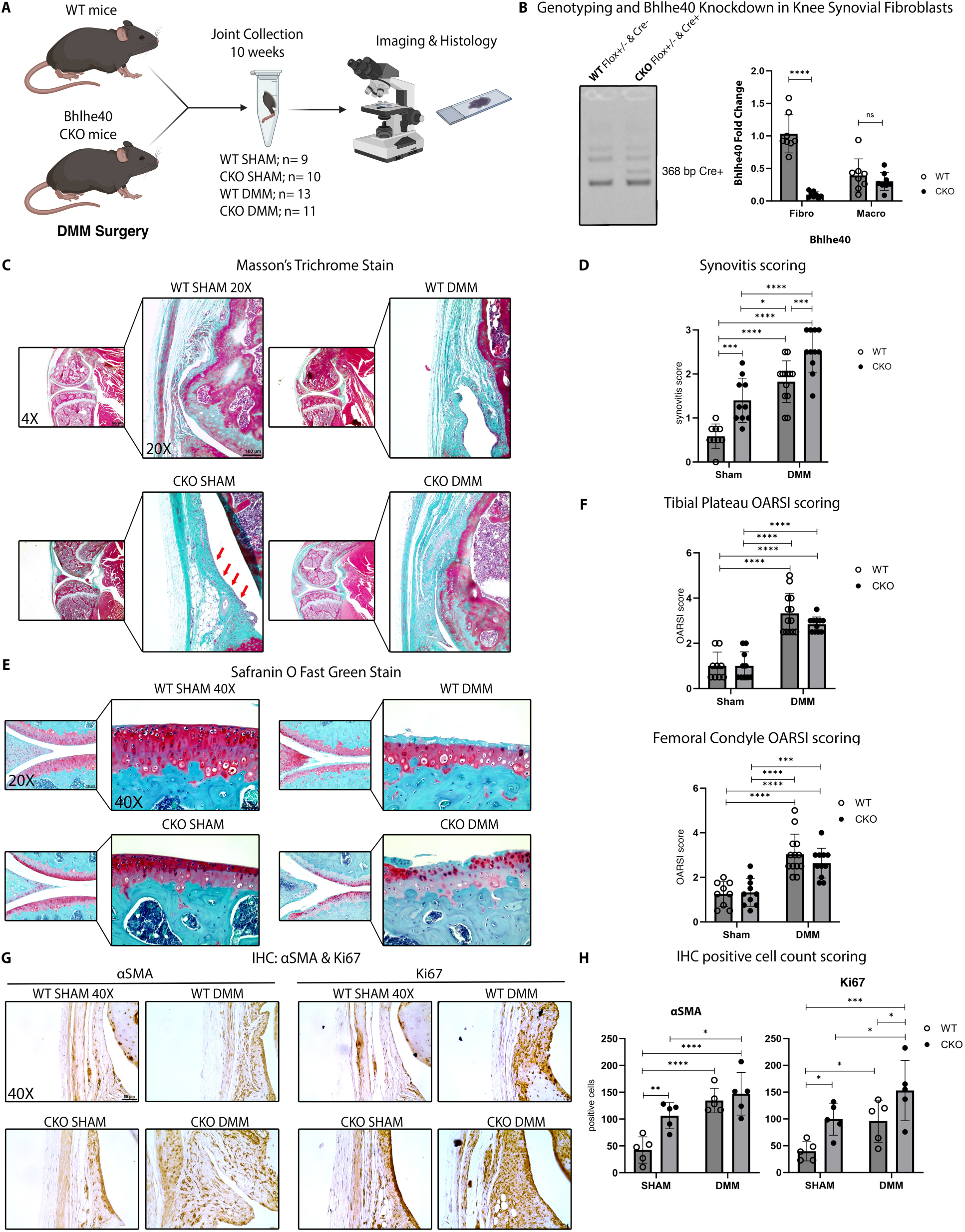
Fibroblast expression of *Bhlhe40* negatively regulates synovial fibrosis in surgically-induced murine KOA. A) Schematic of WT and CKO mouse joint collections subjected to imaging and histology. B) Genotyping of *Bhlhe40* CKO mice in Cre- and Cre+ mice. Graph showing *Bhlhe40* knockdown in fibroblasts and macrophages from synovia of WT and CKO mice. Data were analyzed by unpaired t-tests corrected for multiple testing. C) Masson’s trichrome staining at 4X and 20X images and (D) synovitis scoring of WT and CKO mice in both sham and DMM conditions. E) Knee joint sections stained with Safranin O/fast green at 20X and 40X magnification. F) OARSI cartilage scoring in both WT and CKO mice under sham and DMM conditions. G) αSMA and Ki67 IHC staining of mouse synovia. H) Positive cell count scoring for αSMA and Ki67 IHC. Synovitis, OARSI and IHC scoring data were analyzed by two-way ANOVA with post-hoc tests corrected using the two-stage step-up method of Benjamini, Krieger and Yekutieli. Data in (B), (D), (F) and (H) are presented as mean ± standard deviation. Not significant (n.s.) difference, *P ≤ 0.05, **P ≤ 0.01, ***P ≤ 0.001 and ****P ≤ 0.0001. Figure was created, in part, using Biorender.

12-week-old WT and CKO mice were subjected to sham or DMM surgery to evaluate the contribution of mouse fibroblast-intrinsic *Bhlhe40* to KOA pathogenesis. Interestingly, even in the absence of surgery, sham CKO mice exhibited significant development of synovial fibrosis, evidenced by increased thickening of the synovial lining and matrix deposition, as compared to sham WT mice. Furthermore, DMM CKO mice exhibited even more pronounced synovial fibrosis in comparison to DMM WT, Sham WT and Sham CKO mice (Fig. 7C), with DMM CKO mice having the highest synovial fibrosis scores (Fig. 7D). Cartilage integrity was also evaluated in safranin O-stained sections using OARSI grading criteria for mice (*34*) to assess differences between WT and CKO mice subjected to sham or DMM surgery. Prominent medial cartilage degeneration was present in WT and CKO mice subjected to DMM surgery, as compared to sham-surgical animals (Fig. 7E); however, no significant differences in OARSI scores of the tibial plateau (TP) or femoral condyle (FC) were observed in sham or DMM groups between genotypes (Fig. 7F).

Since we observed enhanced synovial thickening and fibrosis in CKO mice compared to WT mice, we further analyzed changes in protein expression of αSMA (activated fibroblasts), and Ki67, a marker of proliferation (*35*). IHC scoring of αSMA-stained sham mouse synovia showed significant increases in the number of αSMA-positive cells in the CKO mice compared to WT mice, however no significant differences were observed between DMM WT and DMM CKO mice (Fig. 7G, H). Ki67 scoring illustrated a significantly increased number of positive cells in sham CKO mice compared to sham WT mice. DMM CKO mouse synovia also had significantly higher Ki67 scores compared to DMM WT mouse synovia (Fig. 7H). Overall, these results show that loss of *Bhlhe40* leads to severe synovial fibrosis associated with increased fibroblast activation and proliferation *in vivo*.

### BHLHE40 overexpression suppresses TGF-β-induced fibroblast activation

Since *Bhlhe40* CKO mice developed severe synovial fibrosis, and BHLHE40 knockdown in vitro resulted in enhanced αSMA expression, fibroblast activation and stress fiber formation, we sought to overexpress BHLHE40 in KOA fibroblasts using a lentivirus approach to evaluate its effects on fibroblast activation. We first tested our lentivirus transduction efficiency and observed that 50 MOI was sufficient for transduction and target protein expression (Fig. 8A, B, fig. S10). We next used four experimental conditions to evaluate the effect of BHLHE40 on fibroblast activation: control lentivirus; control lentivirus + TGF-β; BHLHE40 lentivirus; and BHLHE40 lentivirus + TGF-β. TGF-β is a known inducer of fibroblast activation and was therefore used in this investigation (*36*). As expected, fibroblast activation and stress fiber formation were increased in cultures treated with control lentivirus + TGF-β. In contrast, BHLHE40 transduction alone did not result in significant morphological or phenotypic changes in fibroblasts compared to the control lentivirus-treated cultures alone. Notably, fibroblasts transduced with BHLHE40 lentivirus and treated with TGF-β exhibited a marked reduction in the activated phenotype, with reduced αSMA expression, stress fibers and a more spindle-shaped morphology, compared to cultures transduced with control lentivirus and treated with TGF-β. These findings show that overexpression of BHLHE40 suppresses fibroblast activation under fibrotic conditions and further supports its role as a crucial mediator of fibroblast activation.

**Fig. 8.**
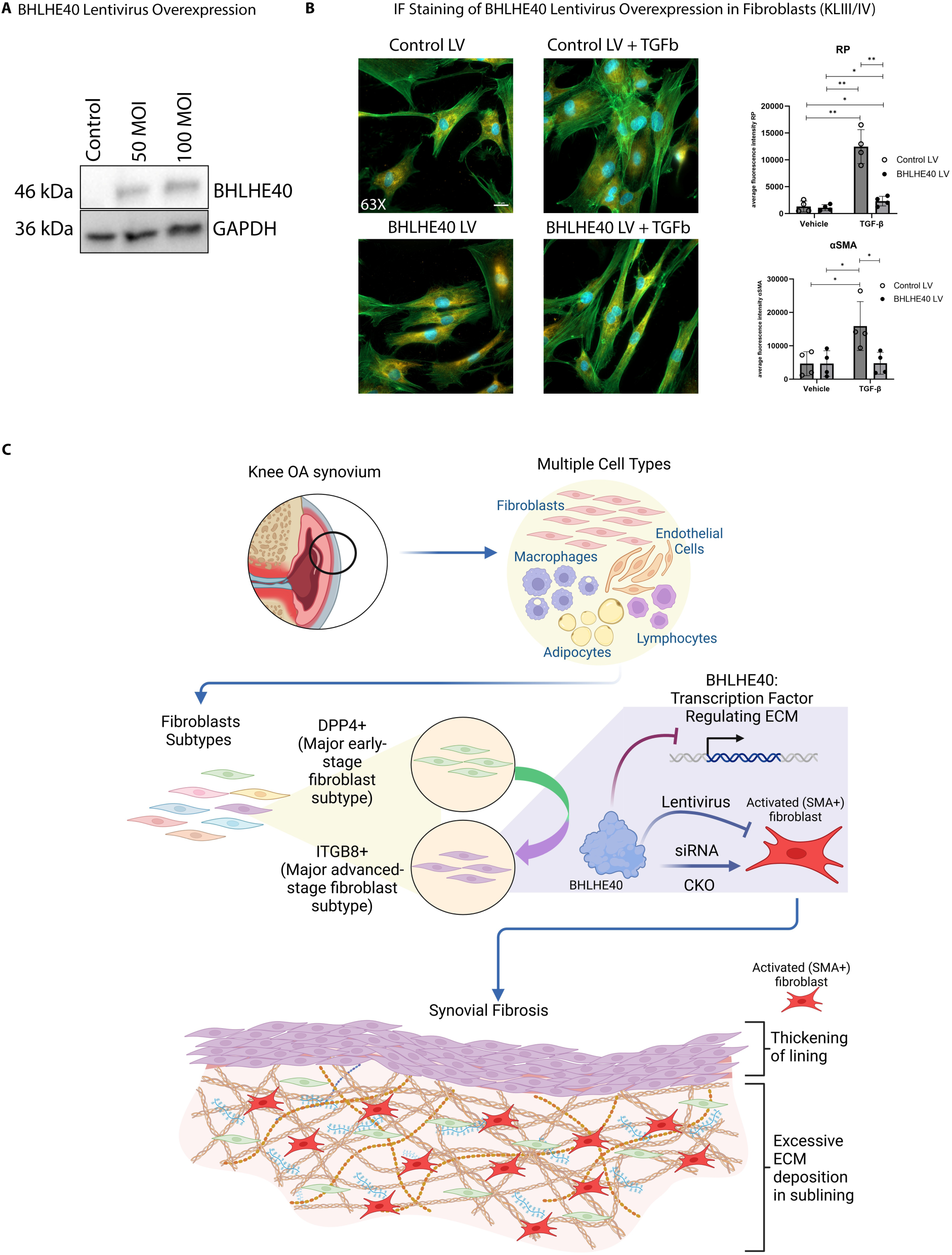
Overexpression of BHLHE40 attenuates TGF-β-induced fibroblast activation. A) Western blot (left) showing lentivirus transduction of 50 MOI or 100 MOI results similar BHLHE40 expression after 2 days in advanced-stage fibroblasts from KOA synovium. B) Fibroblast cultures were transduced with control or BHLHE40 lentivirus (50 MOI) for 24 hours and subsequently treated with or without TGF-β (20 ng/ml) for an additional 24 hours. Cells were immunostained to detect αSMA, and stained with rhodamine phalloidin to detect stress fibers, followed by imaging by fluorescence microscopy. C) Schematic diagram depicting key findings of this study. Multiple cell types were found in early and advanced-stage synovia, of which fibroblasts were the predominant cell type. DPP4+ fibroblasts were found to be the dominating fibroblast subtype in synovia of early-stage KOA while ITGB8+ fibroblasts were found to be the predominant subtype in advanced-stage tissues. BHLHE40, a crucial transcription factor in ITGB8+ fibroblasts, was identified to regulate fibroblast activation, ECM expression, and fibrosis. Figure was created, in part, using Biorender.

## DISCUSSION

Previous studies have subjected advanced-staged KOA synovia or fibroblasts from cultured from synovial tissues to single-cell RNA sequencing, identifying distinct heterogeneous synovial fibroblasts, unique transcriptomic signatures, and pain associated mechanisms (*20, 21, 37-39*). In this study, we provide a comprehensive analysis of cellular diversity within synovia of early and advanced stages of radiographic KOA. Using bulk RNA sequencing, we first show large transcriptomic differences in synovial tissues from advanced-versus early-stage KOA. Using single nuclei RNA sequencing, we found that fibroblasts were the major cell type in synovia, consistent with previous reports (*20, 38, 39*). The two predominant fibroblast populations identified in early- and advanced-stage KOA synovia were DPP4+ and ITGB8+ fibroblasts, respectively. Our trajectory analysis illustrated that advanced-stage ITGB8+ fibroblasts may be derived from DPP4+ fibroblasts over pathological pseudotime. These populations, and their putative transition were confirmed in a second cohort of synovia from healthy, early- and advanced-stage patients by flow-cytometry. Interestingly, similar fibroblast subtypes to human DPP4+ and ITGB8+ cells were observed when we evaluated synovia from a KOA mouse model, which also exhibited similar endotypic transitions of fibroblast subpopulations from early to advanced stages. This data supports the notion that fibroblast subtypes undergo progressive changes over different stages of KOA in both humans and in mouse models (Fig. 8C).

A previous study by Nanus et al. subjected cultured fibroblasts from early and advanced-staged knee OA synovia to single cell RNA sequencing (*38*), revealing differential transcriptomic profiles in synovial fibroblasts from patient reported painful sites. They also reported that synovial fibroblasts from painful sites can promote fibrosis, inflammation and neuron growth. In a separate study, Philpot et al. subjected advanced-stage knee OA synovial tissues to single cell RNA sequencing and described that synovial tissues from patients who experienced more pain were associated with macrophage exhaustion and exhibited synovial changes including increased angiogenesis or nerve growth (*37*). Thus, future investigations should consider analysis of associations of fibroblast subtypes found in select disease stages with pain.

Aberrant ECM production, deposition and mechanisms associated with fibrosis are associated with OA synovial pathology (*40, 41*). However, endogenous mechanisms that drive synovial fibrosis during KOA are poorly understood. In our study, both DPP4+ and ITGB8+ fibroblasts expressed genes associated with ECM-related pathways, with a greater proportion of matrisome-annotated genes expressed by the fibroblasts upregulated in advanced-stage synovia. Although ITGB8+ and DPP4+ fibroblasts were found to have gene-set enrichment for ECM-related pathways, it is crucial to recognize that they may operate through distinct mechanisms to influence ECM regulation at various stages of the disease, potentially mediated by unique transcriptional processes.

Using bioinformatic analyses, we identified the transcription factor *BHLHE40* to be a putative regulator of ECM-related gene expression, particularly in ITGB8+ fibroblasts. Notably, *BHLHE40* expression was increased in ITGB8+ compared to DPP4+ fibroblasts. Furthermore, in vitro knockdown of *BHLHE40* in advanced-stage synovial fibroblasts resulted in a prominent increase in the expression of fibrosis-related genes and promoted fibroblast activation associated with enhanced stress fiber formation and αSMA expression. We subsequently showed that in vitro overexpression of BHLHE40 using lentivirus transduction of KOA fibroblasts attenuated fibroblast activation under TGF-β stimulation. These results suggested that BHLHE40 plays an important role in ECM transcriptome regulation and fibroblast activation. This prompted us to generate *Bhlhe40* CKO mice with a targeted deletion in knee synovial fibroblasts in vivo. Deletion of *Bhlhe40* in synovial fibroblasts in vivo led to enhanced fibroblast activation and proliferation, and severe synovial fibrosis, further supporting the concept that BHLHE40 likely acts to reduce fibroblast activation and their contribution to ECM deposition in the synovium.

Together, these data suggest that BHLHE40 is a vital transcriptional repressor involved with regulating fibroblast activation and ECM-related activities, particularly in ITGB8+ cells from advanced-stage KOA synovium. This observation is consistent with BHLHE40’s suggested role as a transcriptional repressor involved in a variety of other cellular functions, including, but not limited to, regulation of the circadian rhythm (*29, 42*), differentiation of chondrocytes (*43*), regulation of lipogenesis (*44*), regeneration of skeletal muscle (*45*), and cell proliferation (*28, 46*). Knockdown of *BHLHE40* in cultured primary lung fibroblasts has previously been shown to increase the expression of *MMP1* and *MMP3*, supporting that BHLHE40 may be involved in regulating genes associated with fibrotic processes (*47*). However, opposite effects on fibrosis have also been observed in mice deficient in *Bhlhe40* using models of cardiac perivascular fibrosis and pulmonary fibrosis (*48, 49*). Thus, disease context may be important to the function of BHLHE40, including with respect to synovial fibrosis in KOA. Studies have implicated BHLHE40 to be involved in a number of pathways including the PI3K/Akt/mTOR pathway, p53-dependent senescence pathway, and others (*50*); however, how these mechanisms intersect with BHLHE40’s role as a negative regulator of fibroblast activation, ECM regulation and synovial fibrosis (as observed in the current study) requires further investigation.

There are limitations to our study that should be considered. For instance, not all genes in the transcriptomic profiles of fibroblast subsets from mice that were similar to human DPP4+ and ITGB8+ cells were the same; thus, species differences in contributions of these cell populations to disease progression might exist. Furthermore, our CKO mouse model is a targeted *Bhlhe40* deletion in *Col6a1*-expressing cells, which is also expressed in non-fibroblast cells, and may not be expressed in all fibroblasts. We also do not know when *Col6a1* is expressed during mouse joint development. Thus, uncharacterized cellular and tissue structural changes over developmental time may contribute to synovial changes observed, characterizations of which are beyond the scope of the current study. With respect to our studies using flow-cytometry, we also acknowledge the limitation of using PDPN as a marker for synovial fibroblasts (*51*), as its expression is low in fibroblasts from healthy and early-stage synovium (fig. S5C). This observation was reflected by higher PDPN+ cells identified in advanced-stage OA synovium. Thus, there may be PDPN-fibroblasts that were excluded in this analysis, as we strictly sorted for PDPN+ fibroblasts.

Overall, to our knowledge, this is the first study to comprehensively describe synovial fibroblast subtypes at early- and advanced-stages of KOA, identify endotype transitions of fibroblast populations involved in ECM-mediated regulation, and define BHLHE40 as a crucial mediator involved in ECM regulation, fibroblast activation and synovial fibrosis (Fig. 8b). Additional studies investigating the role of BHLHE40 to OA pathology and associated symptoms are necessary to fully appreciate its contributions to synovial fibrosis and OA pathogenesis. Finally, investigations into anti-fibrotic activity of BHLHE40 in synovial fibroblasts will be crucial to evaluate its potential as a therapeutic target.

## MATERIALS AND METHODS

### Study Design & OA Patient Recruitment

Patients were recruited from the Schroeder Arthritis Institute, University Health Network using Research Ethics Board (REB)–approved protocols and under informed written consent (early samples 16-5969-AE, advanced samples 07-0383-BE). All patients met the American College of Rheumatology’s definition of knee OA (*52*). This study ensures compliance with all relevant ethical regulations. For single-nuclei RNA sequencing and bulk RNA sequencing, OA human knee synovial tissue samples were retrieved from patients exhibiting KL I radiographic knee OA undergoing arthroscopic surgery and from with KL III/IV undergoing total knee arthroplasty (table S1A, B). Synovial tissue samples were processed for histology and stored in liquid nitrogen for sequencing. For our independent second cohort, KOA synovium was obtained from patients during total knee arthroplasty, routine knee arthroscopy or tissue donors for healthy controls. All patients provided informed consent to participate in this study (2019/1922-BC-06496; Commissie voor Medische Ethiek, UZ Gent). Healthy knee synovium samples were obtained from organ donors, shortly after circulatory arrest. Collection of healthy tissue was approved by UZ Gent Commissie voor Medische ethiek, ONZ-2022-0534 (table S1C).

### Human Synovium Histology

Harvested human knee OA synovial tissues were isolated and fixed in 10% neutral buffered formalin for at least 48 hours at 4°C. The tissues were embedded in paraffin and serial sections (4μm) were stained with Masson’s trichrome (Sigma-Aldrich) as per manufacturers guidelines. Sections were subsequently de-paraffinized and rehydrated.

### Single-Nuclei RNA Sequencing

Single-nuclei RNA sequencing was performed at the Princess Margaret Genomics Centre, UHN. OA synovial tissue was disaggregated into single-nuclei suspension, as described (*53*). To prepare for mechanical and enzymatic disaggregation, 30-50 mg of synovial tissue were cut into 1-2mm^3^ using a cold razor on dry ice. The tissues were covered with lysis buffer (1M sucrose, 1M CaCl_2_, 1M Mg(Ac)_2_, 1M Tris-HCl, Triton X-100, 0.5M EDTA pH 8, RNase Inhibitor (40U/μl), H_2_0) on ice and was mechanically disaggregated. A Kimble Dounce tissue grinder (Sigma-Aldrich) was used to homogenize the tissues and supernatants were collected in lysis buffer. Solutions were centrifuged three times at 800g for 10 minutes and washed with 2 ml wash buffer (1X PBS, 10% BSA, RNase Inhibitor (40U/μl), with final resuspensions using 1 ml wash buffer. The solutions were filtered using a 40μm Flowmi® cell strainer (Sigma-Aldrich) and transferred into a 1.5 ml LoBind tube on ice. Single-nuclei suspensions were assessed for nuclei quantity and viability with DAPI (Sigma-Aldrich) staining. FACS was used to sort-out the nuclei, which was subsequently centrifuged at 800g for 10 minutes. Nuclei pellets were resuspended in wash buffer to adjust the final concentrations to 1000 nuclei/μl.

The 10x Genomics Chromium system was used to construct 10x barcoded libraries. During the reactions, each 10X barcoded bead (v3.1) was encapsulated with one nucleus by portioning oil into one droplet. Reaction mixtures containing the nuclei, barcoded beads and partitioning oil were loaded separately on a10X chip and a collection of gel beads in emulsions were generated. This was further processed by performing reverse transcription (RT) reactions to form cDNA that shares 10x barcodes with all cDNA from individual nuclei of origin, followed by preparation of the 10x barcoded sequencing libraries, which were subjected to Illumina sequencing. The sequenced data were processed using Cell Ranger pipeline (v 6.1.2) by 10x Genomics (https://www.10xgenomics.com). Sequencing reads were aligned to human reference genome (GRCh38) and mouse reference genome (GRCm38) for mouse and human samples respectively, followed by filtering and correction of cell barcodes and Unique Molecular Identifiers (UMIs). Reads associated with retained barcodes were quantified and used to build gene count matrix.

### Single-Nuclei RNA Sequencing Analysis

Single-nuclei RNA sequencing of human and mouse datasets followed the standard procedures of filtering, normalization, dimensionality reduction and clustering, which was performed using the R package Seurat (v.4.1.0)(*54*). A total of 51,527 nuclei were sequenced from human tissues and 28,188 were sequenced from mouse tissues. All genes not detected in at least three nuclei and all nuclei expressing less than 200 genes were excluded from analyses. Nuclei with high mitochondrial content and potential doublets or multiplets were filtered out from downstream analyses. DoubletFinder was used for the detection of doublets (*55*). Early- and advanced-stage OA samples were merged for subsequent clustering and visualization. Batch correction was performed using the Harmony R package (v1.0)(*56*).

Data was normalized using the log normalization method and highly variable genes were selected using the variance stabilization (vst) method. PCA was then performed on the highly variable genes and the number of significant PCs to include for clustering was determined based on the elbow plot of standard deviations of PCs. A graph-based clustering method was implemented by calculating k-nearest neighbours, followed by Louvian modularity optimization to cluster cells. Non-linear dimensionality reduction and visualization was performed using UMAP (Uniform Manifold Approximation and Projection). Clusters were annotated based on canonical markers and differential gene expression testing was done to determine transcriptomic signatures for each cluster using the Wilcoxon Rank Sum test. Sub-clustering of fibroblast clusters was performed in a similar manner. A total of 11507 skeletal muscle cell nuclei were identified only in early samples and were excluded from the human dataset prior to cluster analysis, as they were considered contaminants from surrounding tissue.

Similarly, all mouse synovial samples from naïve, 2- and 10-week post-surgery samples were merged for clustering and visualization. The mouse fibroblast clusters were scored against human fibroblast clusters using ‘AddModuleScore’ function in Seurat.

### Flow Cytometry

Using synovia from our independent second cohort (UZ Gent), synovia was obtained from patients during total knee arthroplasty, routine knee arthroscopy or from tissue donor subjects.

Knee synovia was dissected from adjacent joint capsule/fat tissue and sectioned into 100 mg pieces for cryopreservation using cryostor following published protocols (*57*). Cryopreserved synovial samples were thawed, minced and immediately digested at 37C with 0.5mg/ml collagenase VIII and 0.1mg/ml DNase in phenol red free RPMI. Cells from the digested synovia were counted with trypan blue. The synovial single cell suspensions were stained in 96 well microplates using the following protocol: Cells were first stained with a fixable viability stain. After washing, the cells were treated with Fc receptor blocking reagent, followed by staining for surface antigens. Cells were fixed with PFA prior to acquisition on the BD Symphony. Two panels were designed with 15-25 channels each, depending on antibody and fluorochrome availability. Basic lineage markers were included to positively identify cells (e.g., fibroblasts were identified as CD31-CD235a-CD45-HLA-ABC+ cells, with CD90 and CD55 defining sublining and lining subsets, respectively). Specific markers identified by snRNA-seq were used to identify cell populations of interest identified in early vs. advanced stage OA synovia (e.g. ITGB8 for advanced-staged OA fibroblasts). Data was analyzed with FlowJo.

## Acknowledgments

Thank you to Dr. Brian Edelson for generously gifting the BHLHE40^f/f^ mice.

## Author contributions

Conceptualization: MK, DE, EG, JSR, RG

Methodology: KT, EG, JSR, MK, DE, RG, IJ, SL, GP

Investigation: KT, EG, AR, JSR, SV, CP, KH, JG, PK, FS, GP, SL, PP, RR, NM, JV, NA, IJ, RG, DE MK

Visualization: KT, EG, CP, IJ

Funding acquisition: MK, DE

Project administration: MK, DE, RG, EG, JR

Supervision: MK, DE

Writing – original draft: KT, EG

Writing – review & editing: KT, EG, AR, JR, SV, CP, FS, GP, KH, JG, PK, SL, PP, CV, RR, NM, JV, NA, IJ, RG, DE, MK

## Data and materials availability

All sequencing data including human snRNA-sequencing (GSE281826), human bulk RNA-sequencing (GSE281825) and mouse snRNA-sequencing (GSE281724), can be accessed through GEO super series accession ID: GSE281827.

