## Supplementary File for "A cell atlas of human and mouse synovium from early and advanced stages of knee osteoarthritis: BHLHE40 regulates fibroblast activation"

**This PDF file includes:**
Supplementary Materials and Methods
Supplementary Figures: Fig. S1- Fig. S10
Supplementary Table: Table S1- Table S8
References (58-71)

### Supplementary Materials and Methods

#### Bulk RNA sequencing

For bulk RNA sequencing analysis, n=6 KL I and n=8 KL III/IV radiographic knee OA synovial tissues were used. RNA was isolated from frozen synovia (~50 mg) using the Qiagen RNeasy Mini kit (Qiagen, 74104). Quality of total RNA isolated was assessed using an RNA Nano chip on an Agilent Bioanalyzer (Agilent, Santa Clara, CA, USA). Samples were fluorometrically quantified using a Qubit RNA BR assay (ThermoFisher) on a Denovix DS-11 spectrophotometer (Denovix, Wilmington, DE, USA). For each sample, 200 ng RNA was used to prepare sequencing libraries using the TruSeq Stranded Total RNA with RiboZero, as per manufacturer's recommendations (Illumina, San Diego, CA, USA), as previously described (*58*). Library quality was assessed on a high-sensitivity DNA chip on the Bioanalyzer (Agilent). Sequencing libraries were volumetrically pooled and sequenced on an Illumina NextSeq 550 sequencer for 150 paired end read-cycles at the Schroeder Arthritis Institute (Krembil, Toronto, ON, Canada).

Raw sequenced reads were assessed for quality with Cutadapt (v3.0)(*59*), used to maintain minimum read length of 25 bp post trimming of adapters along with trimming of N's. Splice-aware alignment of reads using a Hierarchical Graph FM index (HGFM) method was performed using HISAT2 software (v2.2.1)(*60*) against human reference genome (vGRCh38). To populate the abundance of transcripts based on the reference genome and transcriptome, StringTie (v2.1.4)(*61*) was run to generate outputs as table format files.

Gene expression read counts were analyzed to identify DEGs using Bioconductor package DESeq2 (version 1.36.0) (*62*). Lowly expressed genes, where less than 2 samples had counts less than or equal to 10, were filtered out. Genes with adjusted P value <0.05 and absolute log2 fold change >0.5 were considered to have significant altered expression. Unsupervised clustering was used to generate the heatmap using heatmap.2 function.

**Identification of Stage-specific Fibroblast Subcluster DEGs**

Cells identified as fibroblasts were clustered independently and differentially-expressed gene (DEG) lists were generated for each fibroblast subcluster using Seurat. Using a bulk-like approach, DEGs in fibroblasts from early- (KL I) and advanced-stage (KL III/IV) OA patient samples were also determined. Upregulated genes identified in early-stage samples were retained in the clusters dominated by nuclei contributed from early-stage synovium (clusters 1, 2, 4 and 6) and removed from clusters dominated by nuclei contributed from advanced-stage synovium samples (clusters 0, 3 and 5). Similarly, genes that were upregulated in advanced-stage patient samples were retained in clusters dominated by nuclei contributed from advanced-stage synovium samples (clusters 0, 3 and 5) and removed from clusters dominated by nuclei contributed from early-stage synovia (clusters 1, 2, 4 and 6). The resulting list of genes for each cluster represented cluster-specific DEGs associated with their predominant differential expression in early- or advanced- stage KOA synovial tissues.

**Trajectory Analysis**

Monocle (v2.34.0)(*63*) was used to investigate inferred trajectories between fibroblast clusters. SnRNA-seq data previously analyzed with Seurat was imported into monocle and highly variable genes with differential expression were selected to order and place cells along a trajectory based on their expression patterns across the cell population. Dimensionality reduction was performed using the ‘reduceDimension’ function with the reduction method set to "DDRTree" to visualize the trajectory in a reduced dimensional space. The trajectory was then constructed using the ‘plot_cell_trajectory’ function with default parameters. The clusters identified from Seurat analyses were superimposed onto the pseudotime trajectory. The kinetic trends of the expression levels of significant genes through pseudotime was depicted by a heatmap, which clustered the genes into modules that co-vary across pseudotime.

**Cell Crosstalk Analysis**

We used CellChat (version 2.1.2), an R package, to infer the cellular communication network between major cell types and fibroblast sub-types in human synovia. The “netVisual_circle” function in CellChat was employed to generate a circle plot, visualizing aggregated cell-cell communication networks. To examine complex ligand-receptor interactions between source and target cell types, we used the 'netVisual_chord_gene' function to generate a chord diagram. We analyzed outgoing communication patterns to gain insights into how various sub-clusters and signaling pathways coordinate with one another. We identified upregulated and down-regulated signaling ligand-receptor pairs based on the differential expression analysis (thresh.pc = 0.1, thresh.fc = 0.05, thresh.p = 0.05) between early- (KL I) and advanced-stage (KL III/IV) tissues. We then compared the interaction strengths of these pairs, using early (KL I) cluster 1 and advanced cluster 0 (KL III/IV) as senders and the receiver cell types.

**DMM Surgery**

The right knees of 12-week-old C57BL/6 [The Jackson Laboratory (RRID: IMSR_JAX:000664)] mice were subjected to destabilization of the medial meniscus surgery with synovium collected at 2-weeks (n=3) or 10-weeks (n=4) post-surgery (64). Synovia from naïve mice (n=4) were also collected for analysis. Extracted synovia were subjected to snRNA-seq, as described above. Of note, collections include some contaminating tissue including patella, ligament, fat and muscle, as previously described (65).

**Human Tissue Immunohistochemistry**

For immunohistochemistry (IHC), formalin-fixed, paraffin embedded tissue sections were deparaffinized and rehydrated in a graded series of alcohol washes and with distilled water. Antigen retrieval was performed by incubating slides in 10mM citrate buffer (pH 6.0) at 93°C. Endogenous peroxidase activity was blocked by incubation in 3% H2O2 in the dark at RT. The sections were blocked with normal blocking serum for 30 minutes (Goat normal serum, 5425S, Cell Signaling; rabbit serum, 16120099, ThermoFisher). Human sections were then incubated with the following primary antibodies at 4°C overnight: ITGB8 (1:200, PA5-97884, ThermoFisher), and DPP4 (1:50, AF1180, Bio-techne). Sections were incubated with respective biotinylated secondary antibodies (1:200) for 30 minutes: goat anti-rabbit (BA-1000, Vector Laboratories), rabbit anti-goat (BA-5000, Vector Laboratories), or goat anti-mouse (BA-9200, Vector Laboratories), followed by incubation with Vectastain Elite ABC kit (VECTPK6100, Vector Laboratories), and developed with DAB peroxidase (HRP) Substrate Kit (VECTSK4100, Biolynx), as per manufacturer's direction. Sections were counterstained with hematoxylin (H-3404-100, Vector Laboratories). Quantification of IHC-stained human tissues was performed by counting the total cells that stained positive for each antigen in areas of the synovial lining and sub-lining areas, and in whole synovium of the mouse tissue. IHC-stained tissues were assessed by two independent blinded scorers and the final values were calculated as means of total positive cell counts.

**Pathway and Transcription factor Analyses**

The list of genes from each cluster linked to early- or advanced-stage fibroblasts were used to perform pathway and gene ontology enrichment analysis. Pathway enrichment analyses were performed using integrated pathway database pathDIP 4 (http://ophid.utoronto.ca/pathDIP) (*66*) API in R 4.0.3, excluding KEGG, OntoCancro, ACSN2, RB-Pathways and WikiPathways sources, which were not relevant for the current analyses. Only pathways with adjusted P<0.01 were further considered. Gene Ontology Biological Processes enrichment analysis was performed using clusterProfiler v3.16.1 (67) in R and retaining terms with adjusted P< 0.01. ECM related genes from DEG lists of cluster 0 and cluster 1 that were associated with identified pathways of cluster 0 and 1 were extracted. Catrin database v 1.0.6.2.b (68) was used to identify transcription factors (TF) targeting the ECM genes for each cluster. Hypergeometric test to identify upstream regulators was performed in R, and TFs with adjusted P<0.01 were retained. Furthermore, we only considered TFs that targeted DPP4+ or ITGB8+ fibroblast genes. A network of pathway – gene – TF interactions was built using NAViGaTOR v3.0.16 (69). Interactions among genes were retrieved using IID v2021-05 (70). The network was then reduced to show only terms related to extracellular matrix processes.

**Cell Culture**

Primary human synovial fibroblasts were isolated from the synovial tissues obtained from patients with KL III/IV radiographic knee OA. Tissues were washed with Dulbecco's Modified Eagle medium (DMEM, 11995- 065, Life Technologies/Gibco) supplemented with 10% fetal bovine serum, digested with trypsin (T9201, Sigma), and treated with collagenase (C0130, Sigma). The cells were then cultured in DMEM with 10% FBS and 1% PenStrep (450-201-EL, Wisent) in a humidified incubator with 5% $CO_2$ at 37°C.

**Gene Silencing by siRNA Transfection**

Human fibroblasts were cultured to 80% confluency at P4-P5. 12-well plates were filled with DMEM (10% FBS) and 15 nM BHLHE40 siRNA (GS8553, Qiagen), or a negative control siRNA (4390843, ThermoFisher), with 10nM of Lipofectamine RNAiMAX (13778100, ThermoFisher) and then plated with fibroblasts at a density of 100,000 cells per well for 24 h. After 24 hours, cells were starved with 1% FBS overnight or utilized for immunofluorescence staining. Starved cells were then washed with cold PBS and lysed with TriZol. Cell lysates were immediately stored for long term storage at -80°C or immediately underwent RNA isolation.

**RT-qPCR**

The TRIzol-chloroform method was used to isolate RNA. The quality of RNA was assessed using a Nanodrop and cDNA was generated according to manufacturer's instruction using the QuantiTect$^{TM}$ Reverse Transcription kit (catalog 205311; Qiagen). SYBR™ Green PCR Master Mix (catalog 4309155; ThermoFisher) was used to perform RT-QPCR, as per manufacturers' directions, using the following primers: BHLHE40 – forward (5'GGACAGCAAGGAGACCTACA3') reverse (5' AGTGCTTTCACATGCTTCAAG3'); GAPDH – forward (5'AAGGTGAAGGTCGGAGTCAAC3') reverse (5'GGGGTCATTGATGGCAACAATA3'). CT values were normalized to GAPDH using the ΔCT calculation. Relative gene expression was plotted, and significance was determined using a two-tailed paired T-test.

**NanoString**

RNA extracted from human synovium were transported on dry ice to NanoString Technologies (Toronto, Canada) for gene expression analysis. The nCounter Human Fibrosis V2 Panel was used, which includes 760 genes that cover the pathways and processes related to fibrosis alongside 10 internal reference genes for data normalization, and 17 custom genes added to the panel including, *ADAM10*, *ADMATS3*, *BHLHE40*, *CD63*, *CSGALNACT1*, *CYP1B1*, *HTRA1*, *ITGAV*, *ITGB8*, *ITGBL1*, *NTN4*, *PLOD2*, *SAT1*, *SH3PXD2A*, *TNC*, *VAV3* and *VEGFC*. The nCounter data was analyzed with nSolver 4.0 and normalized to standard select housekeeping genes including *ACAD9*, *ARMH9* and *GUSB*. Differential expression analysis was conducted using a paired-sample design for each BHLHE40 treatment compared to control. Univariable generalized least-squares regression models were fit for paired comparisons. $Log_{10}$-transformed read counts were regressed on treatment type, with a random intercept for each subject ID. P-values were adjusted for multiple comparisons using the Benjamini-Hochberg method with a false discovery rate of 5%. To create the heatmaps, $log_{10}$-transformed read counts were subjected to gene-wise re-scaling to mean 0 and standard deviation 1. Differential expression analysis was conducted in R statistical software. $Log_2$-fold changes of BHLHE40 siRNA treatment versus control were graphed for all genes that were significantly different

**Cell Culture Immunofluorescence**

SiRNA treated fibroblast cultures utilized for immunofluorescence staining of αSMA and rhodamine phalloidin were plated on coverslips (12541015CA, Fisher Scientific) in 6-well plates at a density of 150,000 cells per well. After treatment with siRNA for 48 hours, cultures were fixed with 4% paraformaldehyde and incubated with a primary αSMA antibody (A2547, Sigma Aldrich,

1:400) overnight at 4°C, followed by a secondary antibody (AF568, Invitrogen, 1:300) for 1 hour in the dark at room temperature, and lastly with rhodamine phalloidin stain (A12379, Fisherscientific) for 1 hour in the dark at room temperature. Cells were stained with DAPI for 5 minutes in the dark and mounted with anti-fade mounting medium (S3023, Agilent) and left overnight to dry in the dark.

**Fibroblast- specific BHLHE40 Conditional Knockout Mice**

*Bhlhe40*$^{f/f}$ mice were generously gifted by Dr. Brian Edelson (Washington University) (71). *Col6a1-cre*/+ mice (B6.Cg-Tg(Col6a1-cre)1Gkl/Flmg), were purchased from the European Mutant Mouse Archive repository as sperm and re-derived in-house. Fibroblast-specific *Bhlhe40* knockout mice were generated by crossing *Bhlhe40*$^{f/f}$ mice to *Col6a1-cre/+* mice to generate *Col6a1-cre/+; Bhlhe40*$^{f/f}$ mice. *Cre-; Bhlhe40*$^{f/f}$ mice were used as controls. Right knee joints from WT or *Bhlhe40* CKO mice were subjected to sham or DMM surgeries (WT sham, n=9; CKO sham, =10; WT DMM, n=13; CKO DMM, n=11) and collected 10 weeks post-surgery. Genotyping was assessed by standard PCR and confirmed by sorting synovial fibroblasts and macrophages for qPCR.

To sort synovial fibroblasts and macrophages for confirmation of cell-specific *Bhlhe40* deletion, mouse knee synovium was dissected as above and subjected to enzymatic digestion using 1 mg/ml collagenase IV, 0.75 mg/ml collagenase VIII and 0.1 mg/ml DNAse. Cells were stained with CD31-APC.Cy7, F/480-APC, CD45-PB, Ly6G-FITC, CD11b-BV785, PDPN-PECy7 and 7AAD. Live fibroblasts (CD31-CD45-PDPN+) and macrophages (CD31-CD45+Ly6G-CD11b+F4/80+) were sorted using the ARIA III (BD). RNA was extracted from cells using RNease micro kit (Qiagen) before qPCR for the genes of interest. *Eef1a* (F CGTCAGAACGCAGGTGTTG, R

TCGATGGTTCGCTTGTCGAT) and *Rpl13a* (F AAGCAGGTACTTCTGGGCCG, R CCTCGGGAGGGGTTGGTATT) were used as housekeepers for *Bhlhe40* (F CGTTGAAGCACGTGAAAGCA, R TCCCGACAAATCACCAGCTT).

**Mouse Histology and Immunohistochemistry**

Mouse joint tissues were collected and fixed 10% neutral buffered formalin for at least 48 hours at room temperature, washed in PBS, transferred into 70% ethanol and transported to The Centre for Phenogenomics for decalcification, embedding, processing and staining with Masson's trichrome (HT10516, Sigma Aldrich) (26367-04, Electron Microscopy Sciences) and Safranin O (S2255-100G, Signma Aldrich)/Fast Green staining (2353-45-9, Bio Basic Canada) (26367-02, Electron Microscopy Sciences). IHC was performed on mouse sections, using primary antibodies Ki67 (1:200, ab15580, Abcam) or α-SMA (1:200, A2547, Sigma), with total positive cell scoring performed, as described above.

**Construction and Production of Lentivirus Vector**

The full-length human *BHLHE40* gene tagged with eGFP using T2A peptide and control eGFP transgene were individually cloned into the pRS-EF1a-MCS-WPRE lentiviral vector (Tailored Genes Inc, Ontario, Canada) using In-Fusion HD cloning kit (638946, In-Fusion® Snap Assembly Value Bundle Takara Bio USA Inc, San Jose, CA, USA). The source sequence for huBHLHE40/HA-T2A was amplified from a pAAV[Exp]-EF1A>hBHLHE40[NM_003670.3](ns)/HA:T2A:Nluc:WPRE3 vector (VB240712-1285nmb; Vector Builder, Chicago, IL). Lentivirus vectors (LVV) were produced by transfection of HEK293T cells with pRS-EF1a-hBHLHE40/HA-T2A-eGFP-WPRE/pRS-EF1a-eGFP-WPRE pDNAs and with packaging plasmids pLenti gag-pol, pRSV Rev, and pHCMV- VSV-G (Tailored Genes Inc, Ontario, Canada). The supernatant containing the LVVs was collected at 48 h post-transfection, filtered using 0.45-micron filters and concentrated/purified using Amicon ultrafiltration (EMD Millipore corporation, MA, USA). Infectious titer was determined by transducing human osteosarcoma cells with serial dilution of LVV, genomic DNA isolation from transduced cells, and determining vector copy number in transduced cells by digital PCR using primers/probe specific to WPRE sequence.

Fibroblasts were infected with the LVV at a multiplicity of infection (MOI) of 50, in the presence of 8 μg/ml polybrene (Sigma-Aldrich, St. Louis, MO, USA). After overnight incubation with LVV, supernatant was removed, and fresh culture media was added prior to treating the transduced cells with TGFβ and they were further processed for immunofluorescence, as described above. Average immunofluorescence intensity was measured using HALO 3.6 Area quantification and data was analyzed by two-way ANOVA and post-hoc tests with p-values adjusted for multiple comparisons using the Benjamini-Hochberg correction, with adjusted $p<0.05$ considered significant.

**Statistical Analysis**

The proportions of human fibroblast subtypes in early- versus advanced-stage synovia were analyzed using two-tailed t-tests and corrected for multiple comparisons by controlling the false discovery rate using the two-stage step-up method of Benjamini, Krieger and Yekutieli. Data are expressed as mean ± standard deviation (SD). The default statistical tests implemented in the functions from the Seurat, Monocle, and CellChat R packages were applied for snRNA-seq data analysis. These packages include FDR correction to account for multiple comparisons in large datasets. OARSI, synovitis scores and IHC positive cell count for CKO mice (Ki67 & α-SMA) were statistically analyzed using a two-way ANOVA and post-hoc tests with p-values corrected for multiple comparisons by controlling the false discovery rate using the two-stage step-up method of Benjamini, Krieger and Yekutieli. Finally, IHC positive cell count scoring for ITGB8 and DPP4 were analyzed using an unpaired, nonparametric Mann Whitney test. After appropriate adjustment, all tests used p-value <0.05 to define statistical significance.

**Fig. S1. Severity of synovial fibrosis in KOA and expression of top DEGs in fibroblast subtypes.** A) Scoring of synovial fibrosis of Masson's Trichrome staining of early- (KLI, n=8) and advanced-stage (KLIII/IV, n=8) radiographic KOA synovial tissues. Unpaired Mann Whitney test was performed. B) Feature density plots displaying the expression of the top third to fifth markers (identified as differentially-expressed for each cluster using Wilcoxon rank sum test) for early (1,2,4,6) and late (0,3,5) fibroblast clusters along with violin plots below showing the expression levels across all fibroblast subclusters from human knee OA synovia. C) Bar graph showing proportion of nuclei contributing to fibroblast subclusters from early- and advanced-stage KOA synovial tissues. Data was analyzed by multiple unpaired t-tests with FDR correction using the Benjamini, Krieger and Yekutieli two stage step up method. Data in (A) and (C) are presented as mean ± standard deviation. *, P ≤ 0.05, **, P ≤ 0.01 and ***, P ≤ 0.001.

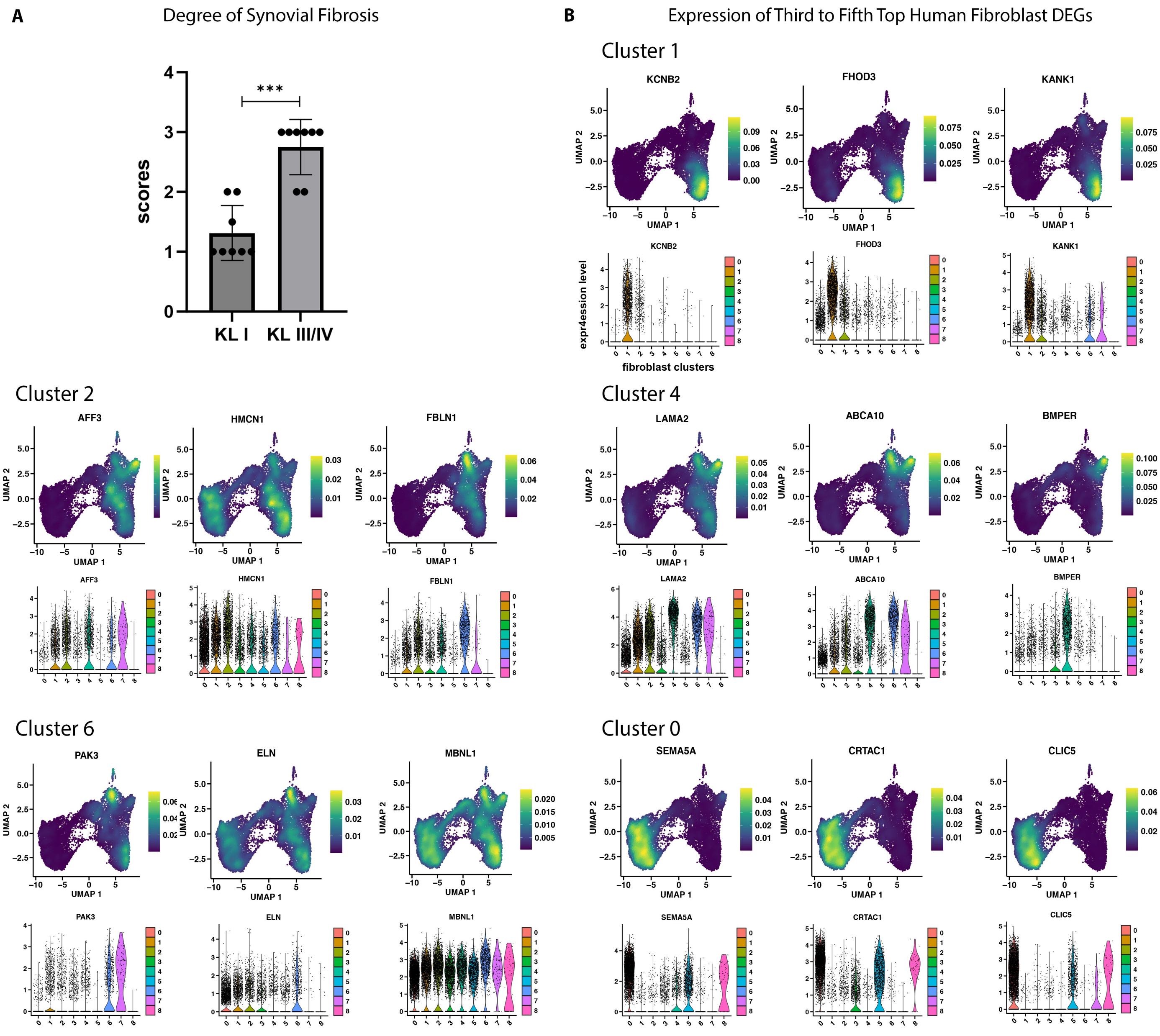

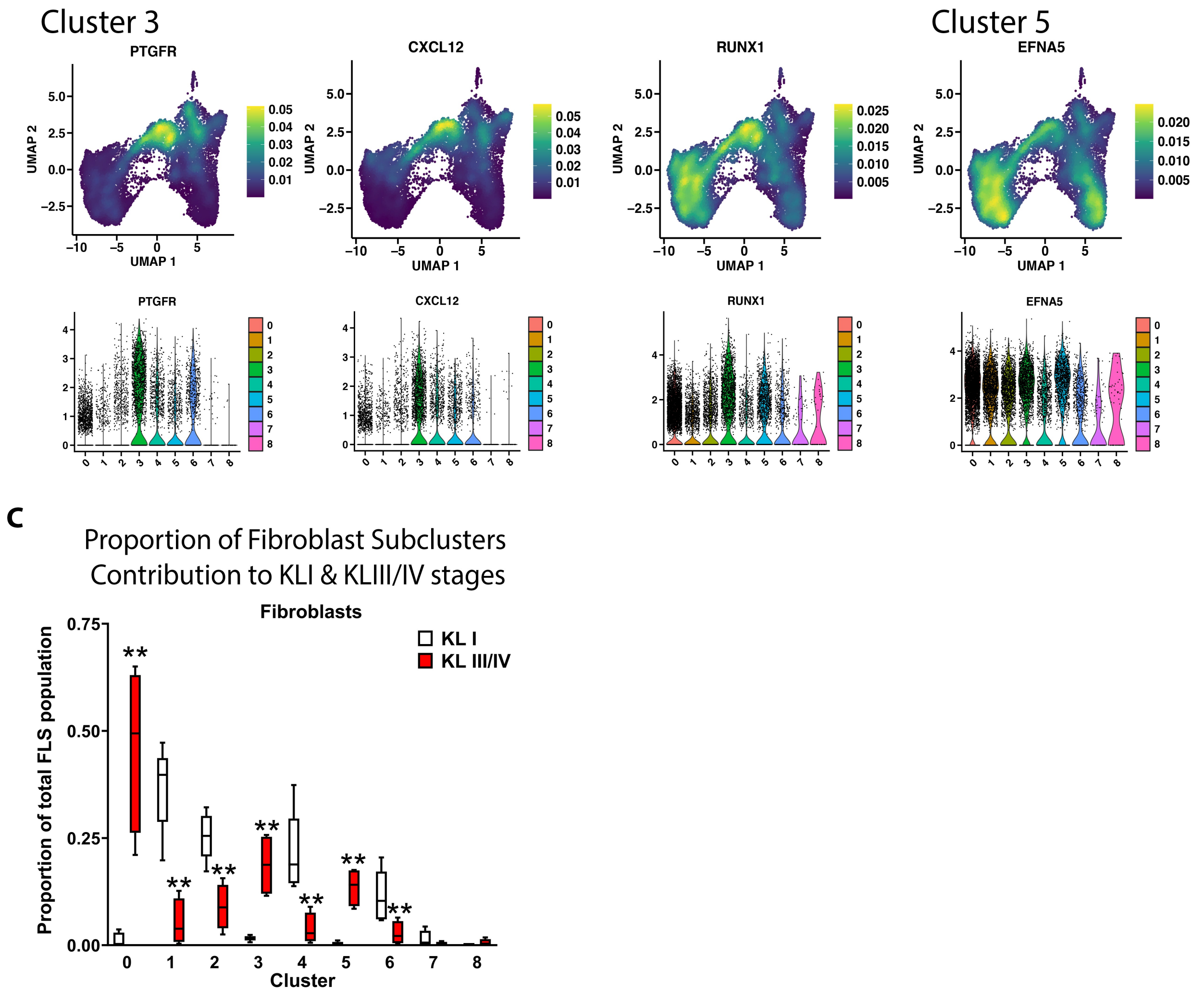
Cluster 3
Cluster 5
PTGFR
CXCL12
RUNX1
EFNA5
UMAP 1
UMAP 2
C
Proportion of Fibroblast Subclusters
Contribution to KLI & KLIII/IV stages
Fibroblasts
KL I
KL III/IV
Proportion of total FLS population
Cluster

**Fig. S2. Unique list of differentially-expressed genes in fibroblast subclusters identified from single nuclei RNA sequencing of 5 early- (KLI) and 4 advanced-stage (KL III/IV) KOA synovial samples.** The genes are ordered by decreasing average log2 fold change values. DEGs were identified using a cut-off of adjusted $p<0.05$ and average log2FC $>0.5$.

**Cluster 0:** PRG4, ITGB8, SEMA5A, CRTAC1, CLIC5, ANK3, SEMA3A, FN1, ZNF385B, COL22A1, ITGBL1, VEGFC, TMEM196, ERRFI1, RGS6, GPR1, AC008691.1, AP002370.2, XYLT1, CEMIP, HTRA1, MYPN, DAPK1, DIRC3, GABRA4, CLU, GABRB1, FGF10, SOX5, KYNU, SHANK2, ANO5, BCAT1, KCNQ5, DIRC3-AS1, ANKH, ZBTB7C, NTN4, FAM49A, TRHDE, SV2B, CTTNBP2, ADAMTS16, HTRA4, SIPA1L1, ZSWIM6, ADAMTSL1, FOXO1, THBS4, COLEC12, MT2A, SLC2A12, MTUS2, SPARCL1, PSD3, PLEKHA1, SORBS2, VCAM1, AC005237.1, GLCCI1, UST, HBEGF, FAM135B, SLC4A7, AC084816.1, CSGALNACT1, ADAM10, EML1, MAGI2, EPB41L1, ARHGAP28, RYR2, AC005062.1, VWC2, FAM155A, SEMA3E, ADAMTS3, GALNT10, CNTNAP3, FGF10-AS1, PPM1L, ADGRA3, CNTNAP3B, SNED1, RERG, SLC1A1, ERBB4, PPP4R4, DDAH1, PTPN22, CRISPLD1, TIMP1, STEAP4, ARNTL2, TNC, AC012636.1, BTC, KALRN, AP003086.1, DOCK3, ARHGAP20, ZDBF2, MITF, TIMP3, SH3PXD2A, TRHDE-AS1, CUX1, IGFBP5, CAV1, STOX2, PLOD2, CFI, MET, PDGFC, APLP2, ITGAV, CDK14, VPS13A, ICA1, DEC1, RAPH1, GULP1, ZNF608, NALCN, CEMIP2, SEMA6D, MTERF4, ERC1, DISP1, PDLIM5, EPHA6, PDE1A, AL355612.1, UBA6-AS1, MAGI1, SOX6, GPC5, NEDD4L, AGFG1, CEP85L, TENT5A, SYN3, CHST11, GAB2, TPD52L1, DCBLD1, AL391117.1, ADAMTS17, STK38L, SAMD5, SEPTIN9, PPARG, AL365295.1, CDON, TSPAN15, PPP1CC, ZEB2, SGK1, COL5A2, AC107021.1, DAB2, S100A6, B2M, LRRC2, KHDRBS3, C1GALT1, UNC5C, CCN2, RETREG1, MSI2, CDH23, AL136441.1, GASK1B, MGAT4C, SESN3, S100A4, IGF2BP2, LMO4, ENAH, BACH1, SAT1, GAREM1, GRAMD1B, AC098829.1, C1QTNF7, CD9, MYO10, CACNA1D, PRKCA, PRUNE2, PRDM6, LRP1B, MAGI2-AS3, TLE4, CHD7, LYN, MAPK10, MAP7, AC073475.1, EPB41, GSAP, GTF2IRD1, SLC8A1, PLCG2, SAMD12, LTBP3, MGP, EPHA3, MALAT1, GPC6, GOLM1, PVT1, ITM2B, MRTFB, SCARA3, FNIP2, ADGRB3, CD63, CWC22, RPLP1, HLA-A, SLC39A14, MTRNR2L12, TRPS1, RPL41, SPAG16, CYP1B1, CREB3L2, VAV3, SH3RF1, NDEL1, DPYD, FOXP2, ITPR2, ROR2

**Cluster 1:** DOCK4, PXDNL, KCNB2, FHOD3, KANK1, FBN1, MFAP5, NHSL1, NTM, SMURF2, STK32B, GUCY1A2, NAV3, TRIO, AC093425.1, KCTD8, PCOLCE2, MAST4, PTGIS, SLC4A4, ADAMTS5, NEGR1, CD55, TBC1D12, SHISA6, ADGRD1, DPP4, FRMD4A, ITGA11, TGFBR3, HUNK, PAMR1, STXBP6, CELF2, LIMCH1, DENND2A, NOVA1, FSTL1, EBF2, PDGFD, TMEM131L, ZEB1, RRAS2, STON2, SNTB1, LMX1A, ABLIM1, DCLK1, EDIL3, MSR1, CBLB, SDK1, PROCR, PLXNA2, GAS7, FAR2, FNDC1, PXN, AL008633.1, UTRN, CYTOR, CRIM1, TNXB, NID1, SPSB4, HDAC7, FBXL7, CPE, EHBP1, ARHGAP29, EFHD1, ELOVL6, CD34, C1orf21, FBLN2, DUXAP8, FMNL2, GALNT15, ARHGEF10, TC2N, LURAP1L, NCKAP5, USP54, ARHGEF3, RBMS3, PI16, MID1, CACNA2D3, STIM1, ACKR3, FREM1, CHL1, PDE7A, RECK, NOX4, PPP2R2A, MBP, RALGPS2, SH3D19, VIT, GALNT16, SRPX2, PRKD1, AOX1, EMILIN2, NTN1, TPM1, APBA1, DDX60L, ADD3, IGFBP6, ANOS1, ATP2B1, ST7, GRB10, DISC1, SDC2, ZHX3, OPHN1, GLIPR2, DPYSL3, SESTD1, GFRA1, FLRT2, APBB2, FAM107B, MGST1, C2orf27A, AGAP1, PLEKHG1, CORO2B, ZBTB16, DDR2, SRPK2, SAMD4A, ST3GAL5, SWAP70, FAT4, ATP11A, ABCA9, AL139383.1, CAB39L, TTLL7, CARMIL1, AL031599.1, TRERF1, PARD3B, DCN, SETD7, STARD9, TWIST2, RAP1A, UPF3A, PALM, ADAM33, LTBP4, HEG1, VGLL3, CPQ, WWC2, FBXL20,

STK39, WASF1, PRKCE, RFTN1, TTN, RADIL, ZNF438, EEF2K, DCBLD2, NLGN1, LEPR, BASP1, AKAP12, ARHGAP10, RAP1B, LPP, NEB, MYBPC1, STK38, STAC, DIPK1A, FOXN3, CTIF, FER, TET1, DMD, ARHGAP42, MYOM1, SMG6, TGFBR2, SPTBN1, LTBP1, TBC1D8, MAP1B, PRRX1, KCND2, PLXDC2, BACH2, PCNX2, ZNF521, NIN, UTY, CACHD1, PCED1B, HIVEP1, RIMKLB, MCUB, SUGCT, PDE7B, TNS3, ATP10A, KIFAP3, ARHGAP12

**Cluster 2:** ROBO2, DHRS3, AFF3, HMCN1, FBLN1, SETBP1, GSN, COL12A1, CNKSR2, LHFPL6, PRICKLE1, ANK2, CACNB4, PODN, SPOCK1, NIBAN1, IL13RA1, BMP5, AC108734.4, TRIP10, COL5A1, TANC2, MEOX2, PIK3R1, TTTY14, COLGALT2, LTBP2, VPS13D, OSBPL1A, PELI2, AGTR1, PER3, HMCN2, ANKRD36C, EBF3, RNF144A, OAF, PCSK5, NFIX

**Cluster 3:** KAZN, GLIS3, PTGFR, AL357873.1, KAZN-AS1, CXCL12, RUNX1, IGFBP4, ELL2, MECOM, PPP3CA, TMTC2, PDE4D, PDE4B, KSR1, PRR5L, REV3L, KCND3, MCTP2, CFAP69, HIF1A, PTPRD, MAML2, ST5, PLA2G2A, LRMDA, ZIC1, FAM13C, RNF24, GMDS, BOC, LNX1, OSMR, TBX15, TLL1, PALMD, LUM, PTPRF, ADAMTS9-AS2

**Cluster 4:** LRRTM4, COL15A1, AL590807.1, LAMA2, ABCA10, BMPER, SMOC2, ABCA6, ABCA8, GREB1L, ARHGAP24, AC024230.1, EDA, PID1, HMGCLL1, LAMC1, AMPH, SVEP1, ADAMTS12, HSPG2, ABCA9-AS1, PDE5A, GHR, PDZRN3, EGFR, FBLN5, ARHGAP6, FAP, SPATA6, BCL2, ECHDC2, DPT, PDZD2, KIAA1217, ANKS1B, PTPRK, GAB1, NECTIN3, SPRED2, RUNX1T1, NFIA, ARHGAP26, ZFPM2, ADGRL2, NR3C1, ARHGAP21, DOCK9, TCF7L1, SYTL4, AC119674.1, MAN1C1, CACNA2D1, FGD4, CPED1, CHD9, PARD3, PLAGL1, PLXDC1, PBX1, NFIB, SSH2, AKT3, LRRK2, FILIP1L, RAB30, INSR, ZC2HC1A, CYTH3, TRDN, AL445426.1, NR1D2, ADAMTSL3

**Cluster 5:** AC092958.1, TEX41, STEAP2, EFNA5

**Cluster 6:** CNTN4, NRP1, PAK3, ELN, MBNL1, C11orf80, ITGA9**,** SORBS1, GLI2, EYA2, PPP1R12B, PLEKHA5, USP53, ZNF704, PALLD, AGL, NBEAL1, MBOAT1, RGL1, BNC2, SPECC1L

**Cluster 7:** MTSS1, TRIM22, SYNPO2, KLF7, MGLL

**Fig. S3. Comparison of gene signatures of fibroblast clusters in human OA synovia with fibroblast sub-types defined in published studies.** A) Collective expression measure of gene signatures of synovial subintimal fibroblasts (SSF) and synovial intimal fibroblasts (SIF) identified in Chou et al. *(20)* scores our early-stage predominant clusters (1, 2, 4 & 6) high for SSF and advanced-stage predominant clusters (0 & 5) for SIF. B) Collective expression measure of the gene signature (top 20 differentially-expressed genes) of four fibroblast clusters (SC-F1, SCF2, SC-F3, SC-F4) identified by Zhang et al. *(19)*. Our early-stage fibroblasts clusters (1, 2, 4 & 6) score high for CD34+ SC-F1 fibroblast cluster and our advanced-stage fibroblast clusters (0 & 5) have a signature similar to the CD55+ SC-F4 fibroblast cluster. Clusters 1 and 2 score relatively high for the SC-F3 cluster and cluster 3 scores high for the HLD-DRA+ SC-F2 clusters.

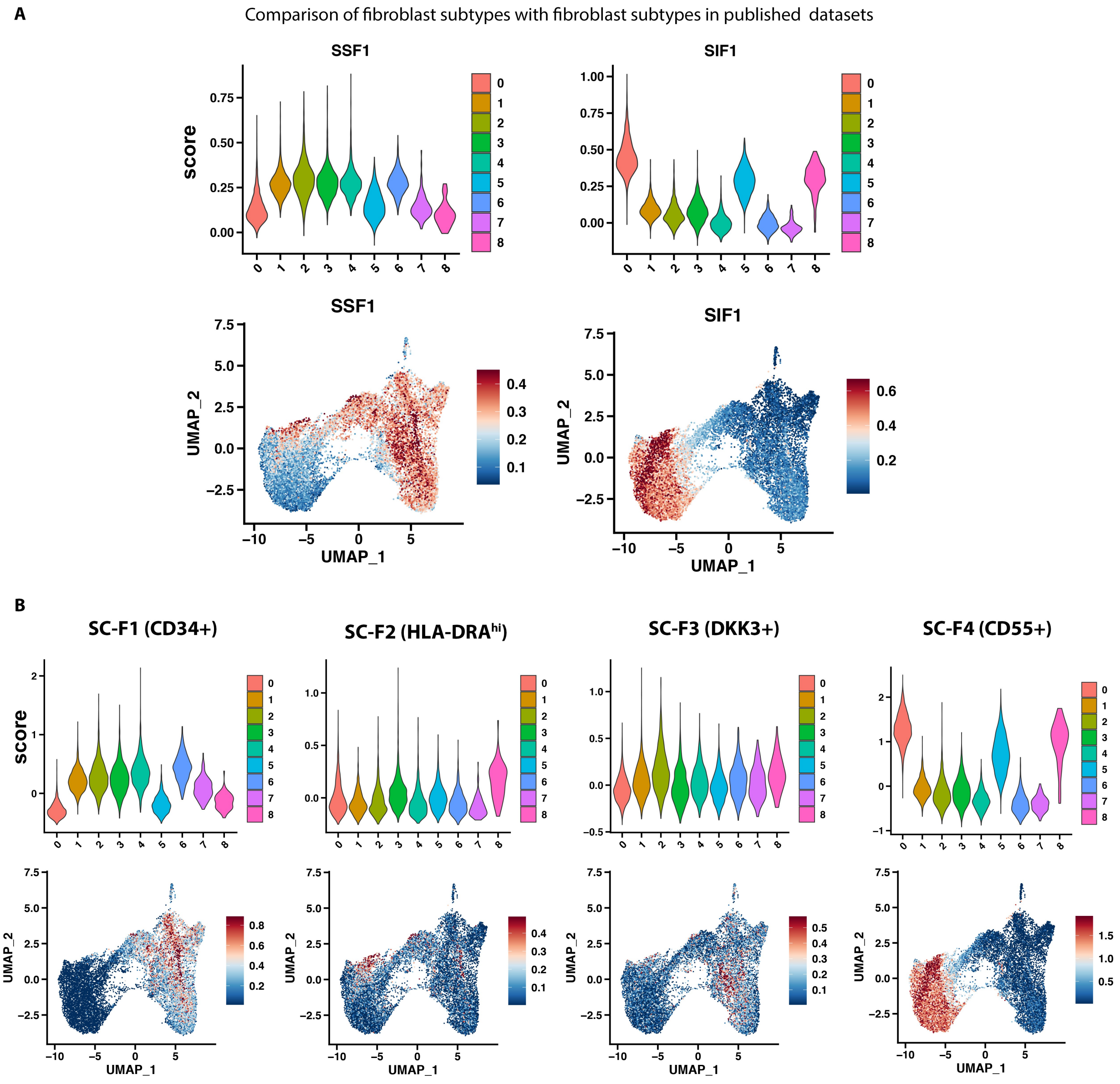

**Fig. S4. Trajectory analysis of fibroblast subclusters.** A) Individual unique fibroblast subcluster trajectories over pseudotime. B) Top third to fifth DEGs for fibroblast subcluster 0-6 showing changes in expression over pseudotime on jitter plots.

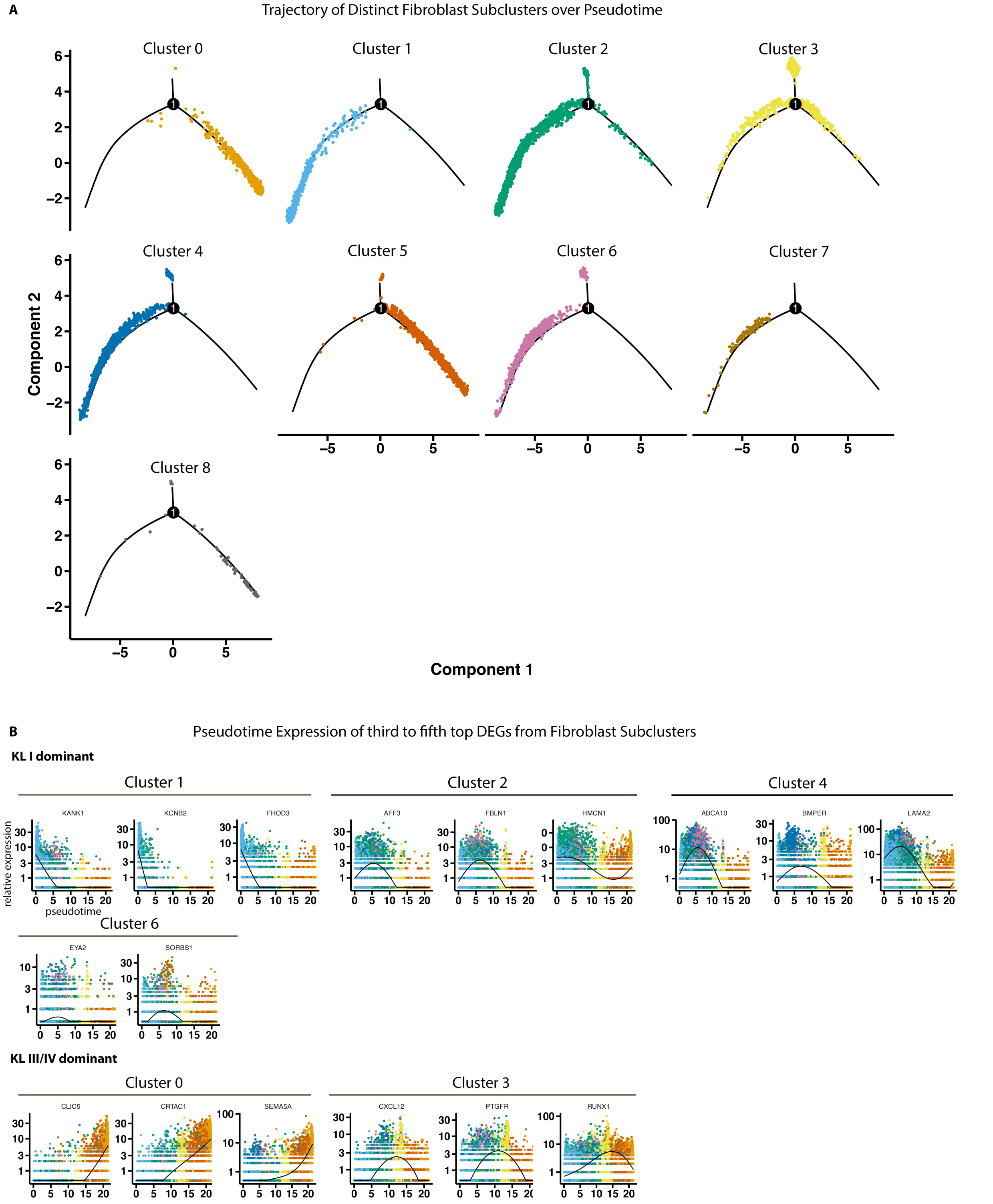

**Fig. S5. CellChat analysis of fibroblast subclusters.** A) CellChat analysis depicting interaction strengths between major cell types. B) Chord diagram showing cell-cell communication between major cell types with fibroblast as the 'receiver' expressing receptors, and all other cell types as 'senders' expressing ligands. C) CellChat analysis depicting interaction strengths between fibroblast subclusters. D) Bubble plot showing communication probability of upregulated and downregulated signaling ligand-receptor pairs in advanced compared to early-stage with fibroblast subcluster 0 as the 'receiver' and fibro subcluster 1 as the 'sender'. E) Bubble plot showing communication probability in up-regulated and down-regulated signaling ligand-receptor pairs in advanced as compared to early with fibroblast subcluster 1 as the 'receiver' and fibro subcluster 0 as the 'sender'. F) Outgoing signaling patterns associated with each fibroblast subclusters in early and advanced conditions G) Cell-cell communication between fibroblast subcluster 1 as the receiver and ligands from all other fibroblast subclusters as senders.

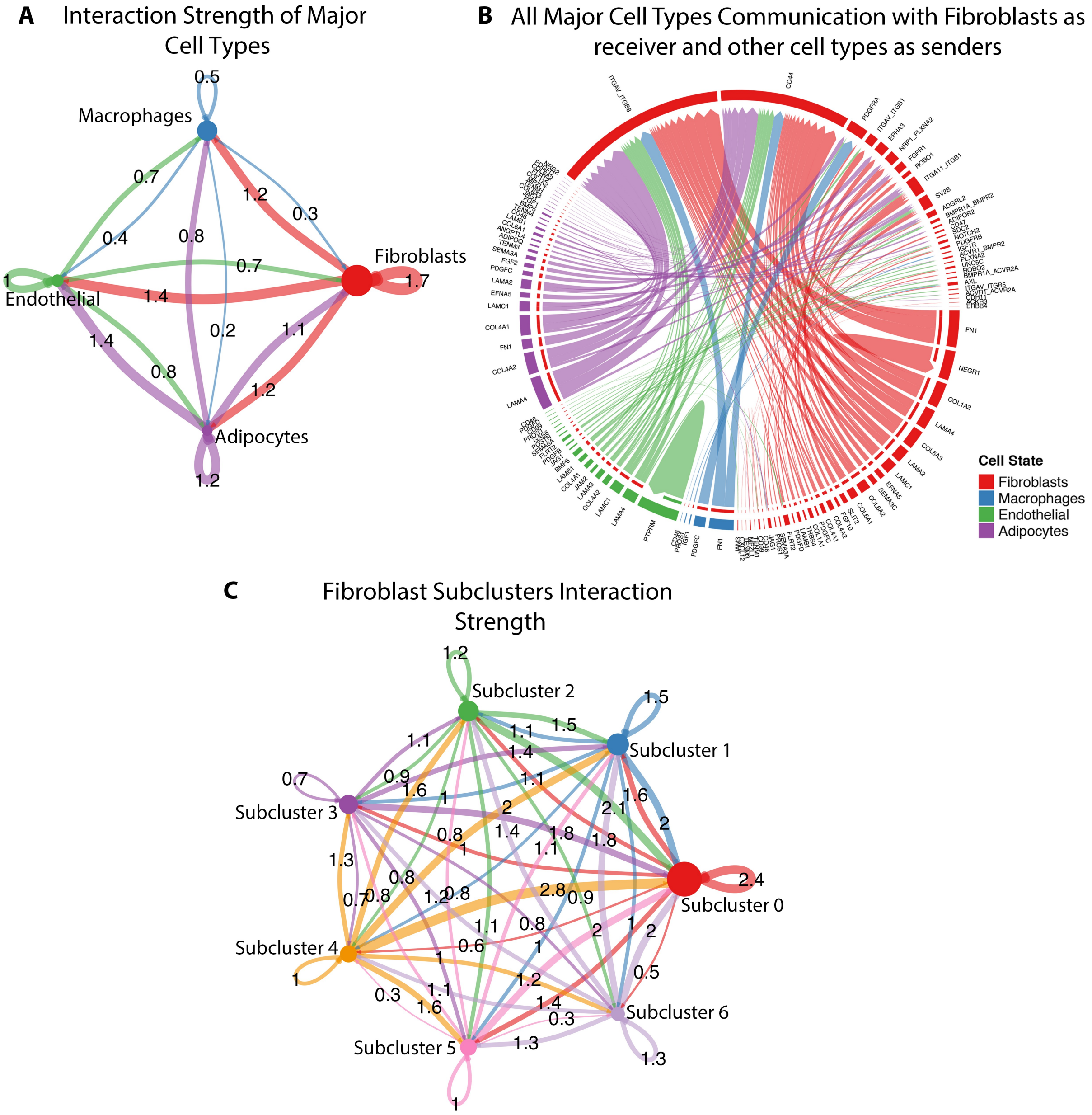

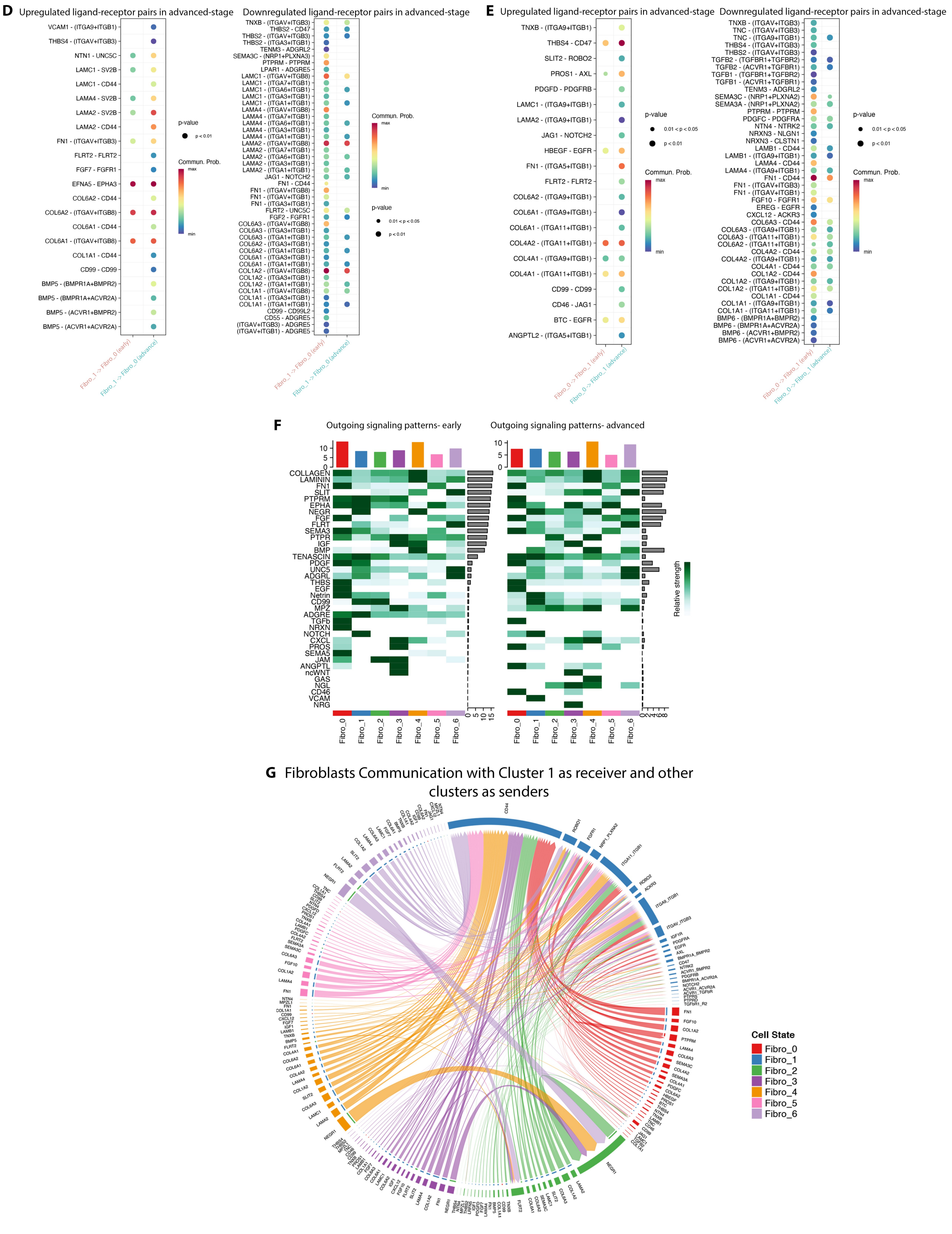
D
Upregulated ligand-receptor pairs in advanced-stage
VCAM1 - (ITGA9+ITGB1)
THBS4 - (ITGAV+ITGB3)
NTN1 - UNC5C
LAMC1 - SV2B
LAMC1 - CD44
LAMA4 - SV2B
LAMA2 - SV2B
LAMA2 - CD44
FN1 - (ITGAV+ITGB3)
FLRT2 - FLRT2
FGF7 - FGFR1
EFNA5 - EPHA3
COL6A2 - CD44
COL6A2 - (ITGAV+ITGB8)
COL6A1 - CD44
COL6A1 - (ITGAV+ITGB8)
COL1A1 - CD44
CD99 - CD99
BMP5 - (BMPR1A+BMPR2)
BMP5 - (BMPR1A+ACVR2A)
BMP5 - (ACVR1+BMPR2)
BMP5 - (ACVR1+ACVR2A)
Fibro_1 -> Fibro_0 (early)
Fibro_1 -> Fibro_0 (advance)
p-value
p < 0.01
Commun. Prob.
max
min
Downregulated ligand-receptor pairs in advanced-stage
TNXB - (ITGAV+ITGB3)
THBS2 - CD47
THBS2 - (ITGAV+ITGB3)
THBS2 - (ITGA3+ITGB1)
TENM3 - ADGRL2
SEMA3C - (NRP1+PLXNA3)
PTPRM - PTPRM
LPAR1 - ADGRE5
LAMC1 - (ITGAV+ITGB8)
LAMC1 - (ITGA7+ITGB1)
LAMC1 - (ITGA6+ITGB1)
LAMC1 - (ITGA3+ITGB1)
LAMC1 - (ITGA1+ITGB1)
LAMA4 - (ITGAV+ITGB8)
LAMA4 - (ITGA6+ITGB1)
LAMA4 - (ITGA7+ITGB1)
LAMA4 - (ITGA3+ITGB1)
LAMA4 - (ITGA1+ITGB1)
LAMA2 - (ITGAV+ITGB8)
LAMA2 - (ITGA7+ITGB1)
LAMA2 - (ITGA6+ITGB1)
LAMA2 - (ITGA3+ITGB1)
LAMA2 - (ITGA1+ITGB1)
JAG1 - NOTCH2
FN1 - CD44
FN1 - (ITGAV+ITGB8)
FN1 - (ITGA1+ITGB1)
FN1 - (ITGA3+ITGB1)
FLRT2 - UNC5C
FGF2 - FGFR1
COL6A3 - (ITGAV+ITGB8)
COL6A3 - (ITGA3+ITGB1)
COL6A3 - (ITGA1+ITGB1)
COL6A2 - (ITGA3+ITGB1)
COL6A2 - (ITGA1+ITGB1)
COL6A1 - (ITGA3+ITGB1)
COL6A1 - (ITGA1+ITGB1)
COL1A2 - (ITGAV+ITGB8)
COL1A2 - (ITGA3+ITGB1)
COL1A2 - (ITGA1+ITGB1)
COL1A1 - (ITGAV+ITGB8)
COL1A1 - (ITGA3+ITGB1)
COL1A1 - (ITGA1+ITGB1)
CD99 - CD99L2
CD55 - ADGRE5
(ITGAV+ITGB3) - ADGRE5
(ITGAV+ITGB1) - ADGRE5
Fibro_1 -> Fibro_0 (early)
Fibro_1 -> Fibro_0 (advance)
Commun. Prob.
max
min
p-value
0.01 < p < 0.05
p < 0.01
E
Upregulated ligand-receptor pairs in advanced-stage
TNXB - (ITGA9+ITGB1)
THBS4 - CD47
SLIT2 - ROBO2
PROS1 - AXL
PDGFD - PDGFRB
LAMC1 - (ITGA9+ITGB1)
LAMA2 - (ITGA9+ITGB1)
JAG1 - NOTCH2
HBEGF - EGFR
FN1 - (ITGA5+ITGB1)
FLRT2 - FLRT2
COL6A2 - (ITGA9+ITGB1)
COL6A1 - (ITGA9+ITGB1)
COL6A1 - (ITGA11+ITGB1)
COL4A2 - (ITGA11+ITGB1)
COL4A1 - (ITGA9+ITGB1)
COL4A1 - (ITGA11+ITGB1)
CD99 - CD99
CD46 - JAG1
BTC - EGFR
ANGPTL2 - (ITGA5+ITGB1)
Fibro_0 -> Fibro_1 (early)
Fibro_0 -> Fibro_1 (advance)
p-value
0.01 < p < 0.05
p < 0.01
Commun. Prob.
max
min
Downregulated ligand-receptor pairs in advanced-stage
TNXB - (ITGAV+ITGB3)
TNC - (ITGAV+ITGB3)
TNC - (ITGA9+ITGB1)
THBS4 - (ITGAV+ITGB3)
THBS2 - (ITGAV+ITGB3)
TGFB2 - (TGFBR1+TGFBR2)
TGFB2 - (ACVR1+TGFBR1)
TGFB1 - (TGFBR1+TGFBR2)
TGFB1 - (ACVR1+TGFBR1)
TENM3 - ADGRL2
SEMA3C - (NRP1+PLXNA2)
SEMA3A - (NRP1+PLXNA2)
PTPRM - PTPRM
PDGFC - PDGFRA
NTN4 - NTRK2
NRXN3 - NLGN1
NRXN3 - CLSTN1
LAMB1 - CD44
LAMB1 - (ITGA9+ITGB1)
LAMA4 - CD44
LAMA4 - (ITGA9+ITGB1)
FN1 - CD44
FN1 - (ITGAV+ITGB3)
FN1 - (ITGAV+ITGB1)
FGF10 - FGFR1
EREG - EGFR
CXCL12 - ACKR3
COL6A3 - CD44
COL6A3 - (ITGA9+ITGB1)
COL6A3 - (ITGA11+ITGB1)
COL6A2 - (ITGA11+ITGB1)
COL4A2 - CD44
COL4A2 - (ITGA9+ITGB1)
COL4A1 - CD44
COL1A2 - CD44
COL1A2 - (ITGA9+ITGB1)
COL1A2 - (ITGA11+ITGB1)
COL1A1 - CD44
COL1A1 - (ITGA9+ITGB1)
COL1A1 - (ITGA11+ITGB1)
BMP6 - (BMPR1A+BMPR2)
BMP6 - (BMPR1A+ACVR2A)
BMP6 - (ACVR1+BMPR2)
BMP6 - (ACVR1+ACVR2A)
Fibro_0 -> Fibro_1 (early)
Fibro_0 -> Fibro_1 (advance)
p-value
0.01 < p < 0.05
p < 0.01
Commun. Prob.
max
min
F
Outgoing signaling patterns- early
Outgoing signaling patterns- advanced
COLLAGEN
LAMININ
FN1
SLIT
PTPRM
EPHA
NEGR
FGF
FLRT
SEMA3
PTPR
IGF
BMP
TENASCIN
PDGF
UNC5
ADGRL
THBS
EGF
Netrin
CD99
MPZ
ADGRE
TGFb
NRXN
NOTCH
CXCL
PROS
SEMA5
JAM
ANGPTL
ncWNT
GAS
NGL
CD46
VCAM
NRG
Fibro_0
Fibro_1
Fibro_2
Fibro_3
Fibro_4
Fibro_5
Fibro_6
Relative strength
G
Fibroblasts Communication with Cluster 1 as receiver and other clusters as senders
Cell State
Fibro_0
Fibro_1
Fibro_2
Fibro_3
Fibro_4
Fibro_5
Fibro_6

**Fig. S6. Flow Cytometry on healthy, KLI and KLIII/IV graded radiographic knee OA synovium.** A & B) Total of all cells (A), and immune, endothelia, mural cells and fibroblasts (B) sorted from healthy, KLI and KLIII/IV radiographic KOA synovial samples. C) Gating strategy to sort out fibroblasts. D & E) Expression of FAP+ and CD55+ fibroblasts. Data was statistically analyzed using one way ANOVA corrected for multiple testing using the Kruskal-Wallis test. Adjusted P<0.05 was considered significant. Not significant (n.s.) difference; *, P≤0.05; **, P≤0.01. F) Expression of select fibroblast metaclusters in the fibroblast population. G) Proportion of fibroblast metaclusters in healthy, KLI and KLIII/IV radiographic KOA synovial tissue.

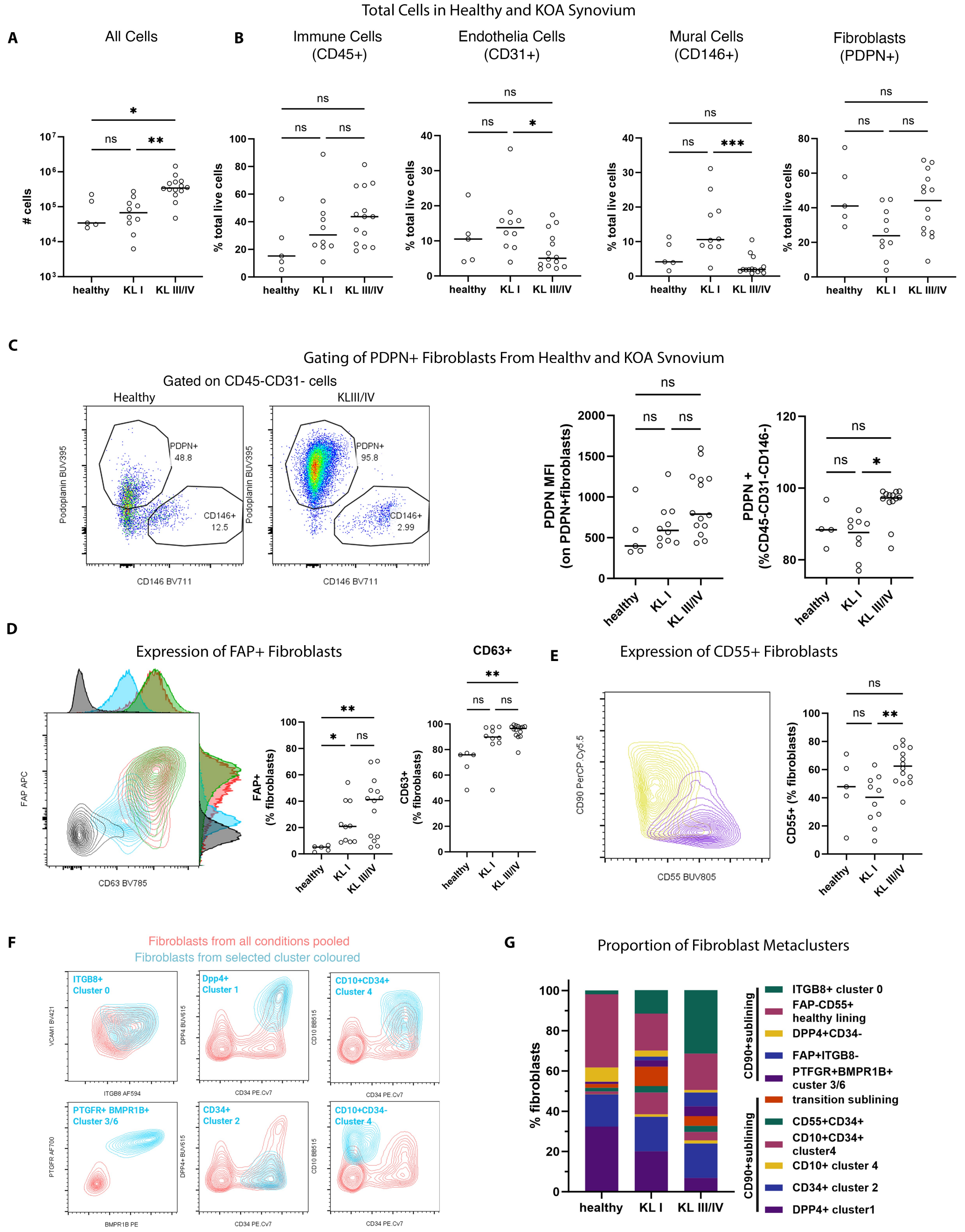

**Fig. S7. Single nuclei RNA-sequencing of mouse synovium at 2- and 10-weeks post-DMM surgery, and control samples.** A) Single nuclei RNA sequencing of mouse synovium resolving multiple cell types. B) Proportion of each cell type in synovia from 2- and 10-weeks post-DMM surgery time points, and control synovia. C) Canonical markers used to identify each cell type. D) Top ten DEGs expressed in each fibroblast subcluster identified in mouse synovium. E) Expression of analogous mouse fibroblast subclusters to human fibroblast subclusters.

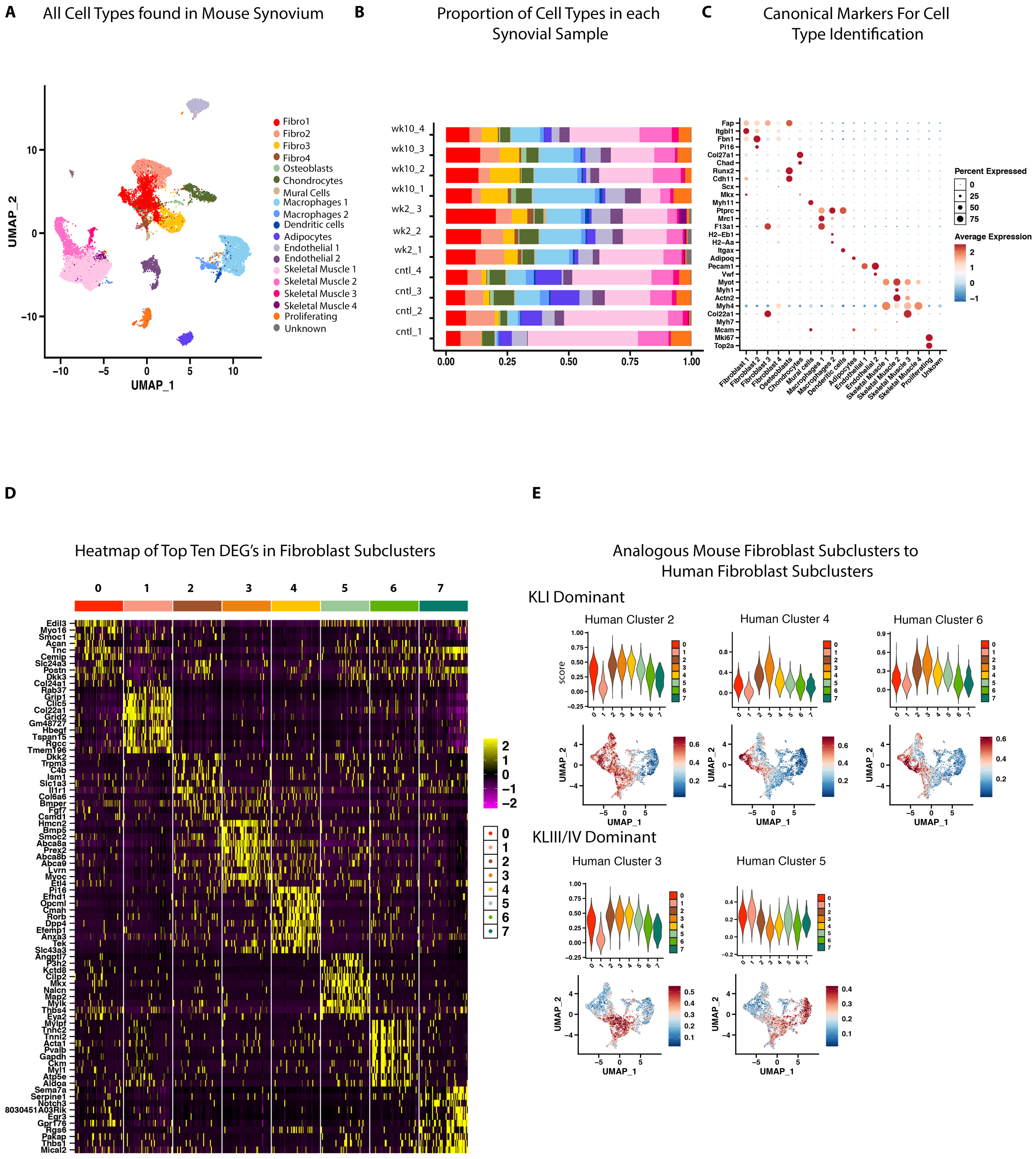

**Fig. S8. Transcription factor-gene-pathway network analysis of all human fibroblast subclusters.** Pathway enrichment analysis performed using PathDIP on all fibroblast subclusters from single nuclei RNA-sequencing analysis of n=5 KLI and n=4 KLIII/IV graded radiographic OA synovial samples. Transcription factor (TF) enrichment analysis on DEGs from each fibroblast subcluster was performed using Catrin. Only pathways and biological processes overlapping between fibroblast subcluster 1 and 0 are shown.

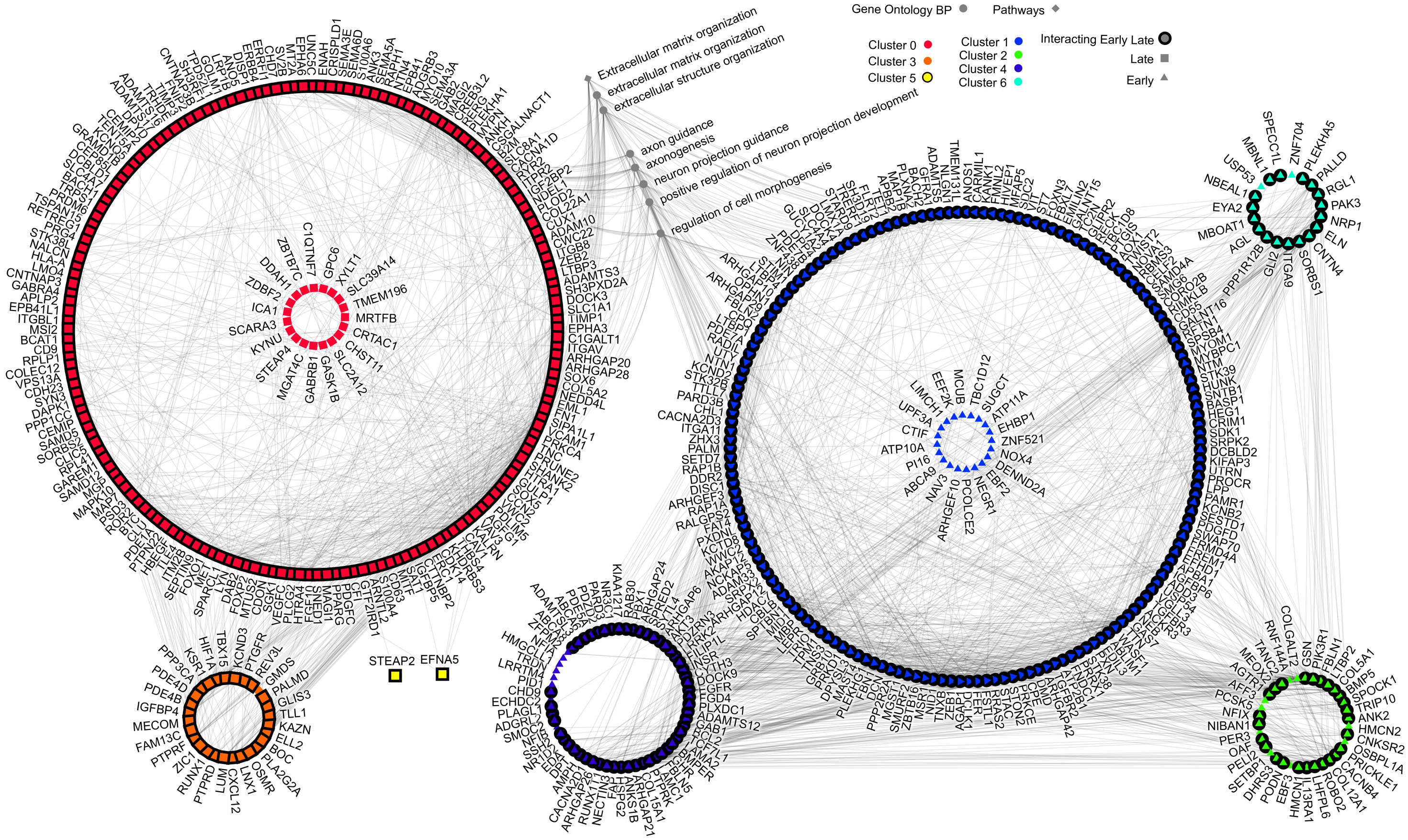

**Fig. S9. Matrisome gene expression and *BHLHE40* siRNA knockdown in FLS extracted from synovium of patients with KLIII/IV radiographic KOA.** A) Proportion of matrisome-annotated genes expressed by fibroblasts that are upregulated, downregulated or not differentially-expressed (DE) in advanced- compared to early-stage KOA synovia. B) Downregulated and C) upregulated genes from the Nanostring fibrosis panel showing significant changes in control versus *BHLHE40* siRNA knockdown treatments. Statistical significance was determined using paired parametric T-tests. *, P ≤ 0.05; **, P ≤ 0.01; *** P ≤ 0.001.

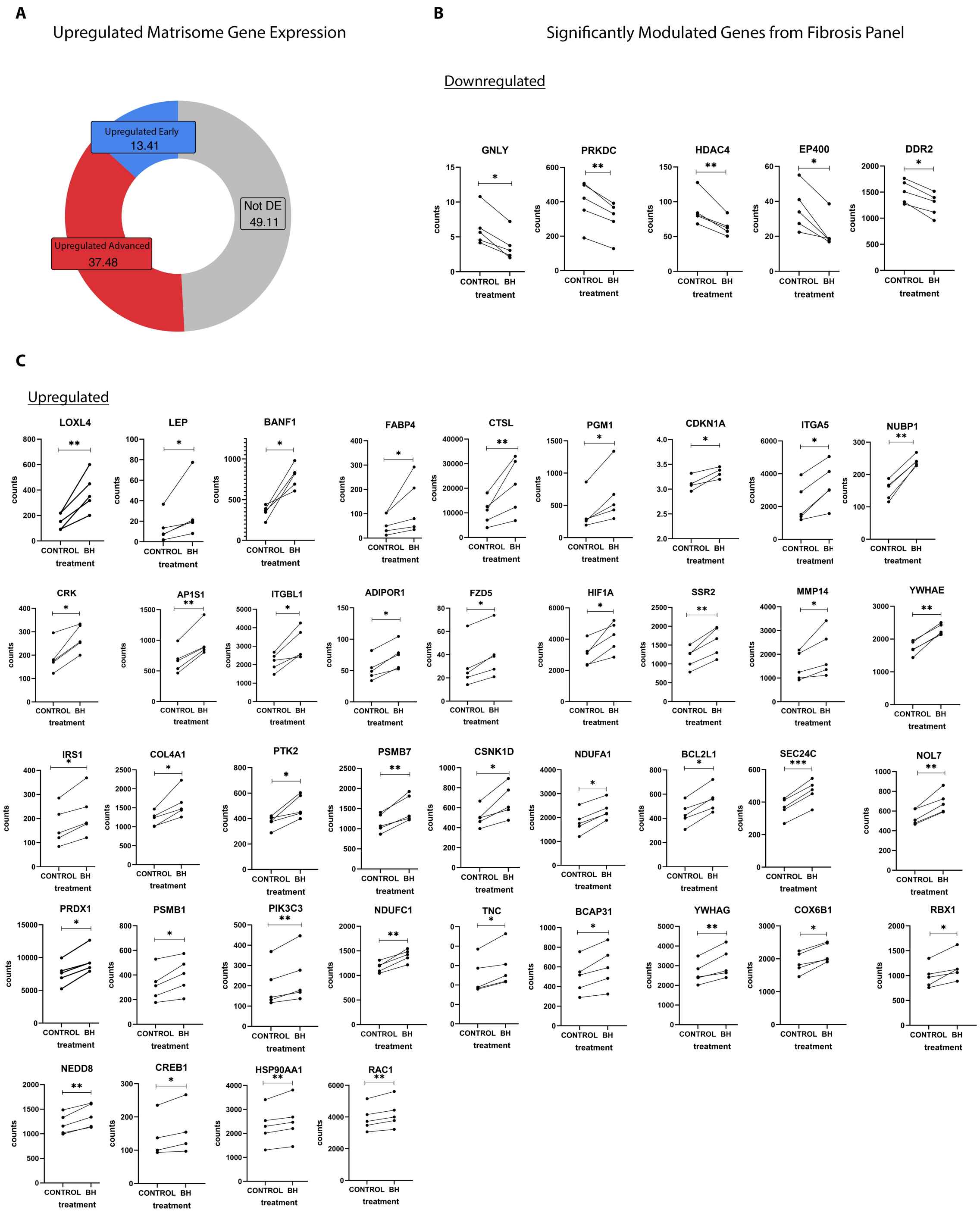

**Fig. S10. BHLHE40 lentiviral transfection at different MOI concentrations.** Advanced-stage fibroblasts (KL3/4) infected with BHLHE40 lentivirus at 0, 25, 50 and 100 MOI.

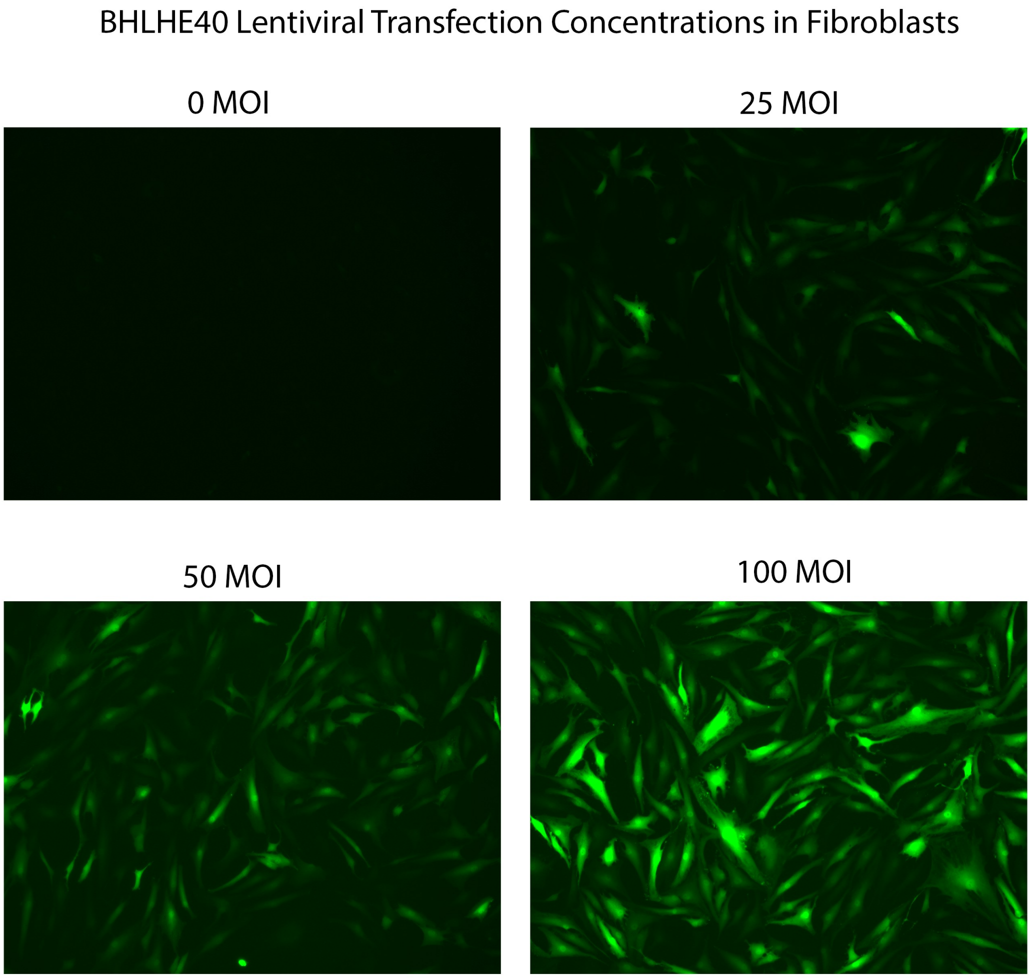

**Table S1. Demographic and anthropometric details of subjects.** Synovial samples from early- and advanced-stage radiographic knee OA patients were collected from arthroscopic surgery (early-stage) or discarded tissue from total knee arthroplasty surgeries (advanced-stage), and normal/healthy knee synovium samples were obtained from organ donors. Collected tissues were utilized in bulk RNA sequencing experiments (KL I, n=6; KL III/IV, n=8) (A), single nuclei RNA sequencing experiments (KL I, n=5; KL III/IV, n=4) (B), flow cytometry (healthy control, n=5; KL I, n=10; KL III/IV, n=12) (C), and siRNA knockdown in vitro experiments (KL III/IV; n=5) (D). All patients described in A-E are independent and never duplicated for independent experiments. The following cohorts are associated with the listed figures: cohort (A); Fig. 1C, 1D, 6C-D, fig. S9A cohort (B); Fig. 2, 3, fig. S1, S4, S5, S8 cohort(C); Fig. 4, fig. S6 cohort (D); Fig. 6F-I, fig. S9B-C, cohort (E); Fig. 6J, cohort (F); Fig. 8b.

A) Bulk RNA sequencing

| Stage | ID | Age | Sex | BMI |
|---|---|---|---|---|
| KL I | 1 | 54 | Male | 38.4 |
| | 2 | 44 | Male | 25.7 |
| | 3 | 37 | Female | 25.5 |
| | 4 | 58 | Male | 27.4 |
| | 5 | 65 | Female | 18.4 |
| | 6 | 35 | Male | 34.7 |
| KL III/IV | 7 | 61 | Male | 27.35 |
| | 8 | 62 | Male | 22.86 |
| | 9 | 60 | Female | 31.28 |
| | 10 | 57 | Female | 27.34 |
| | 11 | 55 | Male | 34.25 |
| | 12 | 58 | Female | 32.61 |
| | 13 | 56 | Female | 24.79 |
| | 14 | 57 | Male | 25.62 |

B) Single Nuclei RNA sequencing

| Stage | ID | Age | Sex | BMI |
|---|---|---|---|---|
| KL I | 1 | 66 | Male | 24.4 |
| | 2 | 52 | Female | 18.8 |
| | 3 | 48 | Female | 29.3 |
| | 4 | 56 | Female | 30.0 |
| | 5 | 57 | Male | 24.8 |
| KL III/IV | 6 | 56 | Female | 21.7 |
| | 7 | 59 | Female | 28.3 |
| | 8 | 61 | Female | 34.6 |
| | 9 | 61 | Male | 21.0 |

C) Flow Cytometry

| Stage | ID | Age | Sex | BMI |
|---|---|---|---|---|
| **Healthy control** | 1 | 72 | F | N/A |
| | 2 | 65 | F | N/A |
| | 3 | 36 | M | N/A |
| | 4 | 39 | F | N/A |
| | 5 | 69 | M | N/A |
| **KL I** | 6 | 62 | F | 31.2 |
| | 7 | 32 | M | 33.2 |
| | 8 | 58 | M | 28.9 |
| | 9 | 46 | M | 24.3 |
| | 10 | 50 | M | 26.3 |
| | 11 | 36 | M | 21.9 |
| | 12 | 60 | F | 34.8 |
| | 13 | 45 | M | 26.3 |
| | 14 | 47 | M | 34.5 |
| | 15 | 32 | M | 28.5 |
| **KL III/IV** | 16 | 61 | F | 29.1 |
| | 17 | 53 | M | 33.0 |
| | 18 | 79 | M | 40.1 |
| | 19 | 61 | F | 50.2 |
| | 20 | 64 | M | 28.5 |
| | 21 | 76 | F | 26.7 |
| | 22 | 77 | F | 24.2 |
| | 23 | 61 | M | 28.1 |
| | 24 | 74 | M | 34.6 |
| | 25 | 60 | M | 27.8 |
| | 26 | 78 | F | 19.6 |
| | 27 | 62 | M | 26.1 |

D) In Vitro SiRNA knockdown experiment

| Stage | ID | Age | Sex | BMI |
|---|---|---|---|---|
| **KL III/IV** | 1 | 72 | F | 29.1 |
| | 2 | 79 | F | 44.6 |
| | 3 | 70 | F | 22.6 |
| | 4 | 57 | F | 25.2 |
| | 5 | 63 | F | 28 |

E) In Vitro SiRNA knockout IF staining

| Stage | ID | Age | Sex | BMI | IF stain |
|---|---|---|---|---|---|
| **KL III/IV** | 1 | 65 | M | 25.2 | RP/ αSMA |
| | 2 | 57 | F | 25.2 | RP/ αSMA |
| | 3 | 70 | M | 24.4 | RP/ αSMA |

| | | | | | |
|---|---|---|---|---|---|
| | 4 | 72 | M | 22.0 | RP/ αSMA |

F) In Vitro Lentivirus overexpression IF staining

| Stage | ID | Age | Sex | BMI | IF stain |
|---|---|---|---|---|---|
| **KL III/IV** | 1 | 65 | M | 25.2 | RP/ αSMA |
| | 2 | 72 | M | 22.0 | RP/ αSMA |
| | 3 | 67 | M | 25.3 | RP/ αSMA |
| | 4 | 70 | M | 24.4 | RP/ αSMA |

**Table S2: Top 50 upregulated genes identified in bulk RNA sequencing of KOA synovia, from a total of 8,904 differentially-expressed genes.** Differentially-expressed genes (DEGs) in synovia from advanced-stage (KLIII/IV, n=8) versus early-stage (KLI, n=6) radiographic knee OA using bulk RNA sequencing. DEGs are organized by decreasing average $log_2$-fold change (FC) and filtered by adjusted p-value $< 0.05$ and $log_2FC \geq 0.5$. Full List available upon request. Raw data deposited in GEO Accession Number: GSE281825.

| gene | baseMean | log2FoldChange | pvalue | padj |
|---|---|---|---|---|
| CPZ | 130.995899 | 9.877889655 | 2.08E-24 | 8.07E-23 |
| LY75-CD302 | 113.675036 | 9.673074438 | 2.84E-19 | 6.97E-18 |
| AOC1 | 107.446259 | 9.590445975 | 2.86E-19 | 7.03E-18 |
| STMN2 | 70.4943346 | 8.98201247 | 1.06E-19 | 2.75E-18 |
| AMTN | 61.5740375 | 8.789255678 | 3.14E-11 | 3.18E-10 |
| PSD | 53.3489358 | 8.584277472 | 1.14E-18 | 2.63E-17 |
| IGKV1-33 | 43.3585002 | 8.275669757 | 2.67E-14 | 3.86E-13 |
| GIMAP1-GIMAP5 | 38.724848 | 8.121903902 | 2.20E-11 | 2.27E-10 |
| IGKV1-6 | 38.6581374 | 8.112750752 | 1.04E-11 | 1.13E-10 |
| C4orf48 | 37.407229 | 8.069825278 | 9.88E-21 | 2.83E-19 |
| SEZ6L2 | 35.8302458 | 8.012098992 | 4.66E-14 | 6.57E-13 |
| IGKV1D-33 | 35.6310418 | 7.99167985 | 1.61E-14 | 2.37E-13 |
| AC083855.2 | 34.4146617 | 7.954330017 | 1.07E-19 | 2.76E-18 |
| IGKV3-11 | 87.8875206 | 7.899053646 | 7.42E-13 | 9.26E-12 |
| LINC02341 | 32.9025006 | 7.886712649 | 2.07E-16 | 3.76E-15 |
| KCNJ6 | 32.7584704 | 7.879477587 | 3.68E-13 | 4.73E-12 |
| BATF | 31.3488053 | 7.82766586 | 7.63E-19 | 1.79E-17 |
| CLEC4G | 28.9236109 | 7.697877368 | 3.07E-07 | 1.77E-06 |
| MT1G | 27.7129245 | 7.621965097 | 8.70E-16 | 1.48E-14 |
| IGKV1-39 | 68.9969604 | 7.470072893 | 4.08E-12 | 4.61E-11 |
| C5orf38 | 22.2677511 | 7.313113491 | 1.87E-15 | 3.07E-14 |
| SPATA20P1 | 21.9616064 | 7.301325556 | 6.67E-13 | 8.36E-12 |
| AC063960.2 | 21.605137 | 7.283836749 | 1.04E-14 | 1.57E-13 |
| KNDC1 | 20.4667395 | 7.205517811 | 1.35E-13 | 1.82E-12 |
| ASGR1 | 20.0898584 | 7.171351626 | 1.94E-15 | 3.16E-14 |
| DHDH | 20.0367102 | 7.168896227 | 1.10E-12 | 1.33E-11 |
| NRTN | 19.686165 | 7.141587675 | 2.52E-14 | 3.65E-13 |
| IGKV1-17 | 65.3603904 | 7.119708311 | 1.21E-10 | 1.14E-09 |
| CHST13 | 19.2700585 | 7.119090321 | 5.07E-14 | 7.11E-13 |
| IGKV3D-15 | 19.0545094 | 7.091828759 | 1.91E-12 | 2.26E-11 |
| IGKV3-20 | 136.329549 | 7.083934858 | 7.54E-16 | 1.29E-14 |
| LINC00603 | 18.4926775 | 7.055021535 | 9.17E-15 | 1.39E-13 |
| IGKV4-1 | 100.461013 | 7.023067534 | 2.89E-14 | 4.15E-13 |
| MYBL2 | 17.9491427 | 7.01579263 | 6.75E-15 | 1.04E-13 |
| IGHV4-39 | 18.0711966 | 7.011129308 | 9.91E-10 | 8.23E-09 |
| CROCC2 | 17.5491058 | 6.981499725 | 2.44E-09 | 1.93E-08 |
| TRIM17 | 16.6921259 | 6.904372295 | 5.06E-14 | 7.10E-13 |
| SIGLEC15 | 16.5123577 | 6.903050452 | 3.54E-14 | 5.05E-13 |
| IGLV3-19 | 16.466106 | 6.876092678 | 4.72E-11 | 4.68E-10 |
| AC087235.2 | 16.0713714 | 6.85516269 | 2.82E-08 | 1.90E-07 |
| CD79A | 16.270992 | 6.851938329 | 5.84E-11 | 5.71E-10 |
| IGKV1-9 | 21.5327347 | 6.773236016 | 3.50E-12 | 3.99E-11 |
| FAM187A | 15.1043337 | 6.758481704 | 3.06E-06 | 1.50E-05 |
| IGLV1-47 | 20.8544524 | 6.737635469 | 1.43E-08 | 1.01E-07 |
| UBE2C | 14.0985277 | 6.669610001 | 2.15E-12 | 2.53E-11 |
| TAC1 | 108.898532 | 6.574142435 | 1.59E-07 | 9.57E-07 |
| AC009630.1 | 13.0202881 | 6.55722184 | 2.85E-12 | 3.29E-11 |
| LINC02224 | 12.9154265 | 6.545450744 | 9.73E-12 | 1.06E-10 |
| CDT1 | 12.7505358 | 6.520196523 | 1.56E-13 | 2.10E-12 |
| CYP4F11 | 12.6198553 | 6.506980412 | 3.17E-10 | 2.81E-09 |

**Table S3: Marker genes for the top four major cell types identified in single nuclei RNA sequencing.** Top 20 differentially-expressed genes (DEGs) for major cell types in human KOA synovia subjected to single nuclei RNA sequencing (KLI; n=5, KLIII/IV; n=4). DEGs are organized by decreasing average $log_2$-fold change (FC) and filtered by adjusted p-value < 0.05, $log_2FC \geq 0.5$ and min.pct=0.25. Full list available upon request. Raw data deposited in GEO Accession Number: GSE281826.

| gene | p_val | avg_log2FC | pct.1 | pct.2 | p_val_adj | cluster |
|---|---|---|---|---|---|---|
| COL22A1 | 0 | 6.480733 | 0.612 | 0.017 | 0 | Fibro1 |
| AP002370.2 | 0 | 5.7160871 | 0.688 | 0.027 | 0 | Fibro1 |
| DIRC3-AS1 | 0 | 5.6005995 | 0.503 | 0.02 | 0 | Fibro1 |
| GABRA4 | 0 | 5.4979383 | 0.71 | 0.027 | 0 | Fibro1 |
| HMGA2 | 0 | 5.2003642 | 0.37 | 0.012 | 0 | Fibro1 |
| CLIC5 | 0 | 5.0975328 | 0.819 | 0.068 | 0 | Fibro1 |
| SHANK2 | 0 | 5.0852716 | 0.578 | 0.024 | 0 | Fibro1 |
| HTRA4 | 0 | 5.048914 | 0.5 | 0.033 | 0 | Fibro1 |
| MTUS2 | 0 | 5.036228 | 0.543 | 0.021 | 0 | Fibro1 |
| LINC01060 | 0 | 4.8426869 | 0.507 | 0.03 | 0 | Fibro1 |
| AC004990.1 | 0 | 4.8111271 | 0.377 | 0.013 | 0 | Fibro1 |
| ITGB8 | 0 | 4.7762716 | 0.995 | 0.225 | 0 | Fibro1 |
| SEMA5A | 0 | 4.7165515 | 0.937 | 0.112 | 0 | Fibro1 |
| GABRB1 | 0 | 4.6875797 | 0.748 | 0.037 | 0 | Fibro1 |
| GOLGA7B | 0 | 4.6592705 | 0.255 | 0.008 | 0 | Fibro1 |
| MYPN | 0 | 4.5997405 | 0.696 | 0.049 | 0 | Fibro1 |
| HBEGF | 0 | 4.5965879 | 0.431 | 0.026 | 0 | Fibro1 |
| PPP4R4 | 0 | 4.4124772 | 0.416 | 0.022 | 0 | Fibro1 |
| AC093801.1 | 0 | 4.4078498 | 0.332 | 0.015 | 0 | Fibro1 |
| ADAMTS16 | 0 | 4.3818099 | 0.611 | 0.043 | 0 | Fibro1 |
| KCNB2 | 0 | 6.3416005 | 0.386 | 0.012 | 0 | Fibro2 |
| KIRREL3 | 0 | 5.322649 | 0.276 | 0.019 | 0 | Fibro2 |
| AC093425.1 | 0 | 5.3191709 | 0.334 | 0.017 | 0 | Fibro2 |
| PXDNL | 0 | 4.937949 | 0.742 | 0.109 | 0 | Fibro2 |
| PAMR1 | 0 | 4.6603835 | 0.426 | 0.023 | 0 | Fibro2 |
| LINC01133 | 0 | 4.427982 | 0.388 | 0.031 | 0 | Fibro2 |
| MFAP5 | 0 | 4.349371 | 0.729 | 0.081 | 0 | Fibro2 |
| KCTD8 | 0 | 4.3092743 | 0.298 | 0.027 | 0 | Fibro2 |
| HUNK | 0 | 4.3039933 | 0.445 | 0.044 | 0 | Fibro2 |
| LINC01915 | 0 | 4.1198716 | 0.334 | 0.045 | 0 | Fibro2 |
| FHOD3 | 0 | 4.1045156 | 0.604 | 0.112 | 0 | Fibro2 |
| DPP4 | 0 | 4.0585763 | 0.492 | 0.036 | 0 | Fibro2 |
| GUCY1A2 | 0 | 3.9630857 | 0.526 | 0.051 | 0 | Fibro2 |
| LMX1A | 0 | 3.7728472 | 0.345 | 0.033 | 0 | Fibro2 |
| ITGA11 | 0 | 3.7609221 | 0.739 | 0.099 | 0 | Fibro2 |
| FNDC1 | 0 | 3.7175379 | 0.408 | 0.041 | 0 | Fibro2 |
| NTM | 0 | 3.6301535 | 0.732 | 0.11 | 0 | Fibro2 |
| ADGRD1 | 0 | 3.5274012 | 0.629 | 0.106 | 0 | Fibro2 |
| FBN1 | 0 | 3.4957033 | 0.923 | 0.494 | 0 | Fibro2 |
| PI16 | 0 | 3.4732873 | 0.253 | 0.025 | 0 | Fibro2 |
| LINC00603 | 0 | 3.7755964 | 0.283 | 0.022 | 0 | Fibro3 |
| SMOC1 | 0 | 3.7729749 | 0.327 | 0.044 | 0 | Fibro3 |
| NTNG1 | 0 | 3.5233428 | 0.305 | 0.037 | 0 | Fibro3 |
| KAZN-AS1 | 0 | 3.5186693 | 0.525 | 0.085 | 0 | Fibro3 |
| KCNH1 | 0 | 3.4807048 | 0.3 | 0.035 | 0 | Fibro3 |
| KCNE4 | 0 | 3.4489258 | 0.305 | 0.034 | 0 | Fibro3 |
| AL357873.1 | 0 | 3.3035368 | 0.556 | 0.115 | 0 | Fibro3 |
| SPON1 | 0 | 3.2493763 | 0.267 | 0.052 | 0 | Fibro3 |
| KAZN | 0 | 3.2010381 | 0.896 | 0.48 | 0 | Fibro3 |
| PTGFR | 0 | 3.1737183 | 0.554 | 0.127 | 0 | Fibro3 |
| BTBD11 | 0 | 3.1345822 | 0.293 | 0.045 | 0 | Fibro3 |
| KIAA1755 | 0 | 3.1078469 | 0.3 | 0.039 | 0 | Fibro3 |
| IGF1 | 0 | 2.9193913 | 0.505 | 0.193 | 0 | Fibro3 |
| WNT5B | 0 | 2.8917853 | 0.388 | 0.104 | 0 | Fibro3 |

| | | | | | | |
|---|---|---|---|---|---|---|
| PDE10A1 | 0 | 2.8527244 | 0.515 | 0.165 | 0 | Fibro3 |
| COL14A11 | 0 | 2.8000221 | 0.757 | 0.245 | 0 | Fibro3 |
| ISM1 | 0 | 2.7649553 | 0.429 | 0.109 | 0 | Fibro3 |
| NID2 | 0 | 2.6937106 | 0.453 | 0.106 | 0 | Fibro3 |
| GRIA3 | 0 | 2.6541814 | 0.339 | 0.067 | 0 | Fibro3 |
| KCND31 | 0 | 2.6461586 | 0.397 | 0.097 | 0 | Fibro3 |
| SCN7A | 0 | 6.5546108 | 0.354 | 0.012 | 0 | Fibro4 |
| LRRTM4 | 0 | 5.6062002 | 0.4 | 0.035 | 0 | Fibro4 |
| AL590807.1 | 0 | 5.3537908 | 0.568 | 0.038 | 0 | Fibro4 |
| LINC00377 | 0 | 5.3326305 | 0.387 | 0.015 | 0 | Fibro4 |
| APOD | 0 | 5.2958298 | 0.35 | 0.017 | 0 | Fibro4 |
| MYOC | 0 | 4.8191683 | 0.29 | 0.012 | 0 | Fibro4 |
| AL445250.1 | 0 | 4.5614492 | 0.345 | 0.031 | 0 | Fibro4 |
| AC062004.1 | 0 | 4.3161222 | 0.332 | 0.025 | 0 | Fibro4 |
| ABCA10 | 0 | 4.3155721 | 0.846 | 0.209 | 0 | Fibro4 |
| MME | 0 | 4.2467783 | 0.367 | 0.06 | 0 | Fibro4 |
| COL15A1 | 0 | 4.0951761 | 0.662 | 0.107 | 0 | Fibro4 |
| LAMA21 | 0 | 3.7748383 | 0.915 | 0.359 | 0 | Fibro4 |
| AMPH | 0 | 3.5244232 | 0.439 | 0.058 | 0 | Fibro4 |
| AL110292.1 | 0 | 3.5091796 | 0.267 | 0.044 | 0 | Fibro4 |
| BMPER1 | 0 | 3.4838746 | 0.483 | 0.106 | 0 | Fibro4 |
| DOK6 | 0 | 3.4338676 | 0.293 | 0.037 | 0 | Fibro4 |
| ABCA81 | 0 | 3.423263 | 0.887 | 0.301 | 0 | Fibro4 |
| F3 | 0 | 3.4024181 | 0.283 | 0.045 | 0 | Fibro4 |
| FMO2 | 5.24E-305 | 3.3771203 | 0.26 | 0.051 | 1.56E-300 | Fibro4 |
| AC024230.1 | 0 | 3.3609459 | 0.266 | 0.041 | 0 | Fibro4 |
| LINC01340 | 4.67E-17 | 2.9686568 | 0.255 | 0.033 | 1.39E-12 | Fibro5 |
| P2RY14 | 9.49E-64 | 2.9316988 | 0.702 | 0.069 | 2.83E-59 | Fibro5 |
| CFAP61 | 1.29E-27 | 2.7304931 | 0.277 | 0.025 | 3.86E-23 | Fibro5 |
| AL162414.1 | 1.65E-56 | 2.7055996 | 0.511 | 0.041 | 4.91E-52 | Fibro5 |
| AC092691.1 | 3.60E-61 | 2.7027493 | 0.66 | 0.062 | 1.07E-56 | Fibro5 |
| DSCAML1 | 6.69E-22 | 2.6442721 | 0.574 | 0.124 | 2.00E-17 | Fibro5 |
| CCDC141 | 1.81E-61 | 2.5989531 | 0.745 | 0.078 | 5.41E-57 | Fibro5 |
| MDGA2 | 7.17E-20 | 2.4333572 | 0.298 | 0.038 | 2.14E-15 | Fibro5 |
| LSAMP | 4.07E-27 | 2.4025752 | 0.66 | 0.134 | 1.21E-22 | Fibro5 |
| RYR1 | 4.54E-33 | 2.3999228 | 0.553 | 0.077 | 1.35E-28 | Fibro5 |
| CCDC136 | 1.72E-13 | 2.2162941 | 0.255 | 0.04 | 5.13E-09 | Fibro5 |
| MED12L | 7.57E-14 | 2.2023171 | 0.277 | 0.046 | 2.26E-09 | Fibro5 |
| GFRA2 | 5.97E-33 | 2.1562113 | 0.468 | 0.055 | 1.78E-28 | Fibro5 |
| SCN9A | 1.28E-41 | 2.1225899 | 0.83 | 0.137 | 3.82E-37 | Fibro5 |
| NCF4 | 8.48E-18 | 2.1188742 | 0.362 | 0.059 | 2.53E-13 | Fibro5 |
| PLEKHG5 | 1.73E-18 | 2.1035988 | 0.426 | 0.077 | 5.15E-14 | Fibro5 |
| HSPA6 | 2.98E-16 | 2.0519422 | 0.362 | 0.064 | 8.91E-12 | Fibro5 |
| CLCN5 | 1.36E-12 | 2.0424706 | 0.553 | 0.167 | 4.07E-08 | Fibro5 |
| LYVE1 | 1.13E-34 | 2.0035412 | 0.702 | 0.114 | 3.38E-30 | Fibro5 |
| DBN11 | 2.98E-15 | 2.0019304 | 0.362 | 0.067 | 8.89E-11 | Fibro5 |
| CXADR | 0 | 5.6038269 | 0.464 | 0.013 | 0 | Mac1 |
| OLR1 | 0 | 5.3765288 | 0.362 | 0.01 | 0 | Mac1 |
| L1TD1 | 0 | 5.3697045 | 0.338 | 0.01 | 0 | Mac1 |
| KCNQ3 | 0 | 5.1847821 | 0.709 | 0.062 | 0 | Mac1 |
| TIMD4 | 0 | 5.1624363 | 0.49 | 0.02 | 0 | Mac1 |
| DLEU7 | 0 | 4.94594 | 0.513 | 0.024 | 0 | Mac1 |
| ABCC3 | 0 | 4.760268 | 0.827 | 0.061 | 0 | Mac1 |
| LINC01094 | 0 | 4.5002395 | 0.503 | 0.029 | 0 | Mac1 |
| RASL10A | 0 | 4.4937885 | 0.414 | 0.018 | 0 | Mac1 |
| SLC11A1 | 0 | 4.4568235 | 0.656 | 0.038 | 0 | Mac1 |
| ST18 | 0 | 4.2187048 | 0.318 | 0.018 | 0 | Mac1 |
| FFAR4 | 0 | 4.1804369 | 0.381 | 0.025 | 0 | Mac1 |
| AC008591.1 | 0 | 4.1276408 | 0.374 | 0.034 | 0 | Mac1 |
| CLEC7A | 0 | 4.1191826 | 0.659 | 0.04 | 0 | Mac1 |
| IGSF21 | 0 | 4.0289304 | 0.468 | 0.027 | 0 | Mac1 |
| LY86 | 0 | 4.0098397 | 0.371 | 0.02 | 0 | Mac1 |
| CDCP1 | 0 | 4.0079327 | 0.307 | 0.018 | 0 | Mac1 |
| LINC02432 | 0 | 3.958251 | 0.266 | 0.017 | 0 | Mac1 |
| FBP1 | 0 | 3.9445127 | 0.325 | 0.024 | 0 | Mac1 |

| | | | | | | |
|---|---|---|---|---|---|---|
| AC138207.5 | 0 | 3.9327731 | 0.272 | 0.015 | 0 | Mac1 |
| F13A11 | 0 | 5.6067105 | 0.975 | 0.084 | 0 | Mac2 |
| CR11 | 0 | 5.2083708 | 0.47 | 0.02 | 0 | Mac2 |
| P2RY141 | 0 | 4.8386761 | 0.6 | 0.031 | 0 | Mac2 |
| AL162414.11 | 0 | 4.8011236 | 0.343 | 0.019 | 0 | Mac2 |
| CCDC1411 | 0 | 4.5856873 | 0.631 | 0.038 | 0 | Mac2 |
| FGF131 | 0 | 4.3372178 | 0.712 | 0.116 | 0 | Mac2 |
| LYVE11 | 0 | 4.2955603 | 0.735 | 0.069 | 0 | Mac2 |
| LINC027981 | 0 | 4.2659603 | 0.583 | 0.069 | 0 | Mac2 |
| SCN9A2 | 0 | 4.1640937 | 0.801 | 0.088 | 0 | Mac2 |
| GFRA21 | 0 | 4.1633005 | 0.382 | 0.032 | 0 | Mac2 |
| RYR11 | 0 | 4.1295904 | 0.533 | 0.043 | 0 | Mac2 |
| PLEKHG51 | 0 | 4.0737037 | 0.428 | 0.052 | 0 | Mac2 |
| IL12RB21 | 0 | 4.0693109 | 0.348 | 0.03 | 0 | Mac2 |
| AC092691.11 | 0 | 4.0339063 | 0.384 | 0.039 | 0 | Mac2 |
| CD163L12 | 0 | 3.9529675 | 0.8 | 0.131 | 0 | Mac2 |
| MRC12 | 0 | 3.6929477 | 0.92 | 0.187 | 0 | Mac2 |
| TNFRSF251 | 0 | 3.6445092 | 0.29 | 0.039 | 0 | Mac2 |
| STAB12 | 0 | 3.58336 | 0.809 | 0.166 | 0 | Mac2 |
| LINC013742 | 0 | 3.5276377 | 0.535 | 0.087 | 0 | Mac2 |
| KHDRBS21 | 0 | 3.4102408 | 0.36 | 0.047 | 0 | Mac2 |
| LINC002782 | 4.86E-32 | 2.9483417 | 0.603 | 0.153 | 1.45E-27 | Mac3 |
| SLC22A151 | 2.69E-36 | 2.7882223 | 0.658 | 0.16 | 8.02E-32 | Mac3 |
| SLC11A11 | 2.05E-38 | 2.6377378 | 0.699 | 0.159 | 6.11E-34 | Mac3 |
| UBXN11 | 3.10E-13 | 2.6119981 | 0.301 | 0.079 | 9.26E-09 | Mac3 |
| FBP11 | 1.78E-13 | 2.5424703 | 0.315 | 0.083 | 5.30E-09 | Mac3 |
| LINC025421 | 2.83E-21 | 2.5237386 | 0.507 | 0.144 | 8.45E-17 | Mac3 |
| FFAR41 | 6.14E-23 | 2.5113227 | 0.425 | 0.095 | 1.83E-18 | Mac3 |
| OLR11 | 3.18E-10 | 2.4986689 | 0.274 | 0.079 | 9.48E-06 | Mac3 |
| FHAD11 | 7.98E-19 | 2.4624054 | 0.466 | 0.128 | 2.38E-14 | Mac3 |
| PFKFB4 | 7.76E-14 | 2.4601603 | 0.26 | 0.057 | 2.32E-09 | Mac3 |
| C3AR11 | 1.39E-12 | 2.4414851 | 0.288 | 0.076 | 4.14E-08 | Mac3 |
| ABCC31 | 5.39E-42 | 2.4228594 | 0.822 | 0.211 | 1.61E-37 | Mac3 |
| MPP11 | 8.18E-16 | 2.395612 | 0.397 | 0.108 | 2.44E-11 | Mac3 |
| FGR1 | 4.77E-14 | 2.3830696 | 0.301 | 0.074 | 1.42E-09 | Mac3 |
| HMOX11 | 2.09E-10 | 2.3675013 | 0.301 | 0.091 | 6.24E-06 | Mac3 |
| SH3TC11 | 2.05E-09 | 2.357133 | 0.274 | 0.085 | 6.11E-05 | Mac3 |
| KCNMA11 | 8.92E-36 | 2.3393042 | 0.932 | 0.414 | 2.66E-31 | Mac3 |
| L1TD11 | 1.95E-12 | 2.3352763 | 0.288 | 0.075 | 5.81E-08 | Mac3 |
| TIMD41 | 4.32E-15 | 2.2920054 | 0.397 | 0.113 | 1.29E-10 | Mac3 |
| MARCO3 | 1.70E-35 | 2.2632898 | 0.808 | 0.217 | 5.08E-31 | Mac3 |
| CADM3-AS1 | 0 | 8.2862405 | 0.394 | 0.002 | 0 | Endo1 |
| RAB3C | 0 | 8.1724416 | 0.368 | 0.004 | 0 | Endo1 |
| ACKR1 | 0 | 7.2604647 | 0.26 | 0.003 | 0 | Endo1 |
| CCL14 | 0 | 6.0742267 | 0.282 | 0.005 | 0 | Endo1 |
| PLCXD3 | 0 | 5.8427906 | 0.479 | 0.019 | 0 | Endo1 |
| POSTN | 0 | 5.702156 | 0.58 | 0.014 | 0 | Endo1 |
| AL589693.1 | 0 | 5.5214521 | 0.783 | 0.034 | 0 | Endo1 |
| TPO | 0 | 5.4730237 | 0.471 | 0.017 | 0 | Endo1 |
| AC105450.1 | 0 | 5.3761997 | 0.293 | 0.012 | 0 | Endo1 |
| RAMP3 | 0 | 5.3407758 | 0.396 | 0.015 | 0 | Endo1 |
| SLCO2A1 | 0 | 5.3158811 | 0.458 | 0.014 | 0 | Endo1 |
| FRMPD4 | 0 | 5.2777431 | 0.299 | 0.019 | 0 | Endo1 |
| AIM2 | 0 | 5.0790618 | 0.266 | 0.016 | 0 | Endo1 |
| CCDC68 | 0 | 5.0766769 | 0.263 | 0.01 | 0 | Endo1 |
| MYRIP | 0 | 5.0304183 | 0.879 | 0.121 | 0 | Endo1 |
| AC097459.1 | 0 | 5.0196603 | 0.271 | 0.011 | 0 | Endo1 |
| RYR3 | 0 | 5.0156883 | 0.497 | 0.038 | 0 | Endo1 |
| ELOVL7 | 0 | 4.9886364 | 0.288 | 0.015 | 0 | Endo1 |
| DIPK2B | 0 | 4.9657748 | 0.702 | 0.032 | 0 | Endo1 |
| DOC2B | 0 | 4.9514891 | 0.264 | 0.011 | 0 | Endo1 |
| BTNL9 | 0 | 8.376537 | 0.741 | 0.017 | 0 | Endo2 |
| RBP7 | 0 | 6.5675822 | 0.276 | 0.008 | 0 | Endo2 |
| NOTCH4 | 0 | 5.9570516 | 0.678 | 0.018 | 0 | Endo2 |
| ADGRF5 | 0 | 5.6524445 | 0.744 | 0.024 | 0 | Endo2 |

| | | | | | | |
|---|---|---|---|---|---|---|
| SPAAR | 0 | 5.5887813 | 0.407 | 0.016 | 0 | Endo2 |
| FLT11 | 0 | 5.5390137 | 0.793 | 0.04 | 0 | Endo2 |
| PLVAP1 | 0 | 5.4646601 | 0.433 | 0.014 | 0 | Endo2 |
| MCF2L1 | 0 | 5.4037485 | 0.636 | 0.036 | 0 | Endo2 |
| KDR | 0 | 5.3918505 | 0.461 | 0.017 | 0 | Endo2 |
| DACH11 | 0 | 5.366186 | 0.771 | 0.042 | 0 | Endo2 |
| PKD1L1 | 0 | 5.3516815 | 0.436 | 0.025 | 0 | Endo2 |
| ZNF3661 | 0 | 5.2319992 | 0.502 | 0.02 | 0 | Endo2 |
| ABLIM32 | 0 | 4.9840774 | 0.702 | 0.128 | 0 | Endo2 |
| SHANK31 | 0 | 4.9383876 | 0.845 | 0.047 | 0 | Endo2 |
| NR5A21 | 0 | 4.8848009 | 0.591 | 0.023 | 0 | Endo2 |
| ARHGEF151 | 0 | 4.8292635 | 0.364 | 0.016 | 0 | Endo2 |
| CDH51 | 0 | 4.7921251 | 0.672 | 0.034 | 0 | Endo2 |
| NEURL1B | 0 | 4.7868361 | 0.466 | 0.025 | 0 | Endo2 |
| RASGRF22 | 0 | 4.7777469 | 0.843 | 0.07 | 0 | Endo2 |
| PIK3R31 | 0 | 4.7368003 | 0.645 | 0.1 | 0 | Endo2 |
| HTR4 | 0 | 9.3910177 | 0.6 | 0.003 | 0 | Endo3 |
| LINC00841 | 0 | 8.4095947 | 0.419 | 0.002 | 0 | Endo3 |
| AL445259.1 | 0 | 7.8656818 | 0.392 | 0.004 | 0 | Endo3 |
| GJA5 | 0 | 7.8026887 | 0.573 | 0.005 | 0 | Endo3 |
| PCDH11Y | 0 | 7.1419152 | 0.319 | 0.007 | 0 | Endo3 |
| PCDH11X | 0 | 7.1028752 | 0.449 | 0.008 | 0 | Endo3 |
| DHH | 0 | 7.0219454 | 0.278 | 0.002 | 0 | Endo3 |
| AC010737.1 | 0 | 6.9538885 | 0.405 | 0.007 | 0 | Endo3 |
| AC008514.1 | 0 | 6.8370299 | 0.332 | 0.003 | 0 | Endo3 |
| LINC00840 | 0 | 6.7612414 | 0.524 | 0.009 | 0 | Endo3 |
| NKAIN2 | 0 | 6.6926368 | 0.897 | 0.093 | 0 | Endo3 |
| WIPF3 | 0 | 6.2576346 | 0.457 | 0.008 | 0 | Endo3 |
| AJ006995.1 | 0 | 6.2124463 | 0.281 | 0.004 | 0 | Endo3 |
| LINC016952 | 0 | 6.1887963 | 0.816 | 0.03 | 0 | Endo3 |
| SV2C | 0 | 6.1552526 | 0.362 | 0.005 | 0 | Endo3 |
| SH3RF2 | 0 | 6.1037146 | 0.697 | 0.026 | 0 | Endo3 |
| IGFBP3 | 0 | 5.8330292 | 0.522 | 0.02 | 0 | Endo3 |
| TGM2 | 0 | 5.661043 | 0.354 | 0.007 | 0 | Endo3 |
| NOS1AP | 0 | 5.6485338 | 0.784 | 0.029 | 0 | Endo3 |
| ATP2A31 | 0 | 5.6287451 | 0.695 | 0.023 | 0 | Endo3 |
| CCL21 | 0 | 9.3181187 | 0.402 | 0.002 | 0 | Endo4 |
| AL121974.1 | 0 | 8.9977554 | 0.251 | 0.001 | 0 | Endo4 |
| PKHD1L1 | 0 | 8.9935872 | 0.869 | 0.008 | 0 | Endo4 |
| RELN | 0 | 8.1952878 | 0.598 | 0.01 | 0 | Endo4 |
| PIK3C2G | 0 | 7.9700053 | 0.464 | 0.006 | 0 | Endo4 |
| DCDC2C | 0 | 7.886752 | 0.45 | 0.004 | 0 | Endo4 |
| STAB2 | 0 | 7.7327673 | 0.282 | 0.005 | 0 | Endo4 |
| PARD6G | 0 | 7.5561587 | 0.653 | 0.011 | 0 | Endo4 |
| PROX1-AS1 | 0 | 7.4832267 | 0.43 | 0.005 | 0 | Endo4 |
| AC093912.1 | 0 | 7.237228 | 0.402 | 0.008 | 0 | Endo4 |
| PROX1 | 0 | 7.1133995 | 0.729 | 0.022 | 0 | Endo4 |
| MMRN11 | 0 | 7.0838882 | 0.784 | 0.02 | 0 | Endo4 |
| DSP | 0 | 7.0771456 | 0.268 | 0.003 | 0 | Endo4 |
| LINC01748 | 0 | 7.0238038 | 0.333 | 0.007 | 0 | Endo4 |
| POF1B | 0 | 6.9571269 | 0.306 | 0.004 | 0 | Endo4 |
| ART4 | 0 | 6.8555411 | 0.378 | 0.008 | 0 | Endo4 |
| NRG3 | 0 | 6.8344625 | 0.825 | 0.016 | 0 | Endo4 |
| FLT4 | 0 | 6.6799478 | 0.46 | 0.008 | 0 | Endo4 |
| AC099792.1 | 0 | 6.4431093 | 0.364 | 0.013 | 0 | Endo4 |
| AL357507.1 | 0 | 6.2699515 | 0.481 | 0.031 | 0 | Endo4 |
| ADIPOQ-AS1 | 0 | 9.3320902 | 0.299 | 0 | 0 | Adipocytes |
| TBL1Y | 0 | 9.282209 | 0.564 | 0.003 | 0 | Adipocytes |
| CIDEC | 0 | 9.004709 | 0.746 | 0.002 | 0 | Adipocytes |
| AL845331.2 | 0 | 8.954722 | 0.614 | 0.002 | 0 | Adipocytes |
| CIDEA | 0 | 8.9149237 | 0.77 | 0.003 | 0 | Adipocytes |
| GLYAT | 0 | 8.8566201 | 0.783 | 0.003 | 0 | Adipocytes |
| LGALS12 | 0 | 8.5793381 | 0.537 | 0.002 | 0 | Adipocytes |
| GYG2 | 0 | 8.5052858 | 0.889 | 0.006 | 0 | Adipocytes |
| ADIPOQ | 0 | 8.4078948 | 0.87 | 0.004 | 0 | Adipocytes |

| | | | | | | |
|---|---|---|---|---|---|---|
| LINC02237 | 0 | 8.1918187 | 0.903 | 0.009 | 0 | Adipocytes |
| PLIN1 | 0 | 8.1158992 | 0.976 | 0.015 | 0 | Adipocytes |
| TRARG1 | 0 | 8.1138099 | 0.462 | 0.001 | 0 | Adipocytes |
| LINCADL | 0 | 7.7872992 | 0.433 | 0.002 | 0 | Adipocytes |
| DGAT2 | 0 | 7.7624792 | 0.462 | 0.003 | 0 | Adipocytes |
| GPD1 | 0 | 7.7381021 | 0.79 | 0.007 | 0 | Adipocytes |
| SLC7A10 | 0 | 7.7275139 | 0.348 | 0.001 | 0 | Adipocytes |
| AQP7 | 0 | 7.5595945 | 0.955 | 0.019 | 0 | Adipocytes |
| KLB | 0 | 7.4169032 | 0.625 | 0.006 | 0 | Adipocytes |
| AC025470.2 | 0 | 7.4063953 | 0.286 | 0.002 | 0 | Adipocytes |
| ALDH1L1 | 0 | 7.2988269 | 0.567 | 0.003 | 0 | Adipocytes |

**Table S4: Mouse fibroblast subcluster transcriptomic profiles.** Top 20 differentially-expressed genes (DEGs) in fibroblast subclusters found in mouse synovia subjected to single nuclei RNA sequencing (healthy control; n=4, 2 weeks; n=3, 10 weeks; n=4). DEGs are organized by decreasing average $\log_2$-fold change (FC) and filtered by adjusted p-value < 0.05, $\log_2FC \geq 0.5$ and min.pct=0.25. Full List available upon request. Raw data deposited in GEO Accession Number: GSE281724.

| gene | p_val | avg_log2FC | pct.1 | pct.2 | p_val_adj | cluster |
|---|---|---|---|---|---|---|
| Edil3 | 1.36E-302 | 2.958026749 | 0.727 | 0.169 | 2.91E-298 | 0 |
| Myo16 | 1.69E-126 | 2.839395865 | 0.335 | 0.066 | 3.63E-122 | 0 |
| Smoc1 | 2.33E-106 | 2.45410652 | 0.371 | 0.098 | 5E-102 | 0 |
| Acan | 1.685E-97 | 2.424836799 | 0.251 | 0.044 | 3.614E-93 | 0 |
| Tnc | 5.49E-140 | 2.330895103 | 0.547 | 0.176 | 1.18E-135 | 0 |
| Cemip | 3.304E-73 | 2.30948628 | 0.294 | 0.085 | 7.087E-69 | 0 |
| Slc24a3 | 1.147E-89 | 1.986965109 | 0.388 | 0.123 | 2.461E-85 | 0 |
| Postn | 2.86E-123 | 1.976906734 | 0.572 | 0.209 | 6.13E-119 | 0 |
| Dkk3 | 6.06E-123 | 1.948905907 | 0.457 | 0.131 | 1.3E-118 | 0 |
| Col24a1 | 8.855E-57 | 1.941761096 | 0.25 | 0.076 | 1.9E-52 | 0 |
| Runx2 | 8.57E-106 | 1.868638109 | 0.511 | 0.19 | 1.84E-101 | 0 |
| Cthrc1 | 5.55E-112 | 1.863328402 | 0.445 | 0.133 | 1.19E-107 | 0 |
| Cdh11 | 1.03E-132 | 1.838156266 | 0.536 | 0.172 | 2.21E-128 | 0 |
| Palld | 4.32E-161 | 1.827437211 | 0.718 | 0.293 | 9.28E-157 | 0 |
| Glis3 | 2.89E-125 | 1.827162864 | 0.714 | 0.343 | 6.2E-121 | 0 |
| Adam12 | 6.12E-100 | 1.794946287 | 0.443 | 0.146 | 1.312E-95 | 0 |
| Gxylt2 | 1.853E-96 | 1.792367427 | 0.463 | 0.164 | 3.974E-92 | 0 |
| Enpp1 | 1.38E-104 | 1.788049423 | 0.53 | 0.203 | 2.96E-100 | 0 |
| Lmo7 | 1.431E-88 | 1.769620331 | 0.309 | 0.076 | 3.069E-84 | 0 |
| Chn2 | 6.799E-69 | 1.76790996 | 0.317 | 0.102 | 1.459E-64 | 0 |
| Rab37 | 9.48E-214 | 5.601540695 | 0.257 | 0.006 | 2.03E-209 | 1 |
| Grip1 | 0 | 5.440330836 | 0.521 | 0.026 | 0 | 1 |
| Clic5 | 0 | 5.348128882 | 0.527 | 0.046 | 0 | 1 |
| Col22a1 | 0 | 5.332464672 | 0.832 | 0.077 | 0 | 1 |
| Grid2 | 0 | 5.245535365 | 0.668 | 0.06 | 0 | 1 |
| Gm48727 | 5.69E-252 | 5.076526288 | 0.361 | 0.023 | 1.22E-247 | 1 |
| Hbegf | 0 | 4.9128264 | 0.521 | 0.053 | 0 | 1 |
| Tspan15 | 1.15E-296 | 4.790740172 | 0.382 | 0.016 | 2.47E-292 | 1 |
| Rgcc | 0 | 4.531379 | 0.72 | 0.085 | 0 | 1 |
| Tmem196 | 0 | 4.450088674 | 0.589 | 0.064 | 0 | 1 |
| Fut9 | 3.6E-199 | 4.374547712 | 0.335 | 0.031 | 7.72E-195 | 1 |
| Htra4 | 2E-249 | 4.358043624 | 0.414 | 0.04 | 4.3E-245 | 1 |
| F13a1 | 0 | 4.289726247 | 0.772 | 0.123 | 0 | 1 |
| Mdga2 | 1.22E-205 | 4.087078252 | 0.377 | 0.044 | 2.62E-201 | 1 |
| Megf10 | 0 | 4.022857401 | 0.534 | 0.056 | 0 | 1 |
| Prg4 | 0 | 3.994884665 | 0.953 | 0.489 | 0 | 1 |
| Dlx3 | 5.25E-199 | 3.928246589 | 0.292 | 0.018 | 1.13E-194 | 1 |
| Robo2 | 6.56E-190 | 3.604843209 | 0.374 | 0.048 | 1.41E-185 | 1 |
| Efnb2 | 1.76E-184 | 3.529877411 | 0.369 | 0.048 | 3.78E-180 | 1 |
| Cyp1b1 | 6.84E-167 | 3.340554584 | 0.323 | 0.039 | 1.47E-162 | 1 |
| Dkk2 | 1.386E-85 | 3.222937788 | 0.318 | 0.081 | 2.973E-81 | 2 |
| Trpm3 | 1.206E-87 | 2.99077034 | 0.256 | 0.049 | 2.586E-83 | 2 |
| C4b | 1.482E-84 | 2.586297543 | 0.323 | 0.083 | 3.178E-80 | 2 |
| Ism1 | 3.113E-89 | 2.288432073 | 0.418 | 0.138 | 6.678E-85 | 2 |
| Slc1a3 | 1.797E-65 | 2.284685462 | 0.271 | 0.073 | 3.856E-61 | 2 |
| Il1r1 | 1.462E-90 | 2.213657526 | 0.562 | 0.253 | 3.137E-86 | 2 |
| Col6a6 | 2.286E-56 | 2.160542038 | 0.278 | 0.085 | 4.903E-52 | 2 |
| Bmper | 2.722E-65 | 2.101915374 | 0.483 | 0.228 | 5.838E-61 | 2 |
| Fgf7 | 2.849E-66 | 2.010838929 | 0.318 | 0.097 | 6.112E-62 | 2 |
| Csmd1 | 1.746E-40 | 1.997683946 | 0.26 | 0.097 | 3.746E-36 | 2 |
| Gm765 | 4.741E-40 | 1.939200429 | 0.33 | 0.149 | 1.017E-35 | 2 |
| Sntg2 | 4.395E-43 | 1.90739111 | 0.253 | 0.087 | 9.428E-39 | 2 |
| Prr161 | 5.592E-51 | 1.882867358 | 0.327 | 0.123 | 1.2E-46 | 2 |
| Lsamp | 4.61E-73 | 1.782941691 | 0.464 | 0.189 | 9.889E-69 | 2 |
| Gas6 | 4.832E-48 | 1.699003681 | 0.425 | 0.206 | 1.037E-43 | 2 |

| | | | | | | |
|---|---|---|---|---|---|---|
| Map1b | 8.056E-48 | 1.636474435 | 0.5 | 0.278 | 1.728E-43 | 2 |
| Rassf2 | 5.839E-30 | 1.583717665 | 0.257 | 0.11 | 1.253E-25 | 2 |
| Foxp2 | 1.571E-24 | 1.571080135 | 0.269 | 0.135 | 3.37E-20 | 2 |
| Dpyd | 4.185E-46 | 1.553302571 | 0.306 | 0.115 | 8.977E-42 | 2 |
| Angpt1 | 1.096E-52 | 1.446855249 | 0.427 | 0.191 | 2.351E-48 | 2 |
| Hmcn2 | 0 | 3.716993949 | 0.727 | 0.106 | 0 | 3 |
| Bmp5 | 1.03E-147 | 3.426347933 | 0.325 | 0.044 | 2.21E-143 | 3 |
| Smoc2 | 1.02E-167 | 3.33275919 | 0.511 | 0.127 | 2.2E-163 | 3 |
| Abca8a | 1.06E-275 | 2.964321677 | 0.756 | 0.19 | 2.28E-271 | 3 |
| Prex2 | 1.89E-115 | 2.842334883 | 0.343 | 0.069 | 4.04E-111 | 3 |
| Abca8b | 1.31E-165 | 2.689296438 | 0.576 | 0.162 | 2.81E-161 | 3 |
| Abca9 | 1.66E-109 | 2.675986523 | 0.447 | 0.137 | 3.56E-105 | 3 |
| Lvrn | 6.68E-111 | 2.659046243 | 0.351 | 0.075 | 1.43E-106 | 3 |
| Myoc | 4.31E-150 | 2.636149818 | 0.484 | 0.113 | 9.24E-146 | 3 |
| Etl4 | 1.397E-87 | 2.516008947 | 0.413 | 0.14 | 2.997E-83 | 3 |
| Sgip1 | 4.04E-124 | 2.403688912 | 0.443 | 0.112 | 8.67E-120 | 3 |
| Abca6 | 2.565E-66 | 2.312449581 | 0.3 | 0.088 | 5.503E-62 | 3 |
| Pcdh7 | 6.55E-208 | 2.255358181 | 0.833 | 0.377 | 1.41E-203 | 3 |
| Podn | 3.189E-46 | 2.254935327 | 0.255 | 0.087 | 6.841E-42 | 3 |
| B3galt1 | 7.641E-51 | 2.20535636 | 0.289 | 0.102 | 1.639E-46 | 3 |
| Wdr66 | 1.9E-52 | 2.151305459 | 0.313 | 0.116 | 4.076E-48 | 3 |
| Ltbp4 | 1.558E-58 | 2.127796465 | 0.363 | 0.144 | 3.342E-54 | 3 |
| Vit | 3.682E-68 | 2.085014845 | 0.372 | 0.132 | 7.899E-64 | 3 |
| Galnt17 | 2.06E-99 | 2.060115374 | 0.628 | 0.316 | 4.428E-95 | 3 |
| Gsn | 9.99E-126 | 2.002899913 | 0.845 | 0.574 | 2.14E-121 | 3 |
| Pi16 | 0 | 4.730129885 | 0.604 | 0.058 | 0 | 4 |
| Efhd1 | 2.02E-255 | 4.361761515 | 0.392 | 0.027 | 4.34E-251 | 4 |
| Opcml | 0 | 4.347825285 | 0.636 | 0.083 | 0 | 4 |
| Cmah | 7.81E-226 | 4.281634257 | 0.398 | 0.039 | 1.68E-221 | 4 |
| Rorb | 9.75E-238 | 3.886666653 | 0.453 | 0.053 | 2.09E-233 | 4 |
| Dpp4 | 0 | 3.732486903 | 0.641 | 0.083 | 0 | 4 |
| Efemp1 | 4.78E-203 | 3.593489218 | 0.431 | 0.06 | 1.03E-198 | 4 |
| Anxa3 | 3.58E-188 | 3.193095517 | 0.579 | 0.151 | 7.68E-184 | 4 |
| Tek | 6.76E-203 | 3.072270922 | 0.467 | 0.072 | 1.45E-198 | 4 |
| Slc43a3 | 4.734E-94 | 2.913122583 | 0.301 | 0.064 | 1.016E-89 | 4 |
| Efna5 | 4.85E-102 | 2.910513455 | 0.405 | 0.117 | 1.04E-97 | 4 |
| Tmeff21 | 4.15E-236 | 2.909432273 | 0.637 | 0.135 | 8.91E-232 | 4 |
| Negr1 | 2.093E-89 | 2.732616419 | 0.306 | 0.071 | 4.49E-85 | 4 |
| Scara51 | 2.57E-250 | 2.668025483 | 0.735 | 0.181 | 5.52E-246 | 4 |
| Dact1 | 8.882E-78 | 2.651752371 | 0.253 | 0.053 | 1.905E-73 | 4 |
| Cadm3 | 1.58E-118 | 2.64857724 | 0.367 | 0.076 | 3.38E-114 | 4 |
| Galnt16 | 1.3E-143 | 2.630660269 | 0.489 | 0.123 | 2.8E-139 | 4 |
| Lurap1l | 4.168E-78 | 2.587799015 | 0.331 | 0.095 | 8.94E-74 | 4 |
| Gas7 | 3.46E-162 | 2.570445095 | 0.677 | 0.24 | 7.42E-158 | 4 |
| Fbn1 | 7.49E-247 | 2.537156199 | 0.929 | 0.566 | 1.61E-242 | 4 |
| Angptl7 | 2.22E-130 | 4.499299757 | 0.26 | 0.021 | 4.75E-126 | 5 |
| P3h2 | 3.5E-197 | 4.410064684 | 0.451 | 0.049 | 7.51E-193 | 5 |
| Kctd8 | 7.17E-136 | 4.377897457 | 0.274 | 0.023 | 1.54E-131 | 5 |
| Cilp2 | 6.65E-211 | 4.291643316 | 0.435 | 0.039 | 1.43E-206 | 5 |
| Mkx | 2.63E-215 | 4.220584504 | 0.54 | 0.072 | 5.65E-211 | 5 |
| Nalcn | 5.18E-216 | 4.133594589 | 0.419 | 0.034 | 1.11E-211 | 5 |
| Map2 | 8.1E-165 | 4.081521513 | 0.372 | 0.038 | 1.74E-160 | 5 |
| Mylk | 6.9E-207 | 4.011924371 | 0.578 | 0.089 | 1.48E-202 | 5 |
| Thbs4 | 2.96E-162 | 3.193136755 | 0.688 | 0.187 | 6.35E-158 | 5 |
| Eya2 | 5.535E-75 | 2.918550491 | 0.307 | 0.062 | 1.187E-70 | 5 |
| Mpp7 | 1.314E-88 | 2.800863636 | 0.581 | 0.222 | 2.818E-84 | 5 |
| Fmod | 7.628E-90 | 2.796632479 | 0.368 | 0.075 | 1.636E-85 | 5 |
| Adamts3 | 8.596E-45 | 2.745323183 | 0.289 | 0.086 | 1.844E-40 | 5 |
| Ppfibp2 | 3.959E-89 | 2.676342203 | 0.428 | 0.106 | 8.493E-85 | 5 |
| Ptgis | 1.684E-85 | 2.643052243 | 0.426 | 0.108 | 3.612E-81 | 5 |
| Kcnma11 | 2.23E-180 | 2.597531606 | 0.915 | 0.358 | 4.78E-176 | 5 |
| Ccdc31 | 1.445E-88 | 2.546628173 | 0.587 | 0.213 | 3.099E-84 | 5 |
| Col11a11 | 4.37E-110 | 2.424465985 | 0.563 | 0.157 | 9.37E-106 | 5 |
| Itga2 | 1.275E-43 | 2.372779183 | 0.298 | 0.092 | 2.736E-39 | 5 |

| | | | | | | |
|---|---|---|---|---|---|---|
| Crispld2 | 1.003E-32 | 2.209780072 | 0.287 | 0.103 | 2.152E-28 | 5 |
| Mylpf | 3.913E-82 | 5.209229621 | 0.305 | 0.052 | 8.394E-78 | 6 |
| Tnnc2 | 2.25E-107 | 4.868750976 | 0.384 | 0.065 | 4.82E-103 | 6 |
| Tnni2 | 2.703E-88 | 4.798000851 | 0.329 | 0.057 | 5.798E-84 | 6 |
| Acta1 | 1.701E-77 | 4.628470669 | 0.424 | 0.114 | 3.648E-73 | 6 |
| Pvalb | 5.685E-77 | 4.34588604 | 0.318 | 0.061 | 1.219E-72 | 6 |
| Gapdh | 4.78E-111 | 4.29641245 | 0.316 | 0.039 | 1.03E-106 | 6 |
| Ckm | 1.029E-69 | 4.155523929 | 0.397 | 0.106 | 2.208E-65 | 6 |
| Myl1 | 1.516E-67 | 3.902125195 | 0.358 | 0.088 | 3.252E-63 | 6 |
| Atp5e | 1.004E-79 | 3.895659904 | 0.297 | 0.05 | 2.154E-75 | 6 |
| Aldoa | 1.11E-82 | 3.804101389 | 0.437 | 0.112 | 2.381E-78 | 6 |
| Tnnt3 | 3.485E-50 | 3.544464279 | 0.408 | 0.144 | 7.475E-46 | 6 |
| Myh4 | 2.853E-43 | 3.379520619 | 0.495 | 0.232 | 6.121E-39 | 6 |
| Cox6c | 1.09E-53 | 3.220317811 | 0.324 | 0.086 | 2.339E-49 | 6 |
| Apoe | 2.578E-70 | 3.214420652 | 0.397 | 0.103 | 5.53E-66 | 6 |
| Ubb | 4.328E-53 | 3.19537247 | 0.279 | 0.065 | 9.285E-49 | 6 |
| Lyz2 | 1.325E-73 | 3.179349643 | 0.411 | 0.105 | 2.842E-69 | 6 |
| Ctss | 6.979E-75 | 3.176583595 | 0.255 | 0.038 | 1.497E-70 | 6 |
| Cox7c | 5.957E-68 | 3.08053027 | 0.371 | 0.092 | 1.278E-63 | 6 |
| Tpm1 | 3.051E-51 | 3.047438719 | 0.561 | 0.265 | 6.545E-47 | 6 |
| Atp2a1 | 2.091E-54 | 3.022500998 | 0.461 | 0.171 | 4.485E-50 | 6 |
| Sema7a | 7.539E-48 | 3.681796152 | 0.302 | 0.03 | 1.617E-43 | 7 |
| Serpine1 | 2.608E-53 | 3.64672101 | 0.49 | 0.073 | 5.596E-49 | 7 |
| Notch3 | 4.278E-41 | 3.542705604 | 0.281 | 0.03 | 9.178E-37 | 7 |
| 8030451A03Rik | 2.275E-29 | 3.411442134 | 0.281 | 0.042 | 4.879E-25 | 7 |
| Egr3 | 9.93E-54 | 3.391170006 | 0.281 | 0.022 | 2.13E-49 | 7 |
| Gpr176 | 3.455E-60 | 3.224968401 | 0.542 | 0.078 | 7.413E-56 | 7 |
| Rgs6 | 3.424E-27 | 3.166258021 | 0.354 | 0.069 | 7.344E-23 | 7 |
| Pakap | 1.901E-31 | 3.105961456 | 0.385 | 0.071 | 4.077E-27 | 7 |
| Thbs1 | 3.129E-35 | 3.060856975 | 0.438 | 0.082 | 6.712E-31 | 7 |
| Mical21 | 5.785E-42 | 2.949780625 | 0.646 | 0.162 | 1.241E-37 | 7 |
| Cd80 | 1.82E-46 | 2.939960701 | 0.521 | 0.09 | 3.905E-42 | 7 |
| Pde3a2 | 9.344E-35 | 2.933561102 | 0.729 | 0.253 | 2.005E-30 | 7 |
| Tnc1 | 4.481E-34 | 2.716046952 | 0.729 | 0.239 | 9.612E-30 | 7 |
| Gm13986 | 4.171E-35 | 2.688204009 | 0.25 | 0.027 | 8.948E-31 | 7 |
| Col18a1 | 5.989E-29 | 2.683570253 | 0.427 | 0.09 | 1.285E-24 | 7 |
| Grem2 | 8.251E-37 | 2.68286437 | 0.333 | 0.045 | 1.77E-32 | 7 |
| Col8a11 | 1.401E-38 | 2.678100345 | 0.75 | 0.233 | 3.006E-34 | 7 |
| Prss23 | 3.262E-26 | 2.633428548 | 0.406 | 0.092 | 6.997E-22 | 7 |
| Carmn | 5.964E-11 | 2.53597048 | 0.25 | 0.075 | 1.279E-06 | 7 |
| Vcl1 | 2.019E-16 | 2.507559267 | 0.625 | 0.314 | 4.332E-12 | 7 |

**Table S5: Enriched transcription factors that putatively regulate differentially-expressed genes of fibroblast subcluster 1 (A) and fibroblast subcluster 0 (B).** Catrin analysis performed on ECM related genes from single nuclei RNA sequencing, identifying transcription factors enriched in fibroblast subcluster 1 (early-stage predominant) (A) and fibroblast subcluster 0 (advanced-stage predominant) (B). Transcription factors are filtered based on adjusted p<0.01.

A

| genes | targets_early_1 | targetsCatrin | p_value | p_adjust |
|---|---|---|---|---|
| CBX8 | 50 | 3427 | 6.602E-05 | 0.0005435 |
| CTNNB1 | 78 | 6445 | 0.000232 | 0.0016742 |
| DROSHA | 19 | 456 | 1.373E-08 | 3.255E-07 |
| EMX1 | 110 | 9069 | 1.932E-06 | 2.406E-05 |
| ETS1 | 166 | 16915 | 0.0002915 | 0.001975 |
| FOXH1 | 78 | 6704 | 0.0008358 | 0.0050149 |
| GATA1 | 169 | 17675 | 0.0013853 | 0.0078841 |
| GRHL3 | 94 | 8239 | 0.0003116 | 0.0020828 |
| GSPT2 | 39 | 2776 | 0.0010385 | 0.0060138 |
| HSF1 | 147 | 14662 | 0.0006701 | 0.004155 |
| IKZF1 | 144 | 13028 | 1.455E-06 | 1.932E-05 |
| KDM6B | 116 | 10013 | 8.794E-06 | 9.418E-05 |
| KLF11 | 128 | 11027 | 1.013E-06 | 1.382E-05 |
| KLF4 | 139 | 12746 | 9.082E-06 | 9.501E-05 |
| KLF9 | 131 | 12129 | 5.387E-05 | 0.0004586 |
| KMT2A | 116 | 10842 | 0.0004897 | 0.003147 |
| LARP7 | 59 | 3993 | 7.512E-06 | 8.222E-05 |
| LIN28A | 67 | 4583 | 1.921E-06 | 2.406E-05 |
| LIN28B | 85 | 6682 | 1.353E-05 | 0.0001321 |
| MED26 | 40 | 2357 | 1.576E-05 | 0.0001495 |
| MYOD1 | 114 | 9506 | 1.788E-06 | 2.312E-05 |
| NEUROG2 | 52 | 3739 | 0.0001571 | 0.0011764 |
| NFYA | 155 | 15734 | 0.0008568 | 0.0051099 |
| NKX2-2 | 101 | 8980 | 0.0002587 | 0.0018147 |
| OTX2 | 130 | 12343 | 0.0002502 | 0.0017803 |
| PAX8 | 29 | 1917 | 0.0016734 | 0.0092082 |
| PBX3 | 152 | 15293 | 0.0006725 | 0.004155 |
| PIAS1 | 96 | 8679 | 0.0008275 | 0.004995 |
| POLR2B | 78 | 6659 | 0.0006758 | 0.004155 |
| POU2F1 | 40 | 2565 | 0.0001047 | 0.0008024 |
| SP2 | 122 | 11832 | 0.0015418 | 0.0086328 |
| SP4 | 136 | 13095 | 0.0002798 | 0.0019223 |
| SUPT5H | 89 | 7877 | 0.0007454 | 0.0045546 |
| TTF2 | 77 | 6732 | 0.0015507 | 0.0086328 |
| WDR5 | 126 | 12314 | 0.0015779 | 0.0087311 |
| XRN2 | 109 | 9880 | 0.0002501 | 0.0017803 |
| ZC3H7B | 63 | 4602 | 3.796E-05 | 0.0003316 |
| ZNF224 | 14 | 594 | 0.0005759 | 0.0036073 |
| ZSCAN22 | 87 | 7555 | 0.0004663 | 0.0030352 |

B

| gene | targets_late_0 | targetsCatrin | p_value | p_adjust |
|---|---|---|---|---|
| AHR | 106 | 9329 | 0.0002973 | 0.0021637 |
| ARNT | 150 | 13633 | 5.66E-06 | 5.76E-05 |
| BHLHE40 | 171 | 16803 | 7.65E-05 | 0.0006118 |
| CDK8 | 134 | 12906 | 0.0014394 | 0.0090207 |
| CHD8 | 133 | 11610 | 5.93E-06 | 5.97E-05 |
| CKAP4 | 49 | 3654 | 0.0010453 | 0.0068381 |

| | | | | |
|---|---|---|---|---|
| CSH2 | 34 | 1947 | 7.04E-05 | 0.000577 |
| CTCFL | 136 | 12673 | 0.000203 | 0.001498 |
| EED | 88 | 7655 | 0.0010871 | 0.0070326 |
| EGLN2 | 89 | 7137 | 4.77E-05 | 0.0004062 |
| EHMT2 | 81 | 6913 | 0.0011096 | 0.0071338 |
| EIF3D | 22 | 1191 | 0.000713 | 0.0047592 |
| ELF1 | 177 | 17490 | 4.36E-05 | 0.0003775 |
| FUS | 60 | 3454 | 5.80E-08 | 9.96E-07 |
| GABPA | 181 | 17807 | 1.27E-05 | 0.0001185 |
| GRHL2 | 91 | 7820 | 0.0005327 | 0.0036488 |
| GTF2B | 135 | 12746 | 0.0004668 | 0.0032183 |
| IRF4 | 158 | 15795 | 0.0013731 | 0.0087212 |
| MECP2 | 103 | 9174 | 0.0006562 | 0.0044659 |
| NRF1 | 150 | 14634 | 0.0006737 | 0.0045548 |
| PGR | 166 | 16138 | 7.10E-05 | 0.000577 |
| POU3F2 | 56 | 4409 | 0.0015389 | 0.0093768 |
| PUM2 | 80 | 5120 | 1.52E-08 | 3.13E-07 |
| RAG2 | 41 | 2562 | 8.22E-05 | 0.0006481 |
| RCOR1 | 182 | 17751 | 4.16E-06 | 4.28E-05 |
| SMAD1 | 118 | 10774 | 0.0004614 | 0.0032023 |
| SMC1A | 136 | 12957 | 0.0006876 | 0.0046195 |
| UBN1 | 49 | 3500 | 0.0003955 | 0.0027634 |
| USF2 | 173 | 16140 | 4.53E-07 | 6.01E-06 |
| XBP1 | 131 | 12579 | 0.0015878 | 0.0096188 |
| ZFP64 | 93 | 7554 | 4.67E-05 | 0.0004011 |

**Table S6: Enriched transcription factors in DEG lists of advanced-stage fibroblast subcluster 0 (ITGB8+) (A) and early-stage fibroblast subcluster 1 (DPP4+) (B) listed.** Table is ordered of most to least number of putatively-targeted ECM genes by individual transcription factors.

A) Subcluster 0 (ITGB8+)

| # | Transcription Factor | Number of Target Genes | Target Genes |
|---|---|---|---|
| 1 | RCOR1 | 23 | ADAM10, ADAMTS3, CD63, COL22A1, COL5A2, CSGALNACT1, CYP1B1, FN1, HTRA1, ITGAV, ITGB8, ITGBL1, LTBP3, NTN4, PLOD2, PRKCA, SAT1, SH3PXD2A, TIMP1, TNC, VAV3, VCAM1, VEGFC |
| 2 | USF2 | 23 | ADAM10, ADAMTS3, CD63, COL22A1, COL5A2, CSGALNACT1, CYP1B1, FN1, HTRA1, ITGAV, ITGB8, ITGBL1, LTBP3, NTN4, PLOD2, PRKCA, SAT1, SH3PXD2A, TIMP1, TNC, VAV3, VCAM1, VEGFC |
| 3 | GABPA | 21 | ADAM10, ADAMTS3, CD63, COL22A1, CSGALNACT1, CYP1B1, FN1, HTRA1, ITGAV, ITGB8, LTBP3, NTN4, PLOD2, PRKCA, SAT1, SH3PXD2A, TIMP1, TNC, VAV3, VCAM1, VEGFC |
| 4 | BHLHE40 | 20 | ADAM10, ADAMTS3, CD63, CSGALNACT1, CYP1B1, FN1, HTRA1, ITGAV, ITGB8, ITGBL1, NTN4, PLOD2, PRKCA, SAT1, SH3PXD2A, TIMP1, TNC, VAV3, VCAM1, VEGFC |
| 5 | ELF1 | 19 | ADAM10, ADAMTS3, CD63, COL22A1, CSGALNACT1, CYP1B1, FN1, HTRA1, ITGAV, ITGB8, LTBP3, PLOD2, PRKCA, SAT1, SH3PXD2A, TIMP1, TNC, VAV3, VEGFC |
| 6 | PGR | 18 | ADAM10, ADAMTS3, CD63, COL22A1, CYP1B1, FN1, ITGAV, ITGB8, LTBP3, NTN4, PLOD2, PRKCA, SAT1, SH3PXD2A, TIMP1, TNC, VAV3, VEGFC |
| 7 | ARNT | 17 | ADAMTS3, COL22A1, COL5A2, CSGALNACT1, CYP1B1, FN1, HTRA1, ITGAV, ITGB8, LTBP3, NTN4, PLOD2, PRKCA, SAT1, TNC, VAV3, VEGFC |
| 8 | SMAD1 | 17 | ADAM10, ADAMTS3, CD63, COL22A1, CSGALNACT1, FN1, HTRA1, ITGAV, ITGB8, LTBP3, PLOD2, SAT1, TIMP1, TNC, VAV3, VCAM1, VEGFC |

| | | | |
|---|---|---|---|
| 9 | SMC1A | 17 | ADAM10, ADAMTS3, CD63, COL22A1, CYP1B1, HTRA1, ITGAV, ITGB8, ITGBL1, LTBP3, NTN4, PRKCA, SAT1, TIMP1, TNC, VAV3, VEGFC |
| 10 | AHR | 16 | ADAMTS3, COL22A1, CYP1B1, FN1, ITGAV, ITGB8, LTBP3, NTN4, PLOD2, PRKCA, SAT1, SH3PXD2A, TIMP1, TNC, VAV3, VEGFC |
| 11 | CHD8 | 16 | ADAM10, ADAMTS3, CD63, CSGALNACT1, FN1, HTRA1, ITGAV, ITGB8, NTN4, PRKCA, SAT1, SH3PXD2A, TIMP1, TNC, VAV3, VCAM1 |
| 12 | CDK8 | 15 | ADAM10, ADAMTS3, CSGALNACT1, FN1, HTRA1, ITGAV, ITGB8, LTBP3, PRKCA, SAT1, SH3PXD2A, TIMP1, TNC, VAV3, VCAM1 |
| 13 | NRF1 | 15 | ADAM10, ADAMTS3, CD63, COL22A1, CYP1B1, FN1, HTRA1, ITGAV, ITGB8, PRKCA, SAT1, SH3PXD2A, TNC, VAV3, VEGFC |
| 14 | IRF4 | 14 | ADAM10, ADAMTS3, CD63, COL22A1, CYP1B1, FN1, ITGAV, ITGB8, PRKCA, SAT1, TIMP1, VAV3, VCAM1, VEGFC |
| 15 | CTCFL | 13 | ADAM10, ADAMTS3, COL22A1, CSGALNACT1, CYP1B1, ITGAV, ITGB8, LTBP3, PRKCA, SAT1, SH3PXD2A, VAV3, VEGFC |
| 16 | XBP1 | 12 | ADAM10, ADAMTS3, CD63, CSGALNACT1, CYP1B1, ITGAV, ITGB8, LTBP3, PRKCA, SAT1, VAV3, VCAM1 |
| 17 | EGLN2 | 11 | CD63, COL22A1, FN1, HTRA1, ITGB8, NTN4, PRKCA, SAT1, TIMP1, VAV3, VEGFC |
| 18 | MECP2 | 11 | ADAM10, ADAMTS3, CD63, COL5A2, CSGALNACT1, FN1, ITGB8, PRKCA, TNC, VCAM1, VEGFC |
| 19 | EED | 10 | ADAM10, ADAMTS3, CD63, FN1, HTRA1, ITGAV, ITGB8, NTN4, TNC, VAV3 |
| 20 | GRHL2 | 10 | ADAM10, ADAMTS3, CD63, CSGALNACT1, CYP1B1, ITGB8, LTBP3, PLOD2, SAT1, VCAM1 |
| 21 | GTF2B | 10 | ADAM10, ADAMTS3, CD63, FN1, HTRA1, ITGAV, ITGB8, PRKCA, SAT1, VCAM1 |
| 22 | ZFP64 | 10 | CD63, CSGALNACT1, FN1, ITGAV, ITGB8, LTBP3, NTN4, PLOD2, SH3PXD2A, VAV3 |

| | | | |
|---|---|---|---|
| 23 | EHMT2 | 8 | ADAM10, ADAMTS3, FN1, ITGAV, ITGB8, NTN4, PRKCA, VEGFC |
| 24 | PUM2 | 7 | ADAM10, ADAMTS3, ITGAV, LTBP3, PRKCA, SAT1, VAV3 |
| 25 | UBN1 | 7 | ADAMTS3, FN1, ITGB8, LTBP3, PLOD2, SAT1, TNC |
| 26 | CKAP4 | 6 | HTRA1, ITGAV, ITGBL1, NTN4, SH3PXD2A, VAV3 |
| 27 | FUS | 6 | ADAM10, ITGAV, ITGB8, PLOD2, SAT1, VAV3 |
| 28 | EIF3D | 5 | ADAM10, FN1, ITGAV, LTBP3, SAT1 |
| 29 | POU3F2 | 5 | ADAM10, COL5A2, FN1, ITGB8, VAV3 |
| 30 | RAG2 | 5 | ADAMTS3, FN1, ITGB8, PRKCA, VAV3 |
| 31 | CSH2 | 3 | PLOD2, SH3PXD2A, TNC |

B) Subcluster 1 (DPP4+)

| # | Transcription Factor | Number of Target Genes | Target Genes |
|---|---|---|---|
| 1 | IKZF1 | 20 | ADAMTS5, APBB2, DDR2, DOCK4, DPP4, EMILIN2, FBLN2, FBN1, FLRT2, ITGA11, LTBP1, LTBP4, NID1, PXN, RAP1A, RAP1B, SDC2, ST7, TNXB, VIT |
| 2 | GATA1 | 19 | ADAMTS5, APBB2, DDR2, DOCK4, DPP4, EMILIN2, FBLN2, FBN1, FLRT2, LTBP1, MFAP5, NID1, PCOLCE2, PXN, RAP1A, RAP1B, RECK, ST7, TNXB |
| 3 | NFYA | 19 | ADAMTS5, APBB2, DDR2, DOCK4, EMILIN2, FBLN2, FBN1, FLRT2, ITGA11, LTBP1, LTBP4, MFAP5, NID1, PCOLCE2, PXN, RAP1A, RAP1B, ST7, VIT |
| 4 | ETS1 | 18 | ADAMTS5, APBB2, DDR2, DOCK4, DPP4, EMILIN2, FBLN2, FBN1, FLRT2, ITGA11, LTBP1, NID1, PCOLCE2, PXN, RAP1A, RAP1B, SDC2, ST7 |
| 5 | PBX3 | 17 | ADAMTS5, APBB2, DCN, DDR2, DOCK4, DPP4, FBN1, FLRT2, ITGA11, LTBP1, NID1, PXN, RAP1A, RAP1B, SDC2, ST7, VIT |
| 6 | HSF1 | 14 | ADAMTS5, APBB2, DOCK4, EMILIN2, FBLN2, FBN1, FLRT2, ITGA11, LTBP1, NID1, PXN, RAP1A, RAP1B, ST7 |
| 7 | KLF11 | 13 | ADAMTS5, APBB2, DOCK4, EMILIN2, FBN1, FLRT2, ITGA11, LTBP1, MFAP5, NID1, PCOLCE2, PXN, RAP1B |

| | | | |
|---|---|---|---|
| 8 | KLF4 | 13 | DCN, DOCK4, DPP4, EMILIN2, FBN1, FLRT2, LTBP1, MFAP5, NID1, PXN, RAP1B, ST7, VIT |
| 9 | MYOD1 | 13 | ADAMTS5, APBB2, DDR2, DPP4, FBLN2, FBN1, FLRT2, ITGA11, MFAP5, PXN, RAP1A, RAP1B, VIT |
| 10 | OTX2 | 13 | APBB2, DOCK4, DPP4, FBN1, FLRT2, ITGA11, LTBP1, NID1, PCOLCE2, PXN, RAP1A, RAP1B, ST7 |
| 11 | WDR5 | 13 | ADAMTS5, APBB2, DOCK4, EMILIN2, FBLN2, FBN1, ITGA11, LTBP1, NID1, PCOLCE2, PXN, RAP1B, ST7 |
| 12 | KMT2A | 12 | ADAMTS5, APBB2, DOCK4, DPP4, EMILIN2, ITGA11, LTBP1, NID1, PCOLCE2, PXN, RAP1B, ST7 |
| 13 | KDM6B | 11 | ADAMTS5, DOCK4, DPP4, EMILIN2, FLRT2, LTBP1, NID1, PCOLCE2, PXN, RAP1A, RAP1B |
| 14 | SP2 | 11 | APBB2, DOCK4, EMILIN2, FBN1, NID1, PCOLCE2, PXN, RAP1A, RAP1B, RECK, SDC2 |
| 15 | EMX1 | 10 | ADAMTS5, DOCK4, EMILIN2, FBN1, LTBP1, NID1, PCOLCE2, RAP1B, ST7, VIT |
| 16 | KLF9 | 10 | ADAMTS5, APBB2, DOCK4, FBLN2, ITGA11, LTBP1, NID1, PCOLCE2, PXN, RAP1B |
| 17 | PIAS1 | 10 | ADAMTS5, DDR2, DOCK4, DPP4, FLRT2, ITGA11, LTBP1, PCOLCE2, PXN, ST7 |
| 18 | SP4 | 10 | APBB2, DOCK4, FBN1, LTBP1, NID1, PCOLCE2, PXN, RAP1B, RECK, SDC2 |
| 19 | GRHL3 | 9 | ADAMTS5, APBB2, DOCK4, FBLN2, FBN1, ITGA11, LTBP4, NID1, RAP1A |
| 20 | POLR2B | 9 | ADAMTS5, APBB2, DOCK4, FBN1, FLRT2, ITGA11, LTBP1, PXN, RAP1B |
| 21 | FOXH1 | 8 | APBB2, DPP4, EMILIN2, FBN1, ITGA11, LTBP4, RECK, ST7 |
| 22 | NKX2-2 | 8 | DOCK4, DPP4, FBN1, FLRT2, LTBP4, PXN, SDC2, ST7" |
| 23 | SUPT5H | 8 | ADAMTS5, DOCK4, LTBP1, LTBP4, PCOLCE2, RAP1B, ST7, VIT |
| 24 | XRN2 | 8 | APBB2, DOCK4, LTBP1, LTBP4, NID1, PCOLCE2, RAP1B, ST7 |
| 25 | CTNNB1 | 7 | DDR2, DPP4, EMILIN2, NID1, PXN, RAP1A, ST7 |
| 26 | LIN28B | 7 | APBB2, DPP4, NID1, PCOLCE2, PXN, RAP1A, SDC2 |

| | | | |
|---|---|---|---|
| 27 | NEUROG2 | 7 | ADAMTS5, DDR2, DPP4, FLRT2, ITGA11, RAP1B, VIT |
| 28 | POU2F1 | 7 | APBB2, DPP4, EMILIN2, FBN1, RAP1A, ST7, TNXB |
| 29 | ZSCAN22 | 7 | ADAMTS5, APBB2, EMILIN2, FBN1, NID1, RAP1B, ST7 |
| 30 | ZC3H7B | 6 | APBB2, DOCK4, PXN, RAP1A, SDC2, ST7 |
| 31 | CBX8 | 5 | DCN, DPP4, EMILIN2, PCOLCE2, RECK |
| 32 | GSPT2 | 5 | DOCK4, LTBP1, PCOLCE2, PXN, RAP1B |
| 33 | LIN28A | 5 | APBB2, NID1, PCOLCE2, PXN, SDC2 |
| 34 | TTF2 | 5 | APBB2, DOCK4, ITGA11, NID1, RAP1B |
| 35 | DROSHA | 3 | FBLN2, ITGA11, TNXB |
| 36 | LARP7 | 3 | NID1, RAP1B, RECK |
| 37 | PAX8 | 3 | FLRT2, PXN, RAP1B |
| 38 | MED26 | 2 | FLRT2, RAP1B |
| 39 | ZNF224 | 1 | ITGA11 |

**Table S7: Expression of enriched transcription factors putatively targeting differentially-expressed ECM genes of fibroblast subcluster 0.** Differential expression of each transcription factor was determined between fibroblast subcluster 0 versus subcluster 1, considering only transcription factors that targeted > 15 ECM genes in fibroblast subcluster 0.

| | p_val | avg_logFC | pct.1 | pct.2 | p_val_adj |
|---|---|---|---|---|---|
| PGR | 1.41E-74 | 1.3636 | 0.216 | 0.039 | 4.22E-70 |
| BHLHE40 | 4.46E-39 | 0.79558 | 0.228 | 0.093 | 1.33E-34 |
| ARNT | 1.57E-21 | -0.5253 | 0.507 | 0.54 | 4.70E-17 |
| AHR | 1.47E-19 | -0.5892 | 0.336 | 0.403 | 4.38E-15 |
| ELF1 | 2.96E-18 | 0.12476 | 0.781 | 0.591 | 8.83E-14 |
| CDK8 | 2.53E-09 | -0.5375 | 0.327 | 0.364 | 7.54E-05 |
| RCOR1 | 6.57E-05 | -0.4922 | 0.374 | 0.373 | 1 |
| GABPA | 0.00022 | -0.152 | 0.211 | 0.16 | 1 |
| SMC1A | 0.00375 | -0.4821 | 0.196 | 0.214 | 1 |
| USF2 | 0.08544 | -0.3238 | 0.106 | 0.089 | 1 |
| SMAD1 | 0.18984 | -0.2968 | 0.143 | 0.151 | 1 |

**Table S8: Differentially-expressed genes in advanced-stage fibroblasts upon BHLHE40 knockdown in vitro.** Nanostring analysis performed on RNA extracted from fibroblasts with or without *BHLHE40* knockdown siRNA in vitro (KLIII/IV; n=5). *BHLHE40* knockdown resulted in 42 upregulated and 5 downregulated genes, compared to control siRNA treatment. Differences between *BHLHE40* siRNA treatment and control siRNA treatment were assessed using a generalized least-squares regression model (paired analysis). Genes are organized by decreasing logFoldChange. P-values were adjusted for multiple comparisons using the Benjamini-Hochberg correction, with adjusted p<0.05 considered significant. Full list of Nanostring Fibrosis V2 Panel genes available upon request.

| Gene | type of test | Comparison | Up or Downregulated | logFoldChange | p value | p value adjusted |
|---|---|---|---|---|---|---|
| LOXL4 | paired | BHLHE40 v Ctrl | Up | 0.398 | 0 | 0.005 |
| LEP | paired | BHLHE40 v Ctrl | Up | 0.397 | 0.001 | 0.026 |
| BANF1 | paired | BHLHE40 v Ctrl | Up | 0.353 | 0.001 | 0.026 |
| FABP4 | paired | BHLHE40 v Ctrl | Up | 0.32 | 0.001 | 0.018 |
| CTSL | paired | BHLHE40 v Ctrl | Up | 0.283 | 0 | 0.006 |
| PGM1 | paired | BHLHE40 v Ctrl | Up | 0.243 | 0.001 | 0.019 |
| CDKN1A | paired | BHLHE40 v Ctrl | Up | 0.207 | 0.001 | 0.028 |
| NUBP1 | paired | BHLHE40 v Ctrl | Up | 0.206 | 0 | 0.007 |
| ITGA5 | paired | BHLHE40 v Ctrl | Up | 0.202 | 0.003 | 0.046 |
| CRK | paired | BHLHE40 v Ctrl | Up | 0.172 | 0.001 | 0.025 |
| AP1S1 | paired | BHLHE40 v Ctrl | Up | 0.164 | 0 | 0.007 |
| ITGBL1 | paired | BHLHE40 v Ctrl | Up | 0.159 | 0.002 | 0.037 |
| ADIPOR1 | paired | BHLHE40 v Ctrl | Up | 0.151 | 0 | 0.011 |
| FZD5 | paired | BHLHE40 v Ctrl | Up | 0.142 | 0.001 | 0.019 |
| SSR2 | paired | BHLHE40 v Ctrl | Up | 0.137 | 0 | 0.003 |
| HIF1A | paired | BHLHE40 v Ctrl | Up | 0.135 | 0.002 | 0.038 |
| MMP14 | paired | BHLHE40 v Ctrl | Up | 0.124 | 0.002 | 0.031 |
| YWHAE | paired | BHLHE40 v Ctrl | Up | 0.123 | 0 | 0.005 |
| IRS1 | paired | BHLHE40 v Ctrl | Up | 0.122 | 0 | 0.011 |
| COL4A1 | paired | BHLHE40 v Ctrl | Up | 0.121 | 0 | 0.012 |
| PTK2 | paired | BHLHE40 v Ctrl | Up | 0.12 | 0.002 | 0.036 |
| PSMB7 | paired | BHLHE40 v Ctrl | Up | 0.12 | 0 | 0.005 |
| CSNK1D | paired | BHLHE40 v Ctrl | Up | 0.117 | 0.001 | 0.018 |
| NDUFA1 | paired | BHLHE40 v Ctrl | Up | 0.112 | 0.001 | 0.026 |
| BCL2L1 | paired | BHLHE40 v Ctrl | Up | 0.109 | 0 | 0.011 |
| SEC24C | paired | BHLHE40 v Ctrl | Up | 0.107 | 0 | 0 |
| NOL7 | paired | BHLHE40 v Ctrl | Up | 0.105 | 0 | 0.004 |
| PRDX1 | paired | BHLHE40 v Ctrl | Up | 0.103 | 0.001 | 0.026 |
| PSMB1 | paired | BHLHE40 v Ctrl | Up | 0.101 | 0.002 | 0.031 |
| PIK3C3 | paired | BHLHE40 v Ctrl | Up | 0.087 | 0 | 0.006 |
| NDUFC1 | paired | BHLHE40 v Ctrl | Up | 0.08 | 0 | 0.005 |
| TNC | paired | BHLHE40 v Ctrl | Up | 0.077 | 0.001 | 0.019 |
| BCAP31 | paired | BHLHE40 v Ctrl | Up | 0.077 | 0 | 0.011 |
| YWHAG | paired | BHLHE40 v Ctrl | Up | 0.071 | 0 | 0.01 |
| POS_D | paired | BHLHE40 v Ctrl | Up | 0.07 | 0.001 | 0.028 |
| YWHAQ | paired | BHLHE40 v Ctrl | Up | 0.07 | 0.001 | 0.029 |
| RBX1 | paired | BHLHE40 v Ctrl | Up | 0.066 | 0.001 | 0.019 |
| COX6B1 | paired | BHLHE40 v Ctrl | Up | 0.066 | 0.001 | 0.026 |
| NEDD8 | paired | BHLHE40 v Ctrl | Up | 0.058 | 0 | 0.005 |
| CREB1 | paired | BHLHE40 v Ctrl | Up | 0.054 | 0.001 | 0.025 |
| HSP90AA1 | paired | BHLHE40 v Ctrl | Up | 0.038 | 0 | 0.004 |
| RAC1 | paired | BHLHE40 v Ctrl | Up | 0.031 | 0 | 0.002 |
| DDR2 | paired | BHLHE40 v Ctrl | Down | -0.079 | 0.001 | 0.022 |
| PRKDC | paired | BHLHE40 v Ctrl | Down | -0.122 | 0 | 0.004 |
| HDAC4 | paired | BHLHE40 v Ctrl | Down | -0.135 | 0 | 0.004 |
| EP400 | paired | BHLHE40 v Ctrl | Down | -0.214 | 0.002 | 0.038 |
| GNLY | paired | BHLHE40 v Ctrl | Down | -0.254 | 0.001 | 0.026 |
